# FriendlyClearMap: An optimized toolkit for mouse brain mapping and analysis

**DOI:** 10.1101/2023.02.16.528882

**Authors:** Moritz Negwer, Bram Bosch, Maren Bormann, Rick Hesen, Lukas Lütje, Lynn Aarts, Carleen Rossing, Nael Nadif Kasri, Dirk Schubert

**Author notes:** shared senior (Nael Nadif Kasri, Dirk Schubert).

## Abstract

Tissue clearing is currently revolutionizing neuroanatomy by enabling organ-level imaging with cellular resolution. However, currently available tools for data analysis require a significant time investment for training and adaptation to each laboratory’s use case, which limits productivity. Here, we present FriendlyClearMap, an integrated toolset that makes ClearMap1 and ClearMap2’s CellMap pipeline easier to use, extends its functions, and provides Docker Images from which it can be run with minimal time investment. We also provide detailed tutorials for each step of the pipeline.

For more precise alignment, we add a landmark-based atlas registration to ClearMap’s functions as well as include young mouse reference atlases for developmental studies. We provide alternative cell segmentation method besides ClearMap’s threshold-based approach: Ilastik’s Pixel Classification, importing segmentations from commercial image analysis packages and even manual annotations. Finally, we integrate BrainRender, a recently released visualization tool for advanced 3D visualization of the annotated cells.

As a proof-of-principle, we use FriendlyClearMap to quantify the distribution of the three main GABAergic interneuron subclasses (Parvalbumin^+^, Somatostatin^+^, and VIP^+^) in the mouse fore- and midbrain. For PV^+^ neurons, we provide an additional dataset with adolescent vs. adult PV^+^ neuron density, showcasing the use for developmental studies. When combined with the analysis pipeline outlined above, our toolkit improves on the state-of-the-art packages by extending their function and making them easier to deploy at scale.

## Introduction

For most of its century-long history, histology has been the 2-dimensional study of 3-dimensional tissues. This was mostly due to technical limitations, specifically 2-dimensional fields of view, in combination with most tissues being too opaque to image at large scale and high resolution for deeper than tens of μm. Even though tissue clearing by refractive index matching was invented more than a century ago ^1^, a lack of imaging and analysis capabilities limited our ability of acquiring high resolution images and quantifying data obtained from transparent tissues. In the last decade, the twin innovations of light-sheet microscopy and improvements in brain clearing techniques have enabled 3-dimensional imaging of centimeter-sized organs with subcellular resolution^2^.

However, 3D imaging data produces complex, multi-gigabyte image stacks that cannot easily be processed manually. This requires the development of specialized image analysis pipelines optimized for specific analysis tasks, such as identifying the features of interest, mapping them to a reference template and visualizing the results in 3D ^3–6^. Unfortunately, those software packages tend to rely on a complex and fragile environment of supporting software, e.g. Python packages in specific versions. As a result, many of those software pipelines are brittle and require frequent manual updates by the user, adding complexity and harming reproducibility.

Here we present FriendlyClearMap, a Docker container hosting adapted versions of ClearMap1 ^4^ and ClearMap2’s CellMap pipeline ^5^. Due to containerization, the entire environment is self-contained and comes with all required dependencies already included, removing the need to maintain a complex Python package environment by hand. The containerized tool can be deployed across hardware and software platforms, as well as cloud environments – we verified function on Microsoft Windows 10 and Ubuntu Linux 20.10 - 22.04, on laptop- and workstation-grade hardware, as well as in an Amazon AWS cloud instance and an Apple Mac Pro.

As a proof-of-principle, we furthermore demonstrate the processing, analyses and visualization routines on three datasets revealing the distribution of the three main classes of GABAergic neurons in the developing wild-type mouse brain: Parvalbumin-positive (PV^+^), Somatostatin-positive (SST^+^) and Vasoactive Intestinal Peptide-positive (VIP^+^) neurons. Together, these cell types account for approximately 90% of GABAergic neurons in the brain ^7–10^. Our data extends on previous reports on the distribution patterns of PV/SST/VIP expressing in the mouse ^11,12,13^ by adding a comprehensive whole-brain PV^+^ neuron quantification across the developing brain, as well as cross-modal validation for all three neuron sets. Since PV^+^ neurons mature during adolescence in the mouse ^14–16^, in contrast to SST^+^ and VIP^+^ neurons^17^, we analyzed PV^+^ neuron density at both adolescence (postnatal day 14 (P14)) and in adulthood at P56. We demonstrate that PV expression in OFC areas seem to mature earlier than surrounding rostral regions such as the mPFC, in which PV^+^ cell density ramps up between P25 and P45 (for review see ^18^). This has relevant implications when considering OFC’s specific deep-brain connectivity ^19^ and its role in prediction error encoding ^20^.

### Findings

Here we present an expanded and containerized version of ClearMap1 ^4^ and the CellMap part of ClearMap2 ^5^. Besides enabling easier deployment via Docker, we have implemented the following features:

- Atlas alignment via landmarks
- Updated atlases for juvenile mice (adapted from ^21^)
- Segment cells with Ilastik
- Import Segmented cells, e.g. from commercial programs or manual annotations
- Check alignment quality
- Easy follow-up visualization with Brainrender ^22^
- Exploring cortical cell distribution in detail with a cortical flatmap

### Interface: Packaging ClearMap in a docker container for accessibility and added flexibility

The use-case of our pipeline is workstation-based processing of image stacks of cleared mouse brains. The problem we aimed for resolving is that published pipelines, though well-documented, require familiarity with Python environments which are present in a dynamically changing landscape of python packages, typically running on top of a Linux operating system. Even though knowledge of those areas is getting more common in (neuro)biology, it is still uncommon and likely to present a significant obstacle to a new brain-clearing biologist. Navigating those computational environments can be a challenging task for the average light-sheet microscope user, especially if it also involves ongoing maintenance of a Python package environment, typically without a local support infrastructure e.g. by researchers in the field of (bio)informatics.

In order to simplify deployment, we have packaged a version of ClearMap1 (rewritten in Python 3.8) and the CellMap portion of ClearMap2 in Docker containers. We provide the Docker containers to create custom containers as well, see Appendix 1. The Docker containers are set up to execute a version of ClearMap’s process_template script upon start, write the results to the same folder, and then exit. All relevant data is retained in the folder. Please see Figure 1 for an overview of the workflow and options of this pipeline, and Appendix 2 for detailed instruction how to set up a Docker environment, prepare the image stacks, segment the cells, and then run the FriendlyClearMap docker containers.

**Figure 1:**
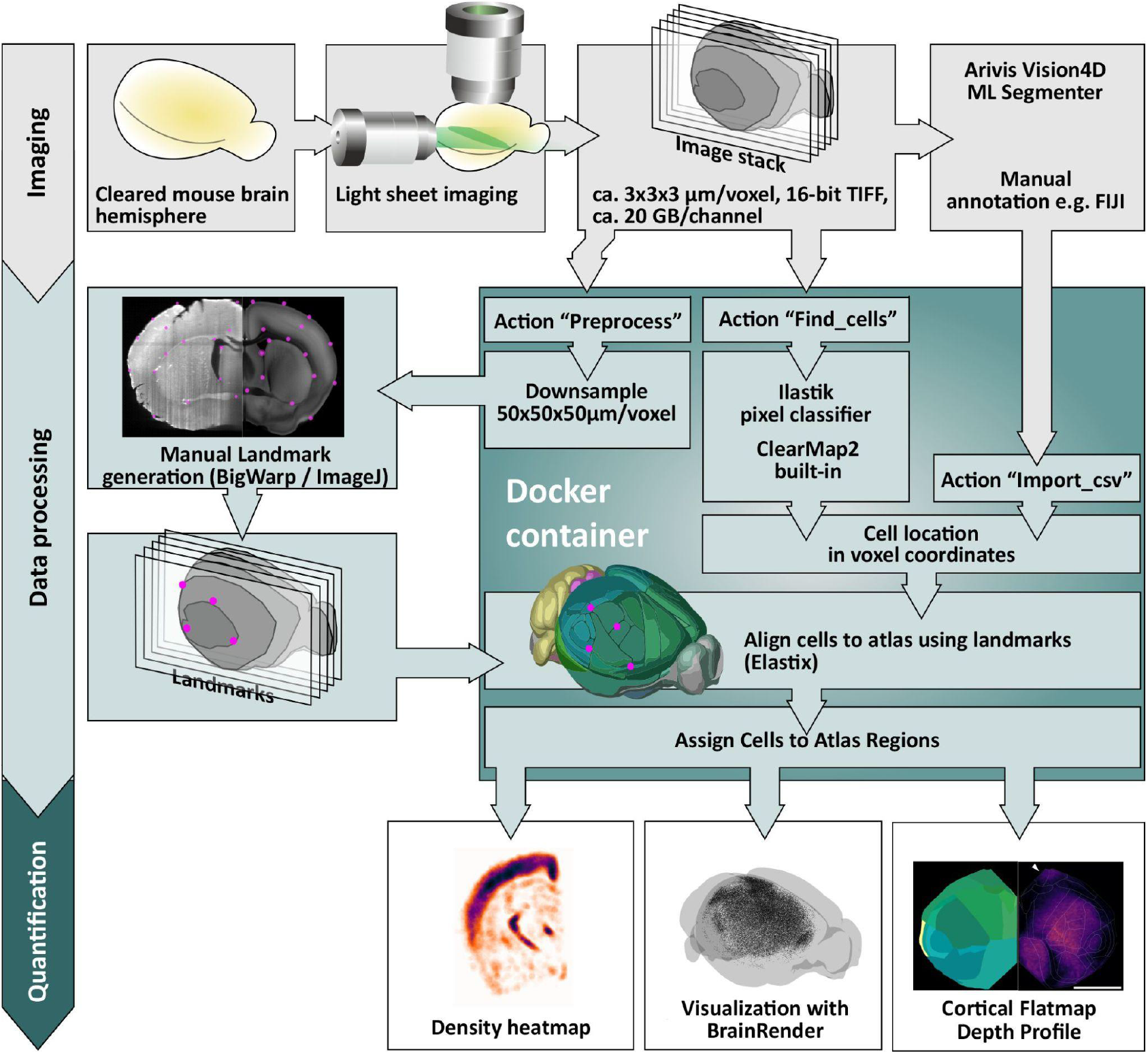
FriendlyClearMap’s processing steps illustrated. Top (gray): Data generation. Mouse brain hemispheres are stained and cleared with iDISCO+, and imaged with a light-sheet microscope. The coordinates of labelled cells can be identified with an external program or marked manually, and subsequently imported into the pipeline. Middle (light blue): Dockerized pipeline steps. Middle left: Landmark-based atlas alignment with BigWarp. Use the preprocessing function (inside the Docker) to generate a downsampled image stack. Subsequently, use the ImageJ plugin BigWarp to mark corresponding landmarks on both downsampled stack and on the atlas. Middle center: Use of ClearMap1 &2’s built-in cell-finding pipeline for threshold-based segmentation, or alternatively use an Ilastik Pixel Classifier workflow to identify cells with Machine Learning. (All of those steps are described in Supplementary protocol 1 and 2. In each case, the landmarks are then used to transform the pixel coordinates from image-stack space to atlas space and assigned to atlas regions. Bottom (dark blue): Quantification steps outside of the Docker container. Left: Density heatmap as generated by ClearMap1/2. Middle: We also provide optimized scripts for visualizing the output with BrainRender, described in detail in supplementary protocol 3. Right: Voxel-level p-value maps and region quantifications as generated by ClearMap1/2.

### Enhancing Atlas Alignment with Landmarks

The ClearMap papers describe that the reference atlases need to be carefully cropped to the same dimensions as the image stacks, which in our own experience was often a time-intensive and produced subpar results (data not shown). Instead, we integrated a graphical landmark-based atlas registration method with the BigWarp Fiji plugin, which allows for a direct comparison of both image and atlas volumes in 3D during landmark generation. The landmarks generated there are then used as input for Elastix’ landmark feature, now part of the “preprocess” workflow (Figure 1). With this semi-automated process, the user generates matching landmark points between the image stack and the reference atlas, which are then exported as csv and used to feed Elastix’ *fixed points / moving points* option. The points are generated with the FIJI plugin BigWarp ^23^, where the stacks can be easily rotated and moved with an intuitive graphical interface. Importantly, BigWarp generates a landmarks file, which can be re-loaded. After re-loading, individual landmarks can be adapted, which allows for an iterative fine-tuning of the alignment. In any case, the graphical interface makes the alignment procedure more accessible to the non-computational neuroscientist.

For the datasets presented here, we used landmark-based alignment exclusively. The default Elastix alignment strategy used in ClearMap (Affine, then b-Spline) tends to work best when the atlas is tailored to match the image stack. However, this requires precise cropping of the atlas, which can be a time-consuming iterative process, in particular if the sample is not cut straight within one of the XYZ planes or has warped during iDISCO+ treatment. Please see Appendix 2 for an in-detail description of the process.

### Additional atlases for developing mouse brains

By default, ClearMap1 and 2 use Elastix ^24,25^ to automatically align a down-sampled image stack (typically of the autofluorescence channel) to the Allen Brain Atlas Common Coordinate Framework (CCF) mouse brain reference template. ClearMap1 uses the reference STPT atlas from ^13^, whereas ClearMap2 uses the newer Allen Brain Atlas CCF3 ^26^. Both templates were generated for adult mice at the age of postnatal day P56. The Allen Institute has generated 2-dimensional slice-based atlases for developing mouse brains down to the embryo stages, however those only have coarse regions and are not available as 3-dimensionally registered image stacks. However, a recent publication independently re-mapped the CCF3 adult regions on their own reference atlases for mouse brains at ages of P7, P14, P21, P28 and P35 to generate high-resolution developmental maps ^21^. We integrate those atlases in our ClearMap2 container as an option, thus enabling a well-controlled mapping of even very young mouse brains. As a proof of concept, we use the juvenile reference mouse brain at P14 in order to map PV^+^ neurons (Fig 2a-f).

**Figure 2:**
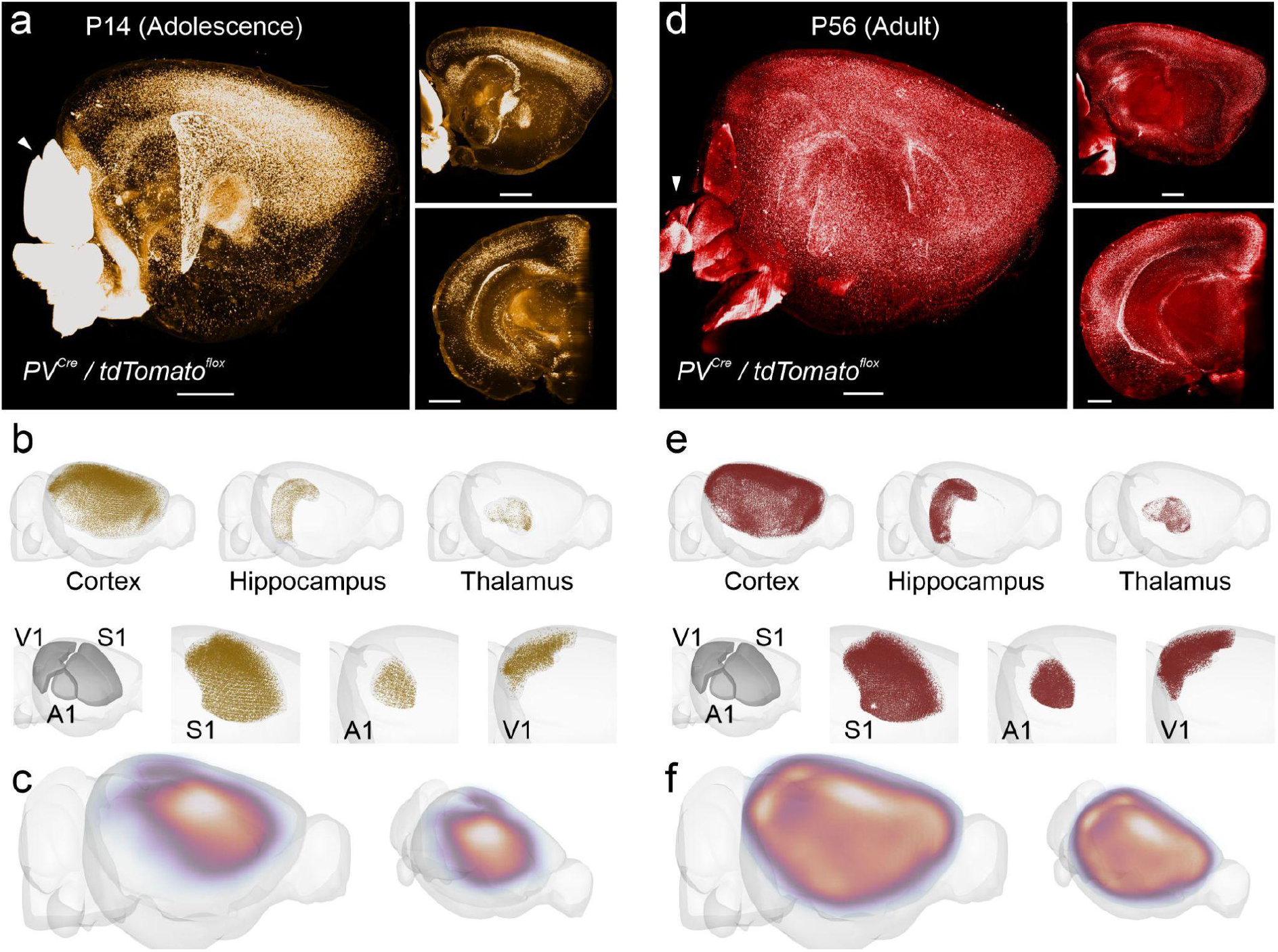
Quantitative assessment of changes in parvalbumin expressing neuron distribution during development. We used our pipeline to identify PV^+^ neurons expressing the TdTomato reporter throughout the adolescent (P14) vs adult (P56) brain. PV^+^ neurons start expressing PV as part of their maturation, driven by sensory inputs. **a), d):** Sagittal views of representative image stacks for P14 (a) and P56 (d). Note that the cerebellum (arrowhead) was so bright for TdTomato due to the local abundance of the PV^+^ Purkinje cell dendrites that it saturated our detector; consequently we only segmented cells outside of the cerebellum for P14 and cut it off for P56. **b), e):** PV^+^ neuron density per brain area, after processing with our FriendlyClearMap pipeline and visualization with BrainRender. PV^+^ neuron density in the P14 cortex (b) is higher in sensory areas than in extrasensory areas, and lower in V1 than in S1. Density is much higher in all of those areas at P56 (e). **c), f):** Relative PV^+^ neuron density in the cortex at P14 (c) and P56 (f). At P14, PV^+^ neuron density is high in the somatosensory and auditory areas, which already receive sensory input at this time. In contrast, the visual cortex only starts receiving input around P12-14, consequently PV^+^ expression is low. At P56 in contrast, PV^+^ expression is high across the cortex, with especially dense clusters in S1 and V1. Note that the heatmaps are only scaled relative to each dataset, therefore the densities are not directly comparable between the two timepoints.

Both CCF2 and CCF3 reference templates were generated by averaging hundreds of mouse brain autofluorescence stacks generated by serial two-photon tomography (STPT). The brain processing for the reference templates notably differs from the solvent-based clearing used here - specifically, the aggressive permeabilization and solvent-based delipidation used in iDISCO+ clearing cause region-specific changes in autofluorescence signal and anisotropic tissue shrinkage, especially in the lipid-rich fiber tracts. This tissue shrinkage complicates the registration to the reference atlas, which can be solved by a clearing-optimized variant of the CCF3 ^27^. This optimized atlas can indeed be used in our ClearMap2 container as drop-in replacement for the default CCF3 standard template.

### Cell Segmentation with Machine Learning

The original ClearMap pipeline relies on simple intensity-based image transformations such as thresholding and watershed segmentation to detect cells ^4^. This approach is designed for fast computation, however, it requires extensive fine-tuning and is strongly dependent on the local background intensity in the tissue. Even in the original ClearMap publication, the conservative size threshold chosen for cell detection using the expression of the immediate early gene cFos meant that approximately 75% of potential cells get discarded compared to a lower-confidence “human error threshold” that averages similar numbers as human raters (see Renier et al., 2016; Supplementary Fig 5C). Furthermore, we found that local variations in background intensity in our samples (e.g. between cortex and thalamus) distorted cell counts in a region-specific way when using the original ClearMap intensity-based processing (data not shown). In order to remedy this, we implemented three approaches: First, we set up an import function for externally annotated cell coordinates that can be imported as .csv table containing the center point XYZ coordinates in the original image stack space. Those coordinates can be hand-annotated, for example when cells are very sparsely distributed.

Secondly, the cell segmentations can also come from commercial image analysis packages, for example Arivis Vision4D’s Machine Learning Segmenter. Those packages can be useful for the end user because they abstract away many details about fine-tuning detection algorithms and can be set up with relative ease. They may have been part of pre-established analysis pipelines and thus benefit from previous work and validation. All comparable packages (e.g. Imaris or Aivia) can export coordinates as .csv, and thus can be integrated into this pipeline without additional setup. We detail the export procedure for Arivis Vision4D’s Machine Learning Segmenter in Appendix 2.

Finally, we closely integrated the machine-learning segmentation offered by the open-source program Ilastik ^28^ into the analysis pipeline. ClearMap1 and 2 already contained code paths for using Ilastik, however this integration has been broken by package incompatibilities since the original publications. We corrected those dependencies, ported the Ilastik portion over to ClearMap2’s CellMap package (by default Ilastik is only used by the TubeMap package), and package a recent version (4.1.8-beta) with both ClearMap1 and 2 in their respective Docker containers. Of note, since Ilastik is open-source, it can run inside the Docker container, which enables parallel processing of several datasets on independent instances, e.g. on inexpensive rented cloud computers. The multi-threaded, CPU-heavy Random Forest Algorithm of Ilastik’s Pixel Classifier workflow scales well to approx. 24 CPU cores and can be run on anything from laptops to desktop PCs to cloud instances and shared cluster environments. Ilastik has a graphical interface to generate training data for the Random Forest Algorithm, which makes training a relatively straightforward task. We describe the setup procedure for a custom Ilastik workflow in Appendix 2.

### Visualization with BrainRender

ClearMap1/2 produces heatmaps for cell density and p-values on the voxel level for quantitative analyses. We preserved this functionality in the Docker containers. However, we decided to include additional visualization tools such as BrainRender ^22^. BrainRender is a recently published visualization pipeline from the Branco lab that allows to semi-automatically plot cells that are mapped onto the ABA CCF3, with a reasonably easy python scripting interface. Here, we also provide pre-made transformations to transform data from the custom P7-P28 reference brains ^21^ and the CCF2 (for ClearMap1 data) on the CCF3 for BrainRender. BrainRender requires access to 3D rendering architecture and is therefore incompatible with running in a Docker container. Therefore, we recommend executing it within a suitable Python development environment such as Spyder (Scientific PYthon EnviRonment, https://www.spyder-ide.org/).

Furthermore, we also provide a script to compare two experimental groups of animals by creating a region-based heat map. This is an extension of ClearMap’s voxel-based heatmap, but it allows for easier comparison between different reference spaces, such as adolescent *versus* adult brains. In our proof-of-principle analyses, i.e. when comparing PV^+^ cell counts from juvenile (P14) *versus* adult (P56) mouse brain maps we demonstrated the feasibility of this approach (Figure 2g). To account for size differences, we also include a script to quantify the size (in voxels) of each reference atlas, allowing a direct neuron density comparison. For detailed usage instructions, please see Appendix 2.

### Quantitative assessments: Cortical depth profiles and Flat-map projections (Fig 4)

Cleared whole-brain imaging offers an additional advantage over sectioning approaches: it allows computing quantitative cortical depth maps across the entire cortex. Previously, the distribution of cells could be assessed ex vivo either in the cortical depth of a limited set of incomplete brain regions (via coronal sections, e.g. ^29^) or over the cortical sheet in a flattened cortex preparation, where the cortex is physically peeled off the rest of the brain and flattened between glass plates prior to sectioning (e.g. ^30^). With the correct mapping, an atlas-aligned cortex can be projected into a flat 3D sheet in silico, which enables both the study of cell distribution across the cortical sheet and the depth.

For this, we expanded on the work released by the Allen Brain Institute as part of their CCF3 documentation^26^ (https://ccf-streamlines.readthedocs.io/en/latest/data_files.html#data-files), integrating it into our pipeline and adding a heatmap generation function. Our flatmap script takes the CCF3-referenced cell coordinates and generates a flattened-cortex (“Butterfly”) projection. This projection preserves the cortical depth (via the “streamlines” approach documented here: https://community.brain-map.org/t/ccfv3-highlights-tilting-at-the-cortex/1000/4). As a consequence of this in silico projection the cortical surface is not isotropic anymore. This means that central areas such as the primary somatosensory cortex are rendered smaller than they would be in a curved cortex, while peripheral areas such as the temporal association area is rendered larger. However, this in silico projection does allow a direct comparison of the distribution of different cell types across the cortical sheet. Furthermore, it enables straightforward plotting of depth profiles, both for the entire cortex and specific sub-areas, which is relevant for assessing layer-specific effects^16^ and more fine-grained phenotypes such as gradients within layers ^31^.

## Reuse Potential

### Pipeline Reuse Potential

FriendlyClearMap offers an improvement over the state-of-the-art cleared mouse brain analysis pipelines, as it is specifically geared to the average neurobiologist who does typically neither have the required technical knowledge to maintain a Linux machine, much less to maintain a Python environment nor extensive experience in script coding. With FriendlyClearMap, we address this by packaging the core pipeline in a docker container that can also be run on Windows PCs, which are far more common in a typical biology laboratory. The versatility of the docker platform also includes executability on different operating systems or environments, such as MacOS and in cloud-based environments. This allows users to run FriendlyClearmap without extensive computation capabilities of their own.

The current state-of-the-art pipelines also require parameter optimalization for e.g. atlas alignment and cell detection, which requires experience with Elastix and image analysis packages. This presents a steep learning trajectory for the average neurobiologist, who might be more used to judge the quality of microscope images. We address this issue by two additions that enable interactive atlas registration (using BigWarp for landmark generation) and custom-tailored cell classifier data sets (using Ilastik). In both cases, the interaction is directly with the image stacks instead of parameter files, which allows for easier, visual, fine-tuning without having to learn the underlying code’s functions first. Thus, we believe that our additions will make the pipeline very accessible and therefore useful to a larger neurobiology community.

Scientifically, the current state-of-the-art pipelines require the use of an adult mouse brain atlas (see ^4,5,27^), which poses problems for the increasing number of developmental studies. We addressed this by the inclusion of additional juvenile-brain atlases ^21^ that enable developmental studies starting from as young as postnatal day 7. Of note, juvenile mouse brains are exceptionally well suited for whole-brain clearing and fluorescent labelling, which makes this pipeline directly relevant for a previously underserved part of the developmental neuroscience community.

Lastly, the current state-of-the-art pipelines encompass limited data analysis capabilities, generating tables and heatmaps, and relying on external tools for further visualization. We address this issue by including BrainRender not only for advanced visualization (notably the ability to generate region-specific renderings of cell density), but also for in-depth quantitative cortical analyses via flatmap projection (notably cortical depth maps). This analysis will enable more in-depth analysis of upcoming as well as pre-existing datasets, especially with a focus on the cortex and the developmental trajectory and cellular compositions.

To conclude, we are convinced that our additions to the pipeline optimizeize, and easier to get high-quality visualizations than the current state of the art pipeline. it for a neurobiologist audience: We make it easier to install, easier to learn, easier to optim

### Dataset reuse potential

With the proof-of-principle datasets in this report, we present the first 3-dimensional whole-brain map of GABAergic neurons generated by using iDISCO+ whole-brain clearing in combination with light sheet microscopy. Furthermore, the adolescent PV^+^ dataset represent (to our knowledge) the most complete map of PV^+^ cells in the P14 mouse brain that is publically available. We foresee that the datasets will be useful to the neuroscience community as reference for future mouse studies focusing on cell type-specific (GABAergic) neuronal distribution patterns during development, or in mouse models for neurological disorders.

Furthermore, the background autofluorescence in our image set also contains useable information, for example on the position of the barrels in layer 4 of the primary somatosensory cortex, which represent relevant structural and functional processing modules within the somatosensory system. While all current atlases denote a barrel field, they do not specify individual barrels, and to date for visualization and quantitative assessment barrels had to be delineated manually ^32^. With the cortical flatmaps produced by our pipeline, delineating barrels could become sufficiently easy to be automated in future studies. Besides barrels, whole-brain autofluorescence can be used to delineate white matter tracts, as well as subcortical structures.

### FriendlyClearMap maps the spatial distribution of the three main GABAergic neuron types in the mouse brain across space and time

In order to highlight the utilitisation of our pipeline, we used it to investigate the three main GABAergic interneuron population’s 3-dimensional distribution throughout the mouse brain. Specifically, we investigated the distribution of PV^+^, SST^+^ and VIP^+^ neurons which comprise approximately 90% of GABAergic neurons in the brain ^7–10^. Being able to quantify numbers and distribution of different classes of inhibitory neurons is relevant for the study of neuropathological phenotypes, where dysfunction of GABAergic neuron classes is a common feature ^33^. In those studies, whole-brain datasets can help identify especially hard-hit brain regions and identify functional networks that are likely to be especially impaired^34^. In order to visualize the labelled neurons, we used reporter mice expressing tdTomato, which we then visualized with an anti-RFP antibody staining and whole-hemisphere iDISCO+ clearing. We scanned the transparent brains with a light-sheet microscope, and used our pipeline to analyze the resulting image stacks.

### Dataset 1: PV^+^ neuron density at Adolescence and Adulthood

We first investigated the largest group of GABAergic interneurons, i.e. neurons expressing the calcium-binding protein Parvalbumin. PV^+^ neurons are the most abundant cortical GABAergic subclass, accounting for ca. 40% of all GABAergic neurons ^9^ and comprising 10 genetically identifiable subpopulations ^10^. The canonical PV^+^ neuron is a fast-spiking basket or chandelier cell, driven by local circuit activity and delivering quick, intense and reliable inhibition to either neuron somata (basket cells, ^35,36^) or modulating activity at the axon initial segment (chandelier cells, ^37–40^, but see ^41^). PV^+^ neurons are thought to finely tune local activity to match input intensity, and synchronize a neuron population’s output, thus driving cortical oscillations in the gamma range ^42,43^.

PV^+^ neurons mature late in comparison to the other groups, driven by local circuit activity ^44–47^, long-range inputs ^15,43,48^, and environmental factors such as the trans-synaptic transcription factor Otx2 ^49–52^, for reviews see ^17,43,53,54^). We therefore chose an early postnatal timepoint (P14), which is shortly after the eyes open and when all three primary sensory cortices start receiving sensory input (Somatosensory cortex since shortly after birth, Auditory cortex after P10 and Visual cortex after P14^17^). We compare the PV^+^ neurons labelled via PV-Cre driven tdTomato expression at P14 (Fig 2a-c) to the situation at P56, i.e. adulthood (Fig 2d-f).

Our data show that at P14, the density of PV^+^ neurons in the cortex is highest in the primary somatosensory cortex (Fig 2c, Fig. 4b). The cortical depth profile (Fig 4c) shows that cell density is highest in the middle of the cortical depth, i.e. cell density at this age is by the somatosensory cortical layer 4. Both are consistent with this regions earlier development starting around P8 ^15,17,43^.

**Figure 3:**
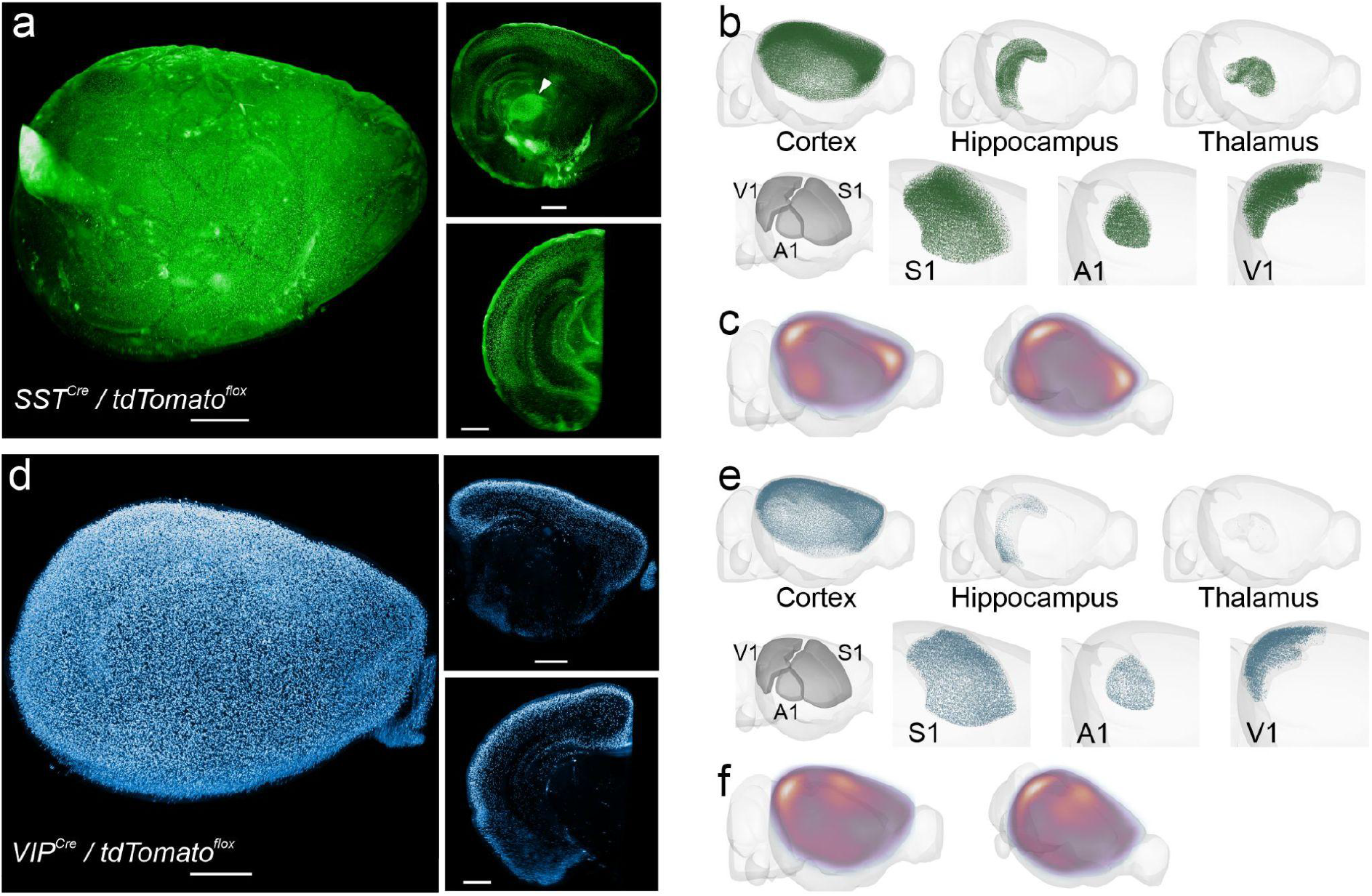
Somatostatin^+^ and VIP^+^ neurons are distributed in a cell-type specific pattern in the late-adolescent (P28) mouse brain. **a):** Example image stack of a SST^+^ reporter brain. The cortical SST^+^ neurons express TdTomato reporter in their axons, whose dense branches in layer 1 are visible as a bright layer even after iDISCO+ treatment. SST^+^ neurons are visible in the granular (layer 4) and infragranular (5-6) layers. Subcortically, they appear especially dense in the Globus Pallidus (arrowhead, top-right image). **b):** SST^+^ neuron distribution in the Isocortex, Hippocampus, Thalamus and a subset of sensory cortical regions, after processing with our FriendlyClearMap pipeline and visualization with BrainRender. **c):** Cortical SST^+^ neuron density showing a reduced density in the somatosensory and auditory regions at the center of the cortex, and an elevation towards the edges of the cortical plate. **d):** Example image stack of VIP^+^ reporter neurons in a mouse brain. VIP^+^ neurons are found mostly in the supragranular (layers 2-3) in the cortex and the hippocampus, but barely elsewhere in the brain. **e):** VIP^+^ neuron density in the Isocortex, Hippocampus, Thalamus and the three primary sensory cortical regions, after processing with our FriendlyClearMap pipeline and visualization with BrainRender. Note the high density of VIP^+^ neurons along the upper half of the cortex is caused by the overlap of both sensory cortical areas and the motor / retrosplenial cortex along the midline. Also note the complete absence of VIP^+^ neurons in the thalamus. **f):** Average VIP^+^ neuron density in the cortical plate, showing a relatively high density in the somatosensory areas and along the posterior and ventral edges of the cortex.

**Fig. 4:**
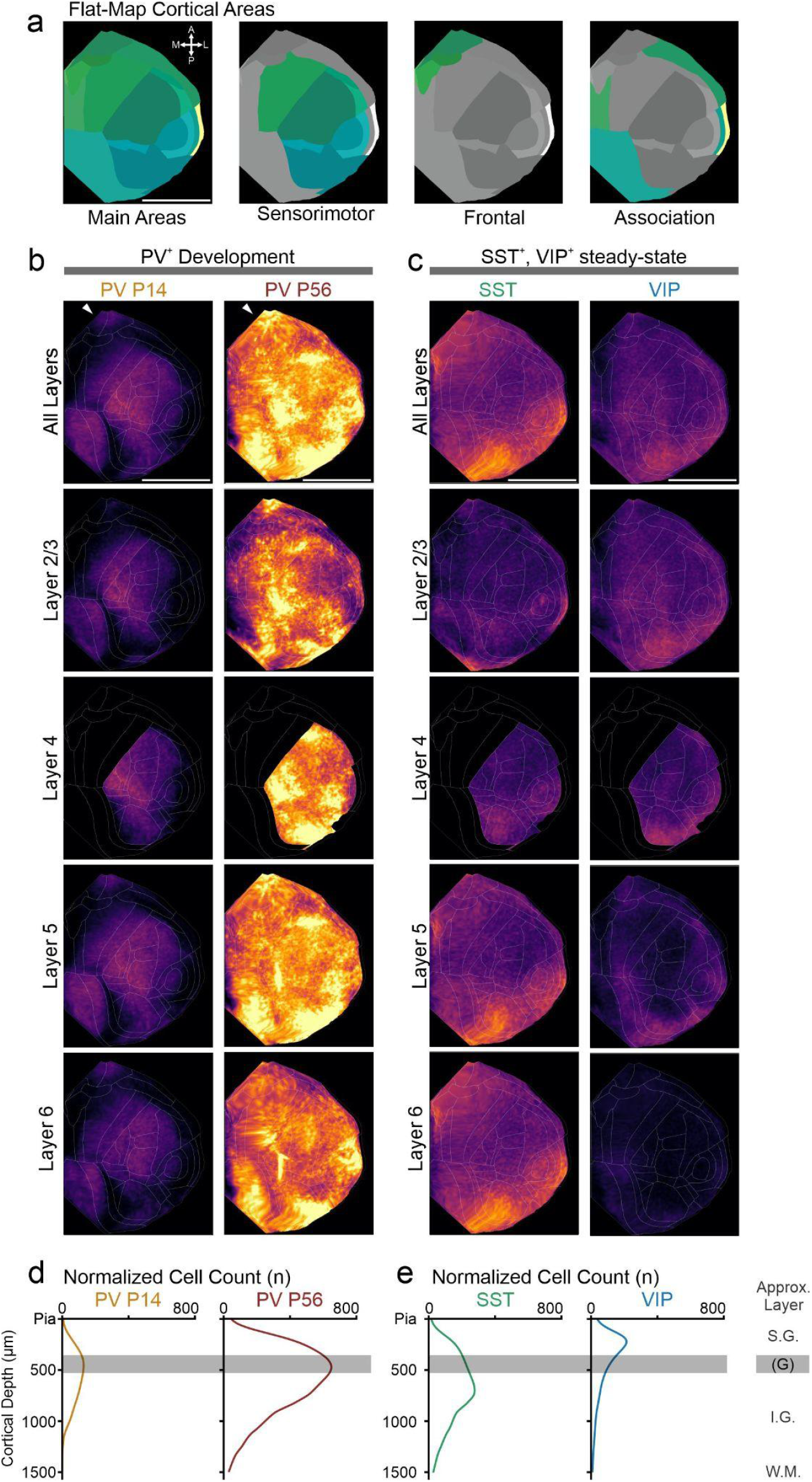
Cell distribution across cortical span and depth visualized by cortical flatmaps. **a)** Overview of the cortical areas that are visible in the flatmap (Butterfly) projection, color-coded as in the CCF3. Scale bar: 500 px (note that this projection does not preserve the spatial relations of the standard atlas space, so there is no direct μm equivalent except for depth). Orientation compass: A = Anterior, P = Posterior, M = Medial, L = Lateral. **b)** Flatmap projections of normalized cell density for PV^+^, SST^+^ and VIP^+^ cells, with cortical region boundaries outlined in white. The cell density is scaled to the same min/max values, enabling a direct comparison of densities across datasets. Top Row: All cortex, rows below: Computed cell densities for cortical layers 2/3 – 6. Note that cortical layer 4 (i.e. granular layer), which would be found approximately underneath the indicated “midline” is only present in granular cortices, such as sensory cortices. PV^+^ cell density at P14 (left column, n=7) is highest in the primary somatosensory cortex, with detectable expression in auditory, visual, retrosplenial, and interestingly orbitofrontal cortices (arrowhead, top row). At adulthood, PV^+^ cell density (right column, n=7) is much higher across the entire cortex, however relatively still highest in the sensory cortices, but again with interestingly high densities in retrosplenial and orbitofrontal cortices (arrowhead). SST^+^ cell density (column 3, n=3) is generally lower and mostly found at the frontal and posterior poles, respectively retrosplenial and OFC / PFC. **c)** In contrast to both PV^+^ and VIP^+^ cells, SST^+^ cells (left column, n=3) are overrepresented in the infragranular layers 5 and 6, consistent with a population largely consisting of Martinotti cells. Lastly, VIP^+^ cells (right column, n=7) are mostly found at the periphery of the cortical sheet, similar to SST^+^ cells. However, their density is lower and they are primarily found in the supragranular layer 2/3. **d)** Cortical depth plots enabled by the flatmap projection. The progression from PV^+^ P14 to adulthood is clear: Both have the highest density in the approximate location of layer 4, however there are substantially more PV^+^ cells in adult mice. **e)** In contrast, SST^+^ cells are mostly found in the infragranular layers 5-6, whereas VIP^+^ cells are mostly found in the supragranular layers 2/3. d) and e): Normalized cell density (n/μm depth), starting from the pial surface (mapped to CCF3 atlas space for comparability). As this projection preserves cortical thickness differences between different areas, we have only roughly indicated the position of the layers: S.G. = Supragranular layers 1, 2/3. G = Granular layer 4, if present. I.G. = Infragranular layers 5-6. W.M. = White Matter. b) - e): Averages of n=7 (PV P14), n=7 (PV P56), n=3 (SST) and n=7 (VIP) independent mice.

We also found some early maturation levels in the retrosplenial cortex, where early ethanol exposure at P7 has been reported to affect PV^+^ neurotransmission in adulthood ^55^ However, in addition we found that the orbitofrontal cortex did show a relatively high density at P14 as well, compared to e.g. visual cortex, which is an interesting deviation both temporally as well as spatially PV^+^ neurons are typically thought to mature first in the sensory areas, then in a general rostral-to-caudal gradient. However, PV expression in OFC areas seem to mature earlier than surrounding rostral regions such as the mPFC, in which PV^+^ cell density ramps up between P25 and P45 (for review see Klune et al., eLife 2021^18^). This developmental trajectory is relevant in the light of the OFC’s specific deep-brain connectivity ^19^, its role in prediction error encoding ^20^ and social interactions ^56^.. To the best of our knowledge, this early OFC PV^+^ maturation is a novel finding that has not been described before.

In the adult mouse brain at P56, we found PV^+^ neuron density to be uniformly higher than at P14, in line with a developmental phenotype (Fig 4b-c). Consistent with previous reports ^11^ we found that PV^+^ neuron density is highest in sensory areas (primary/secondary somatosensory, auditory and visual areas), and is generally reduced towards the edges of the cortical plate with notable clusters in orbitofrontal cortices and the ectorhinal cortex (Fig. 4b). Cortical depth profiles (Fig 4c) show that adult PV^+^ density is highest around the middle of the cortical plate, mostly driven by the granular layer 4 of the sensory cortices. Furthermore, the depth profiles show PV^+^density is much higher than in adolescence (indicating substantial maturation) and higher than both SST^+^ and VIP^+^ expression and in line with PV^+^neurons being the most abundant cell population^9^. In the subcortical areas, we found PV^+^ neuron density is especially high in the reticular nucleus of the thalamus (Fig 2e), again in line with previous publications ^11^. Unexpectedly, we found a small number of positive neurons scattered throughout the thalamus besides the high density found in the reticular nucleus.

To conclude, our dataset confirms that PV^+^ neurons are still early in their maturation at P14, with unexpected early maturation in the orbitofrontal and retrosplenial cortices in addition to the expected sensory cortices. In adulthood, cortical PV^+^ cell density is much higher, with the highest peaks in the sensory areas layer 4, but also substantially high densities in the orbitofrontal, retrosplenial and entorhinal cortices.

### Dataset 2: SST^+^ neuron density at P28

In contrast to PV^+^ neurons, the neurons from the second-largest subpopulation (the neuropeptide Somatostatin) start expressing their marker in the first postnatal week ^57,58^. However, the neurons keep maturing functionally until late adolescence and play an important role in the targeting and maturation of other cortical network components, such as thalamocortical projections and PV^+^ neurons ^58^. While the SST^+^distribution in adulthood is known ^11^, we here decided to investigate the SST^+^ distribution in adolescence (P28). We used SST-Cre to drive tdTomato expression in neurons expressing the neuropeptide Somatostatin (SST^+^, also known as SOM^+^; see ^8,9^). SST^+^ neurons account for approximately 30% of cortical GABAergic neurons ^9^ and contain 18 genetically different subgroups ^10^, with the most prominent cortical subpopulation being Martinotti cells ^59–62^. Martinotti cells are located in the infragranular cortical layers 5-6 and extend axon collaterals vertically up to layer 1, capable of inhibiting input to distal apical dendrites of lower-layer pyramidal cells. SST^+^ neurons are thought to play a role in blocking input from entire dendrite branches via shunting inhibition, thus providing a selective filter for long-range input which arrives at distal dendritic segments ^63,64^. Their axons branch in layer 1, which can be seen in Fig. 3a overview pictures as a bright, uniform layer on the outer cortical surface. Subcortically, SST^+^ neurons are also present in the hippocampus (e.g. Stratum Oriens-Lacunosum Moleculare (OLM) neurons, see ^65–67^), in the amygdala, and the inferior colliculus ^11^.

In the cortex, we indeed found that the majority of SST^+^ cells are located in the infragranular layers 5 and 6 (Fig 4b, c). SST^+^ cells were denser in the primary sensory areas, notably the S1 barrel field (Fig. 3b), pointing to a specific filtering role in the sensory processing of sensory information ^68,69^. Outside the primary sensory areas, SST^+^ neurons are present in a region-based gradient, with higher densities towards the edge of the cortical plate (Fig. 4b, consistent with ^11^). In the hippocampus, SST^+^ neurons were denser in CA1-3 compared to the Dentate Gyrus, in line with the somatic location of hippocampal SST^+^O-LM neurons. We also found dense SST^+^ neurons in the ventral portion of the striatum, as well as the Globus Pallidus. To conclude, with this dataset we provide the first whole-adolescent brain map of SST^+^ distribution. This dataset will be useful in future studies of SST^+^ development, and as a comparison to SST^+^distribution in neurodevelopmental disorder models with a suspected impact on SST^+^neurons.

### Dataset 3: VIP^+^ neuron density at P28

The third main type of GABAergic neurons we investigated were those expressing the neuropeptide Vasoactive Intestinal Peptide (VIP^+^). This category accounts for approx. 10% of cortical GABAergic neurons ^9^ and comprises 14 genetically identifiable subpopulations ^10^. The canonical VIP^+^ neuron fine-tunes local circuit excitability by inhibiting local SST^+^ and PV^+^ neurons and nearby VIP^+^ neurons, thus setting the level of local inhibition ^17,64^. Their recursive connectivity makes them good candidates for top-down control via long-range input from e.g. frontal cortical regions ^70–72^. VIP^+^ originate in a different location compared to PV^+^ and SST^+^ neurons, namely the Caudal Ganglionic Eminence rather than the Medial Ganglionic Eminence ^73^, and express their marker early in development ^57^. However, similarly to SST^+^ neurons, VIP^+^ neurons are mapped in adulthood but not in adolescence ^11^. We therefore used our processing pipeline to map the density of tdTomato-labelled VIP^+^ neurons throughout the mouse cortex.

In our dataset, we found that VIP^+^ neurons were present almost exclusively in the Isocortex, with most neurons in the upper half of cortical layer 2/3 (Fig 3c, 4b-c), consistent with previous reports which reported a predominantly upper-layer cortical location ^11^. VIP^+^ neuron density was lower than SST^+^ or PV^+^ in the same cortical regions, with a general trend towards higher density at the edge of the cortical plate (Fig 4b, consistent with ^11^). We found a small subset of VIP^+^ neurons in the hippocampus, with a ventral high - dorsal low gradient (Fig 3d). VIP^+^ neurons were almost completely absent from all other brain structures, which is in agreement with earlier reports using a different imaging modality ^11^. Thus, here we provide the first whole-adolescent brain-mapping of VIP^+^ neurons. This dataset will be relevant for developmental studies to compare to other timepoints, or as control for neurodevelopmental disorder studies^74^.

## Materials and Methods

### Transgenic mice

For obtaining the datasets used as proof-of-principle in this report, we used reporter mice expressing a subpopulation-specific Cre Recombinase driver (see below) and a floxed tdTomato reporter (ai14, see below), on a mixed C57/Bl6J background. We crossed Cre homozygotes with tdTomato-flox homozygotes, and used the first-generation offspring that was heterozygous for both Cre and tdTomato-flox reporter expression. Specifically, we used the following mouse lines:

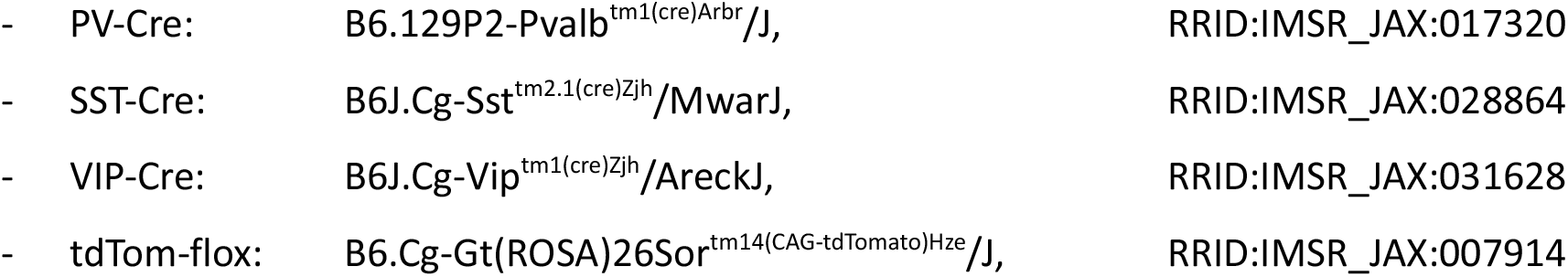

Mice were kept as described previously ^16^. In brief, we used mice from both sexes for our study. Animals were kept in wire-top cages (type III) and group-housed with the with the nest until weaning (P21-25), or and afterwards group-housed (3-6 animals, littermates, both genotypes, segregated by sex). Animals had access to rodent chow and water *ad libitum* and were kept on a 12h/12h light-/dark cycle (lights on at 07:00 AM). Animal experiments were conducted in conformity with the Animal Care Committee of the Radboud University Nijmegen Medical Centre and the National Committee on Animal Experiments (CCD), The Netherlands, and conform to the guidelines of the Dutch Council for Animal Care and the European Communities Council Directive 2010/63/EU.

### iDISCO+ staining

We performed iDISCO+ stainings as described in ^16^: We processed one hemisphere per brain following the iDISCO+ histochemistry protocol ^4^ as described for adult brains, with all buffers according to the protocol and all incubation steps taking place on a shaker/rotor, in 5 ml screw-top Eppendorf tubes, and lasting 1h unless mentioned otherwise. For details, please see the protocol at protocols.io (dx.doi.org/10.17504/protocols.io.eq2lynnkrvx9/v2

dx.doi.org/10.17504/protocols.io.yxmvmn9pbg3p/v2

dx.doi.org/10.17504/protocols.io.dm6gpbdwdlzp/v1

dx.doi.org/10.17504/protocols.io.36wgq77m5vk5/v1)

Briefly, whole mouse brain hemispheres were dehydrated in a methanol gradient (from 20%to 100%), bleached in 5% H_2_O_2_ in methanol at 4°C overnight, then rehydrated. Hemispheres were subsequently permeabilized for 5-7 days at RT, blocked for 5-7 days at 37°C, then incubated with primary antibodies (Rabbit Anti-RFP, Rockland 600-401-379, 1:2000, 2 ml / sample) for 6 days at 37°C. Subsequently, brains were washed 5×1h + 1x overnight at RT, and incubated for 7 days with secondaries (Goat anti-rabbit Alexa-568, Invitrogen A11036, or Goat anti-Rabbit Alexa-647, Invitrogen A-21245 1:500, 2 ml / sample) at 37°C. Following 5x 1h + 1x overnight washing at RT, samples were dehydrated in a methanol gradient, then twice more in 100% methanol, 66% DCM/33% methanol, 2x 15 min 100% DCM, and finally cleared in 100% Dibenzyl Ether (DBE, Sigma) in airtight glass vials. Brains were typically transparent within 2h, and completely cleared overnight. Alexa fluorescence in the samples remained at useable levels for at least 12 months of storage in DBE and over (maximally) 3 imaging rounds.

| Name | Dilution | Supplier | Cat# | RRID | Used for |
| --- | --- | --- | --- | --- | --- |
| Goat anti-Rabbit Alexa 647 | 1:500 | Invitrogen | A-21245 | RRID:AB_2535813 | VIP-tdTom |
| Goat anti-Rabbit Alexa 568 | 1:500 | Invitrogen | A-11036 | RRID:AB_10563566 | PV-tdTom (P14 / P56), SST-tdTom |
| Rabbit anti-RFP | 1:2000 | Rockland | 600-401-379 | RRID:AB_2209751 | All |

### Imaging

The cleared samples were imaged on a LaVision Ultramicroscope II Light-sheet microscope outfitted with a NTK Photonics white-light laser and filter sets for 488 nm, 568 nm, and 647nm, imaged through a long-working distance objective (LaVision) at 1.1x magnification (effective 2.2x, NA 0.1), and recorded with an Andor Neo 5.5 cooled sCMOS camera. We imaged with a 480 nm signal for autofluorescence for alignment with the Allen Brain Atlas, and 560 nm to record the TdTomato-Alexa signal. We used a single light-sheet from one side at 0.54 NA, scanning at 2.95 / 2.95 / 3 μm x/y/z resolution (3 μm z-steps) with the “horizontal focus” method and 17-18 horizontal focus steps. The sample was imaged submerged in DBE in sagittal configuration, and the entire cortex fit inside a single field of view (x/y), with a typical brain producing ~1600 z-planes of 3 μm each.

### System requirements

We recommend minimum system requirements of 16 GB Ram, >100GB Swap space on a fast SSD, and a modern x86 Processor with >4 cores (most modern Intel or AMD processors should work), with hardware virtualization (e.g. Hyper-V) enabled. Administrator access is required for installing the Docker environment (Docker Desktop), if it is not already present. We verified that the Docker containers with the core FriendlyClearmap pipeline run on Linux (Ubuntu 20.04 and 22.04), Windows (Windows 10 and Windows Server on Amazon AWS) and MacOS (MacOS 12.6.2 Monterey).

For Windows, installation of the Windows Subsystem for Linux (WSL2) is required prior to installing Docker Desktop. For Windows on PCs/Workstations, enabling Hyper-V might require accessing the machine’s UEFI. When using a Windows Server environment on a cloud instance such as Amazon AWS, a “bare metal” instance with enabled Hyper-V is required, as non-bare metal instances at present (February 2023) do not allow nested virtualization via WSL2.

We tested MacOS on a Mac Pro 6 (late 2013) with a quadcore Intel Xeon E5 processor and 16 GB of RAM, running MacOS 12.6.2 Monterey. In order to accommodate the 16 GB of RAM, we reduced the parallel processing settings for downsampling and processing to 1 (in the parameter_file_Ilastik_template.py for ClearMap1 and the process_Ilastik_run_this.py for ClearMap2) and set RAM and swap to maximum values in the Docker desktop “Settings” tab. We also tested installing Docker on an Amazon AWS Mac instance, however we could not successfully complete the docker desktop installation via ssh because finalizing the Docker Desktop installation requires GUI access.

## Supporting information

Supplementary protocol 1: Docker for Clearmap 1

Supplementary Protocol 2: Docker for Clearmap 2

Supplementary Protocol 3: iDISCO+ protocol for tdTomato

Supllementary Protocol 4: Visualize Data with BrainRender

## Data processing

- Project name: FriendlyClearMap
- Project home page: https://github.com/MoritzNegwer/FriendlyClearMap-scripts
- Operating system(s): Platform independent (Linux / Windows / MacOS, on x86 processors)
- Programming language: Python 3
- Other requirements: Docker runtime
- License: GNU GPL v3

## Data availability

Data will be available upon publication. Until then, find the containers, install scripts and a proof-of-concept dataset on dropbox: https://www.dropbox.com/scl/fo/rz1sm2egzc5iqtyddn7qy/h?dl=0&rlkey=rhhowpjt6df13fir7g3sy9qz4

## Acknowledgements

The tdTom-flox mice were a kind gift from Michael Valente, RIMLS Nijmegen. The PV-Cre and SST-Cre mice were kindly provided by Tansu Celikel, Radboud University Nijmegen (now Georgia Tech). We furthermore would like to thank Bram Geenen, Gert-Jan Bakker, Shaghayegh Abghari, Dewi van der Geugten, Katharina Foreman and Kari Bosch for helpful discussions during the iDISCO+ setup and imaging. We would like to thank Astrid Oudakker and Chantal Bijnagte-Schoenmaker for helping with ordering supplies and genotyping, and members of the Centraal Dierenlaboratorium for support with mouse breeding and keeping. We thank Thom Oostendorp for access to his Mac Pro for testing.

## Work Distribution

MN, DS, NNK conceptualized the study. NNK acquired funding. MN, MB, LL, RH and LA executed all iDISCO+ stainings and imaging. MN, BB and CR updated ClearMap1 to Python 3.8, set up Arivis Vision4D segmentation and extended the pipeline. MN built the Docker containers and integrated BrainRender. DS and NNK co-supervised students in this project. MN and DS generated figures. MN analyzed data and wrote the manuscript, MN, NNK and DS revised manuscript.

