## Supplementary protocol 1: Docker for Clearmap 1 for "FriendlyClearMap: An optimized toolkit for mouse brain mapping and analysis"

CN53VG8N

### Run Clearmap 1 docker V.(cn53vg8n)

Moritz Negwer[1]

[1]Radboudumc Nijmegen

M Moritz Negwer

DISCLAIMER

Dropbox links are temporary and will be replaced eventually.

**Protocol Info:** Moritz Negwer . Run Clearmap 1 docker. **protocols.io** https://protocols.io/view/run-clearmap-1-docker-cn53vg8nVersion created by Moritz Negwer

**Created:** Feb 09, 2023

**Last Modified:** Feb 15, 2023

**PROTOCOL integer ID:** 76699

ABSTRACT

This protocol is a supplement to our upcoming publication "FriendlyClearMap: An optimized toolkit for mouse brain mapping and analysis".

In this protocol, we describe in detail how to run Clearmap1 in a Docker container.

#### Docker Setup

1

**On Windows:**
If you haven't already, download and set up Docker Desktop for Windows. This requires administrator privileges and will also install the Windows Subsystem for Linux (WSL2).

Download:
https://www.docker.com/products/docker-desktop

Alternatively, use our install script via powershell: github link

Download the docker container from our repository:
Dropbox

Load the docker image with

```
docker image load -i ./friendlyclearmap_4_2.tar
```

Download and install Ilastik for Windows:

https://www.ilastik.org/download.html

Download and install FIJI for Windows:
https://imagej.net/software/fiji/downloads

Update FIJI to the latest version
  FIJI Plugin update: FIJI -> Help -> Update, then restart, click Apply Changes
  FIJI program update: FIJI -> Help -> Update ImageJ

Verify that BigWarp is installed as FIJI plugin. It should be installed by default on newer FIJI versions.
https://imagej.net/plugins/bigwarp
(FIJI -> Plugins -> BigDataViewer -> BigWarp)

**On Linux:**

Install docker as described here: https://docs.docker.com/engine/install/ubuntu/

Alternatively, use our install script: github repo

Download the docker container from our repository:
Dropbox

Load the docker image with

```
docker image load -i ./friendlyclearmap_4_2.tar
```

Download and install Ilastik for Linux
https://www.ilastik.org/download.html
(we recommend a direct install of the newest Beta, as the versions in e.g. Ubuntu repositories often lag a few versions behind)

Download and install FIJI for Linux:
https://imagej.net/software/fiji/downloads

Update FIJI to the latest version
  FIJI Plugin update: FIJI -> Help -> Update, then restart, click Apply Changes
  FIJI program update: FIJI -> Help -> Update ImageJ

Verify that BigWarp is installed as FIJI plugin. It should be installed by default on newer FIJI versions.
https://imagej.net/plugins/bigwarp
(FIJI -> Plugins -> BigDataViewer -> BigWarp)

**On MacOS:**
Download Docker Desktop for Mac and follow the instructions: Download link
(N.B. we only tested Intel Macs, use Apple Silicon Macs at your own risk)

Alternatively, use our install script: github repo

Then, download the docker container from our repository:
Dropbox

Load the docker image with

```
docker image load -i ./friendlyclearmap_4_2.tar
```

Download and install Ilastik for MacOS
https://www.ilastik.org/download.html
(we recommend a direct install of the newest Beta that is compatible with your MacOS version).

Download and install FIJI for Linux:
https://imagej.net/software/fiji/downloads

Update FIJI to the latest version
FIJI Plugin update: FIJI -> Help -> Update, then restart, click Apply Changes
FIJI program update: FIJI -> Help -> Update ImageJ

Verify that BigWarp is installed as FIJI plugin. It should be installed by default on newer FIJI versions.
https://imagej.net/plugins/bigwarp
(FIJI -> Plugins -> BigDataViewer -> BigWarp)

#### Getting your data ready

2 This pipeline assumes that the samples are saved as a series of single-channel tiffs, one per z-plane.

This means that if you have tiled images, you will need to stitch them so that there is only one per z-plane. This can be done with e.g. the FIJI stitching plugin (Fiji -> Plugins -> Stitching -> Deprecated -> Stitch Sequence of Grids of Images.

Double-check that the folder with the tiff series does only contain this series and nothing else (i.e. tiling information or other images). If you have multiple channels,

At a minimum, this pipeline expects a single channel with the signal of interest (e.g. tdTomato or cFos). The autofluorescence channel is used in the original ClearMap 1/2 pipelines for atlas alignment only - if your signal channel has some autofluorescence, it can be used for atlas alignment as well. If you recorded an autofluorescence channel, you can use this for atlas alignment as well.

For fast processing, we recommend at a minimum 32GB RAM and an SSD for fast access. We tested this pipeline on a Laptop (64GB RAM, Ryzen 5 4600H 6 core / 12 threads, SSD) and a desktop PC (32GB RAM, Ryzen 5 3200G 4c/8t, external SSD via USB3), both running Windows 10.

#### Setting up Segmentation with Ilastik (optional)

3 **Generate a substack for Ilastik with FIJI:**

In FIJI, load a representative section of your image stack. We recommend >200 z-planes at 3 µm z-height, from the middle of the stack.

FIJI -> Import -> Image Sequence
  Select your folder
  Set Start image: e.g. 800 if you have 1600 z-planes
  Set count: 200

Once loaded, save as  a hdf5 image for Ilastik:
  FIJI -> Plugins -> BigDataViewer -> Export Current Image as XML/HDF5
  Leave settings as below:

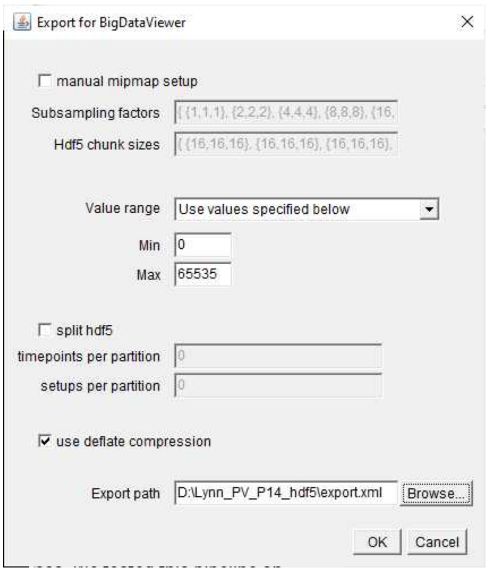


FIJI will then generate the image pyramid for the hdf5 stack and save it in the appointed location.

4 **Loading the data in Ilastik**

Generate a Pixel Classification workflow in Ilastik

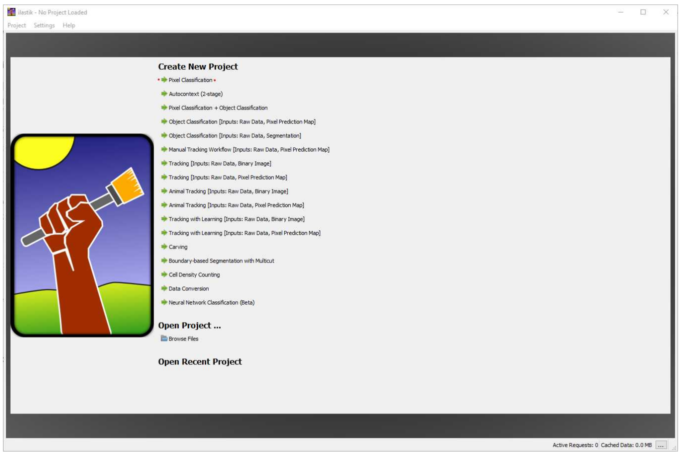


Load the hdf5 file via "Add Single Image", then select t0000/s00/1/cells to display the

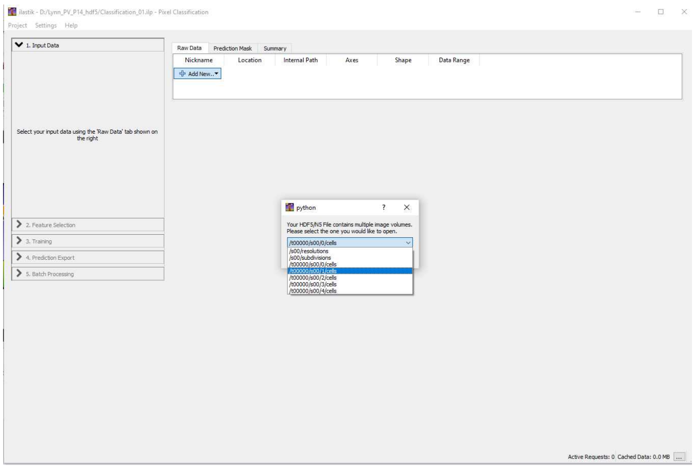

Copy the stack into the project file by right-clicking on the image -> "Edit", then selecting "Copy into project file". This results in larger Ilastik files, but is important for loading the same files in the docker environment (relative paths will not be preserved).

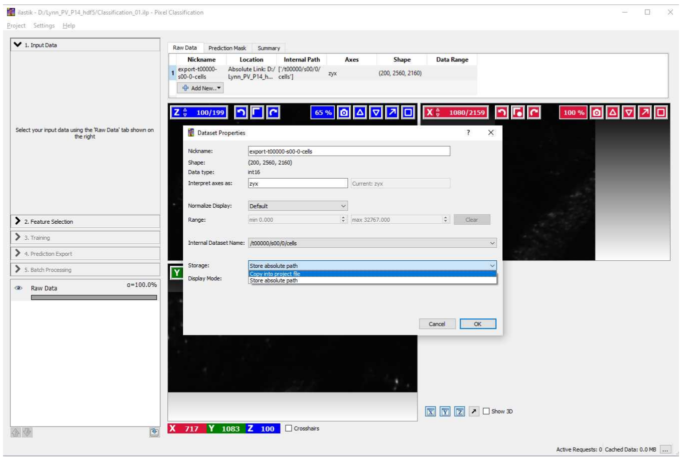

If you want to adjust the brightness of the displayed images, you can do so by right-clicking on "Raw Data" in the left column -> "Adjust Thresholds". Then uncheck "Automatic Range" and set the maximum value so that your cells of interest are visible.

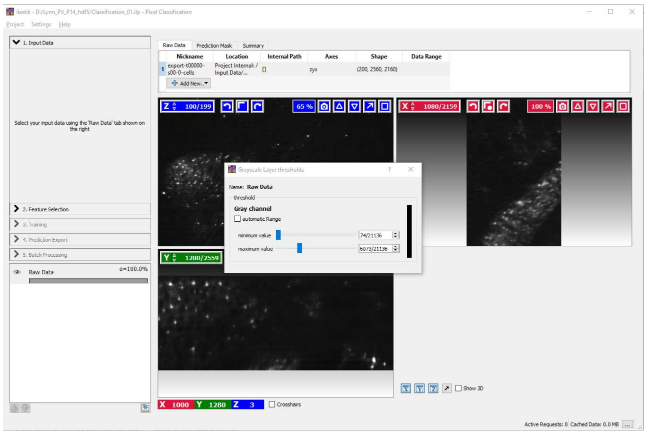

#### 5 Segmentation with Ilastik

In the "Select Features" pane, add a 30 (pixels wide) filter by clicking to the right of the 10 column and entering 30.
Then, select the "Edge" and "Texture" boxes for all σ (filters).

(Note that those are empirically determined settings. Adding a 30px filter approximately captures single neurons when imaged at 1.1x magnification on a 2160 x 2560 pixel camera. Different imaging conditions, notably with different magnifications, might require different-size filters. Also, skipping the intensity-based filters speeds up calculations without noticeable worse segmentation quality.)

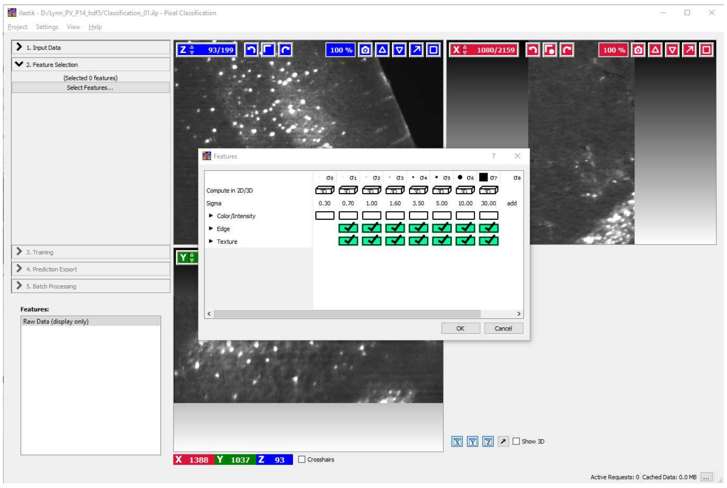

##### 5.1 In the "Training" pane, define the background (Label 2)

- Zoom out in the XY projection (Ctrl + mousewheel) until the entire brain is visible
- Select Label 2, set to max size (61) and draw along the empty space besides the brain. Take

care not to draw over the edge of the image, as that can lead to problems down the road with some Ilastik versions.
- Subsequently, draw single dots along the edges of the brain and in the neuropil where there are no labelled cells.

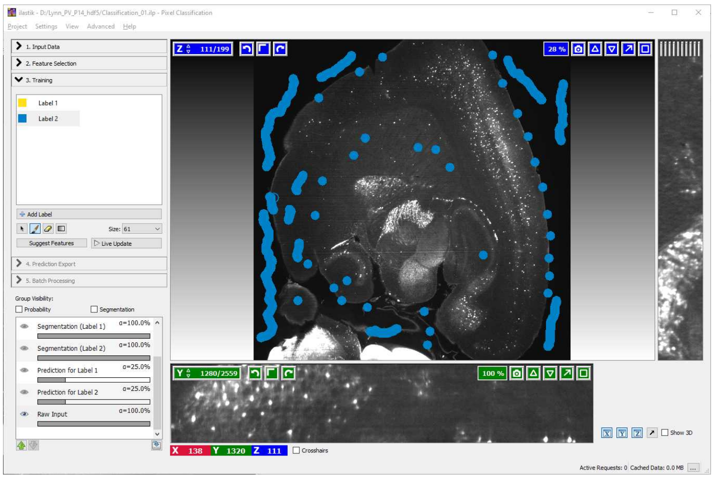


5.2 Subsequently, zoom in on your signal (Ctrl + Scroll wheel). To syncronze the view between the XY projection and the other two, double-leftclick on an object of interest, e.g. a cell. The voxel you clicked will then be centered in the other projections as well.

1) Select a few cells as signal (Label 1, yellow). For our images, a size 5 brush covered most of the labelled cell bodies at their maximum extent.

2) Surround the selected cell with a thin circle of background (1-3 px wide). We found this to be especially helpful when marking cells close to each other, as otherwise Ilastik may wrongly mark the region in between also as signal.

3) Repeat for the same cells for the XZ and YZ projections.

4) We recommend marking ca. 100 cells in total, spread out over the stack. Take care to select regions that are representative of the entire brain, i.e. with different densities and background intensities.

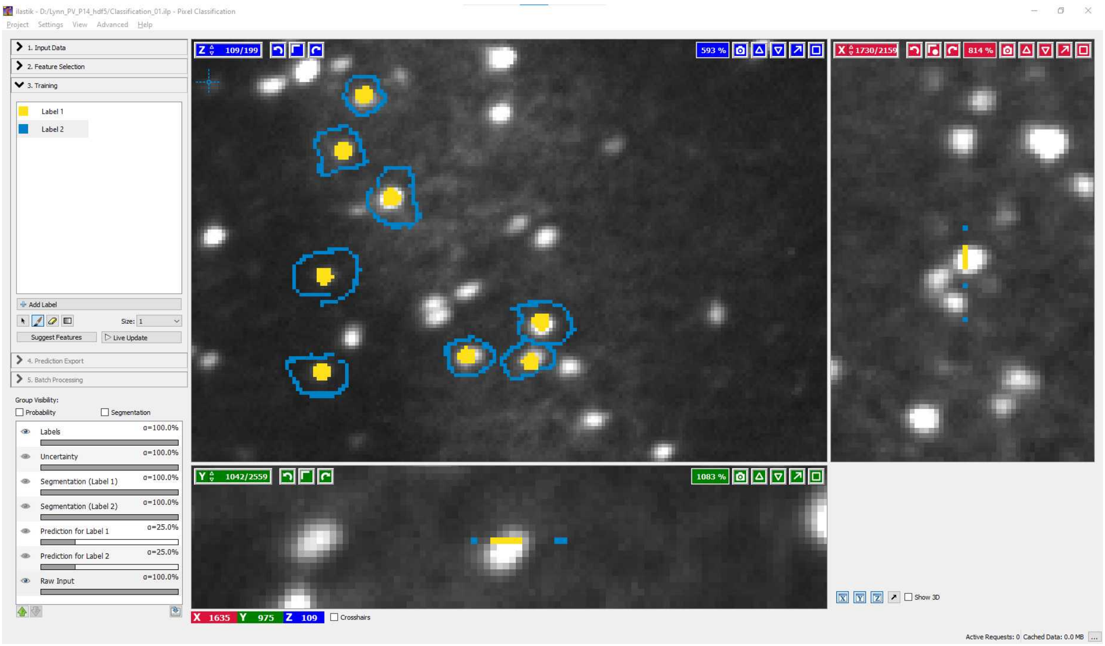


5.3 Once you have marked ca. 100 cells, zoom in on one region in all three projections and click "Live Preview" to trigger the training of the random forest algorithm with your input data.

**Note:** Do *not* add new input data during training, as this will trigger a re-training from the start.

Depending on the number and speed of your CPU cores, this may take a long time (10-45 minutes) to complete. Once it is completed, the prediction will be overlaid over your field of view.

Once the training has finished, verify that the segmentation is correct. If not and more training data needs to be added, **turn off "Live Preview" before adding new data** - otherwise, every new addition will trigger a fresh round of training, leading to long delays.

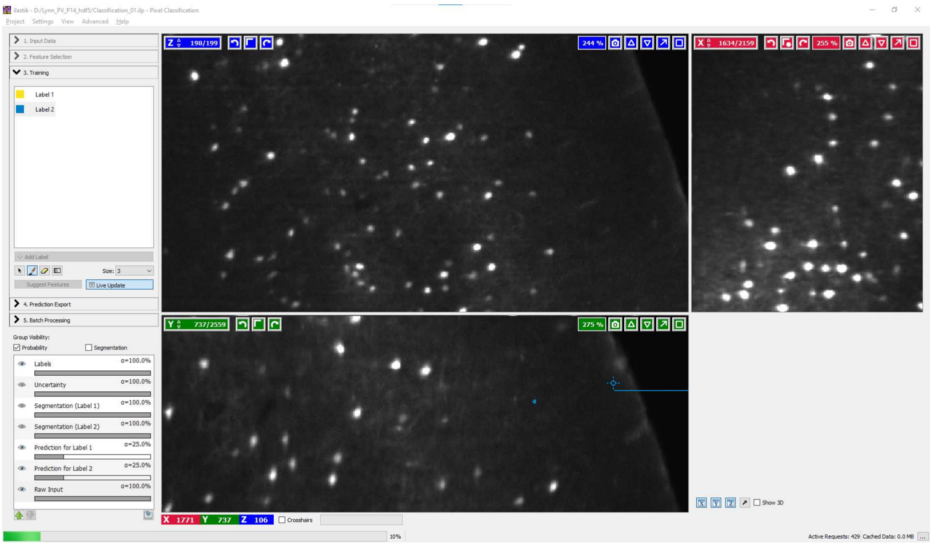

Training in progress. Note the progress bar in the bottom left corner.

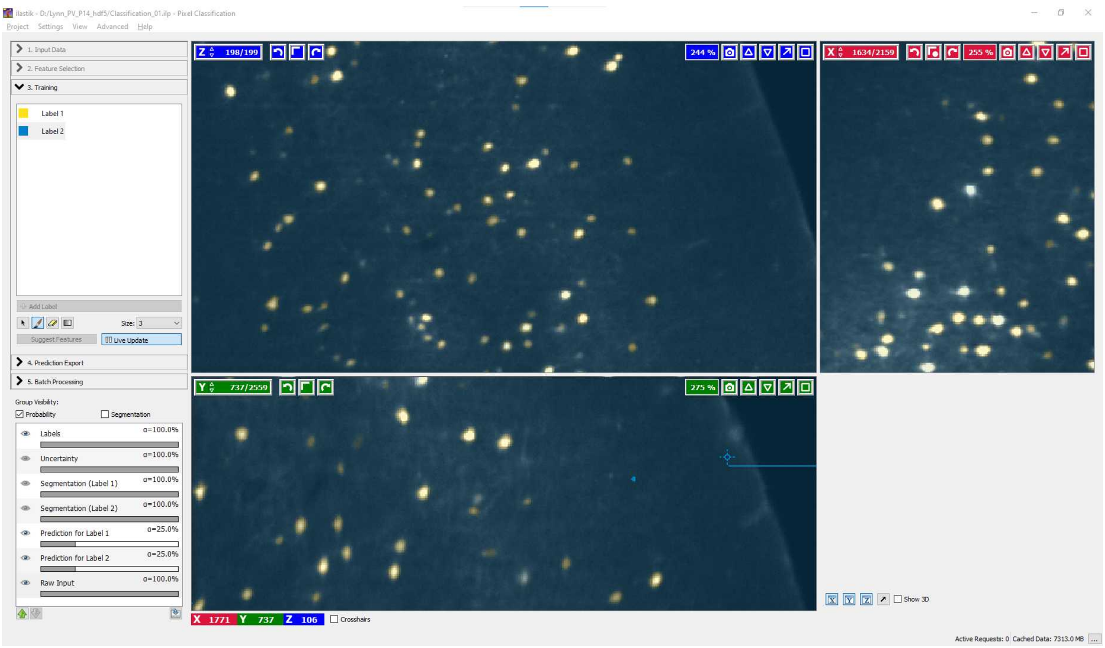

Training finished. Note the cell in the center of the XZ projection is predicted to be background. However, scrolling through the XZ projection will reveal that the following plane is segmented as a cell. Therefore, no action is needed.

#### 5.4 Auto-selecting the best features for training (optional)

Ilastik offers a "Suggest Features" option. After successful training, this option compares subsets of features to find which reduced subset still represents the training data.

Doing so can take a long time (several hours), but once done, it reduces the set of required features from the 14 selected in Step 5 of this example to a smaller amount (default 7).

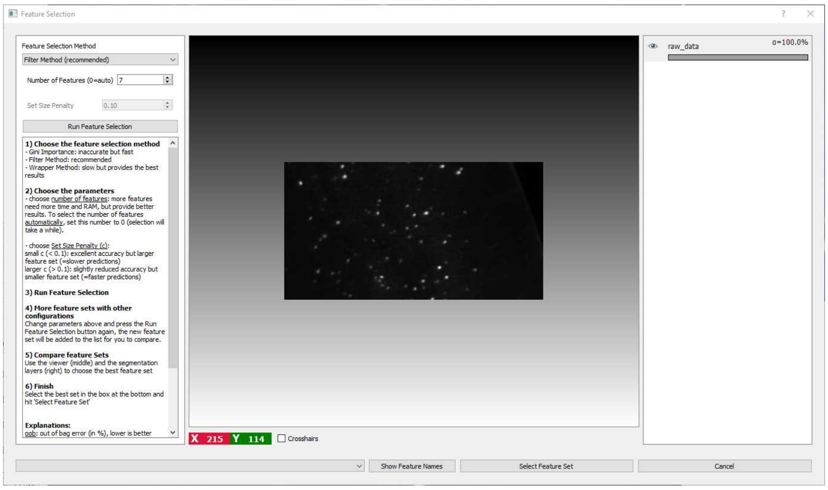


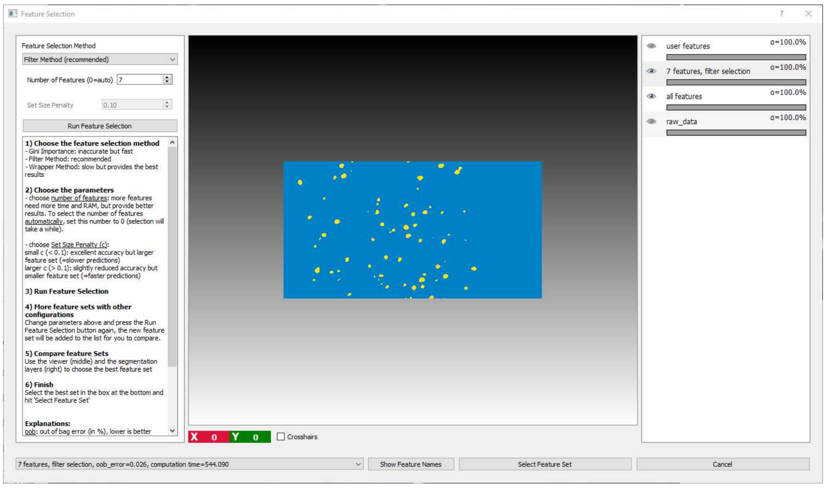


Compare the feature by clicking on the eye symbols in the left column. If you agree with the selected subset ("7 features"), change the features in the bottom right drop-down menu. There, you can also see the computation time as a comparison. In our example case, the auto-selected subset is approx. 40% faster than the full set of features.

Finally, click "Select Feature Set" to confirm. Then, retrain the random forest algorithm with the new subset of parameters by clicking "Live Preview" once more.

#### 5.5 Setting the Ilastik Export settings correctly

Once the training is complete, proceed to the "Prediction Export" pane. There, click "Choose Export Image Settings", and set the following parameters:

- Change "Convert to data type" to "unsigned 8-bit"
- Check "Renormalize [min,max] from 0.0 / 1.0 to 0 / 255

The rest can be left at default. For the final settings, see below:

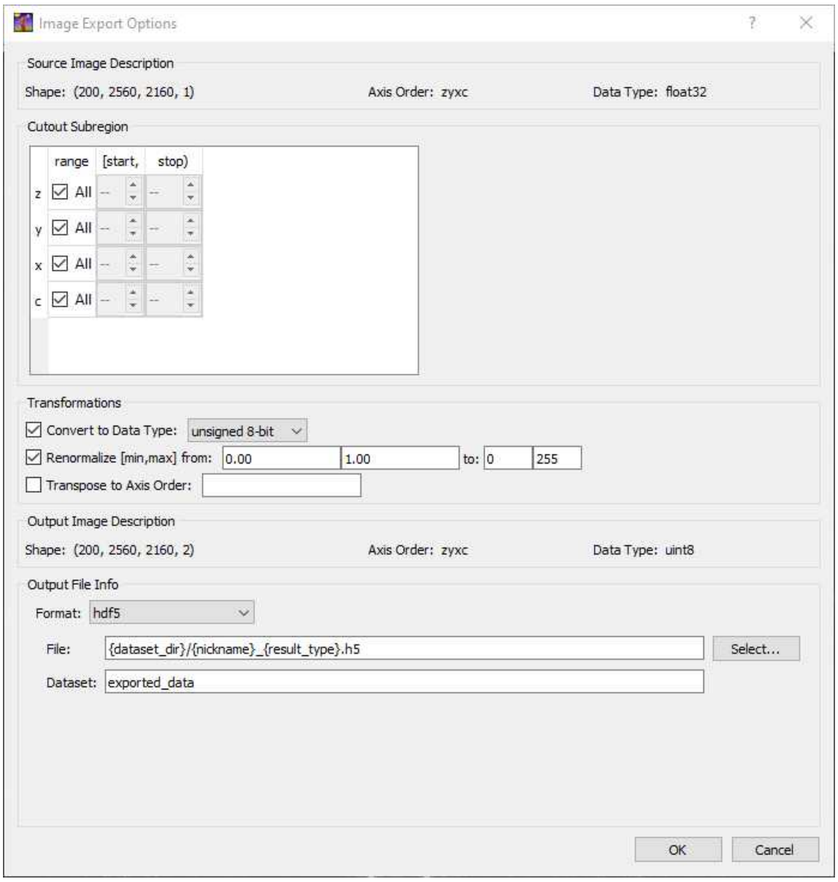


Confirm with OK. Then save (File -> Save), and close the project. It should now be ready for use in both Clearmap1 and Clearmap 2.

#### Segmentation with Arivis Vision4D Machine Learning Segm...

6 If you have an install of Arivis Vision4D with the Machine Learning Segmenter package, you can proceed as follows to generate a finished cell coordinate file.

6.1 Convert your dataset with the Arivis SIS converter

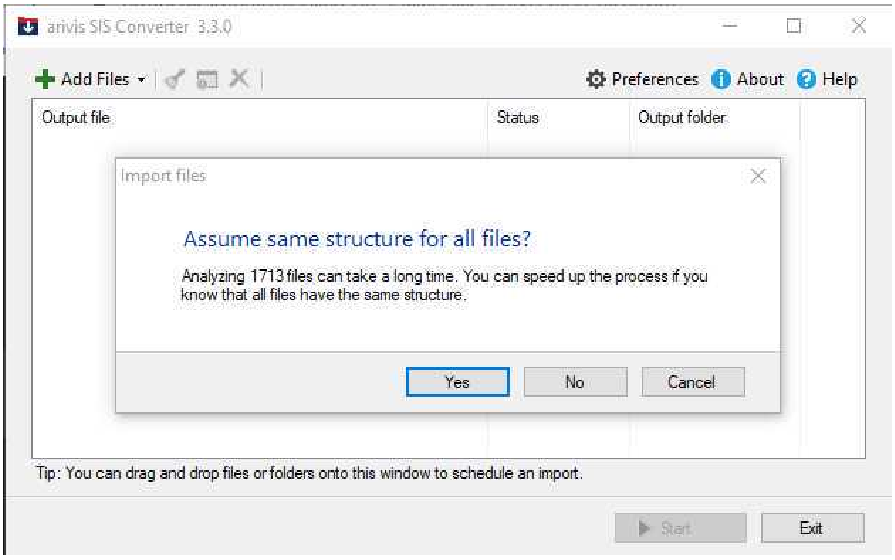


Load all files from the image stack as z-planes into SIS converter

6.2 Then, load them into Arivis, and in the left Processing Pane, add a "Machine Learning Segmenter" processing step. No pre-processing should be necessary.

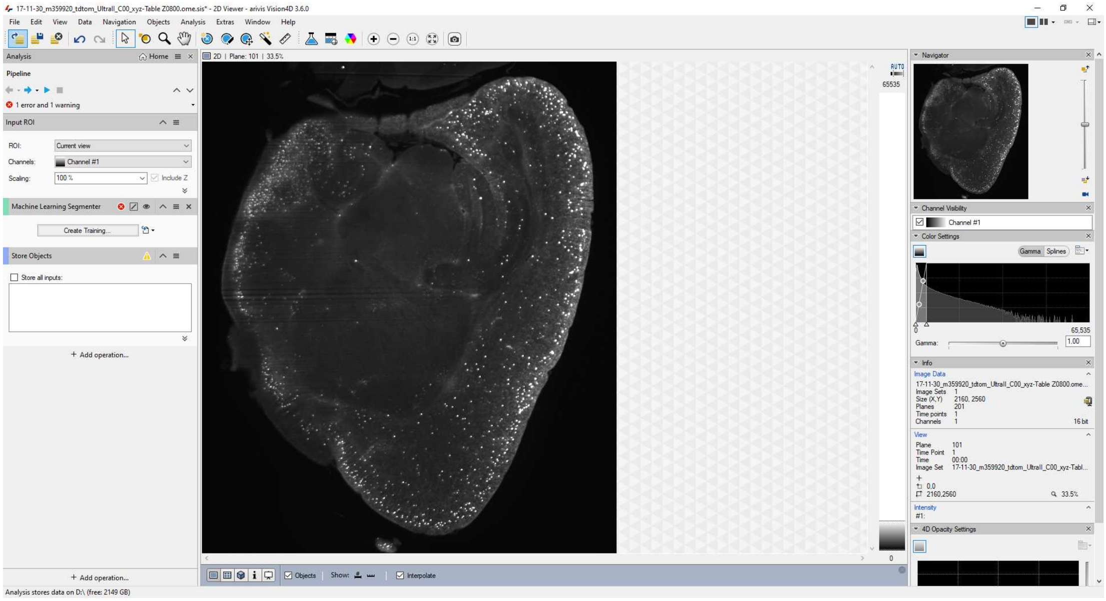

6.3

Subsequently, open the "Train Machine Learning Segmenter" pane and start adding foreground and background labels. We recommend the following:
- Make sure to assign some background label to the blank space around the brain and the edges of the sample.
- Mark a representative subset of cells (we recommend 100 throughout the brain).

- Mark a small circle of background around the marked cells. In our experience, this helps considerably with separation of dense cells.
- Make sure to mark dense projections, fiber tracts, membranes, blood vessels, etc. as background.

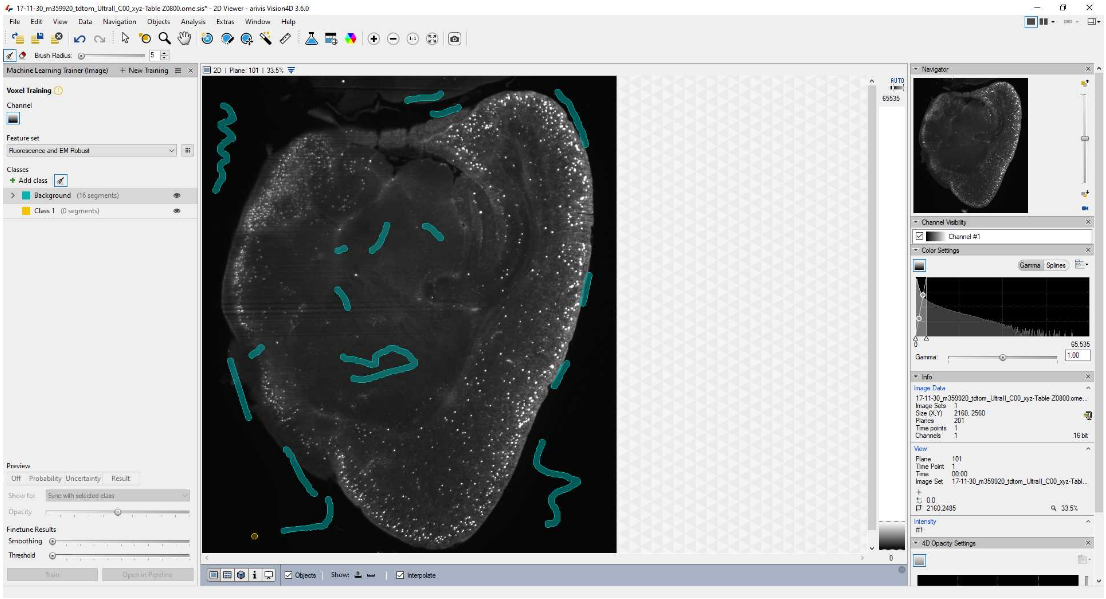


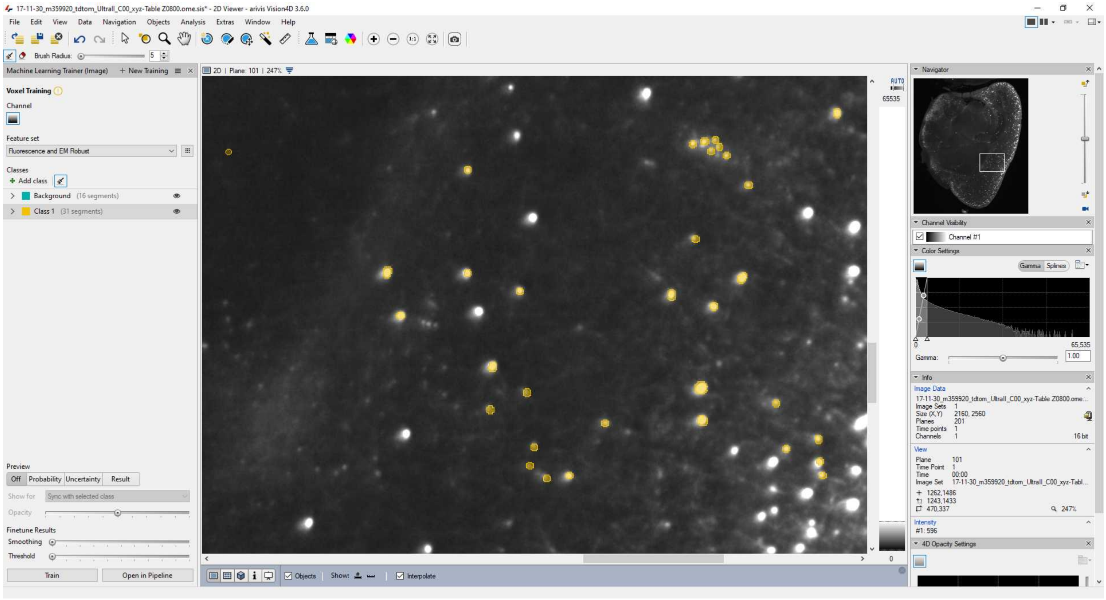

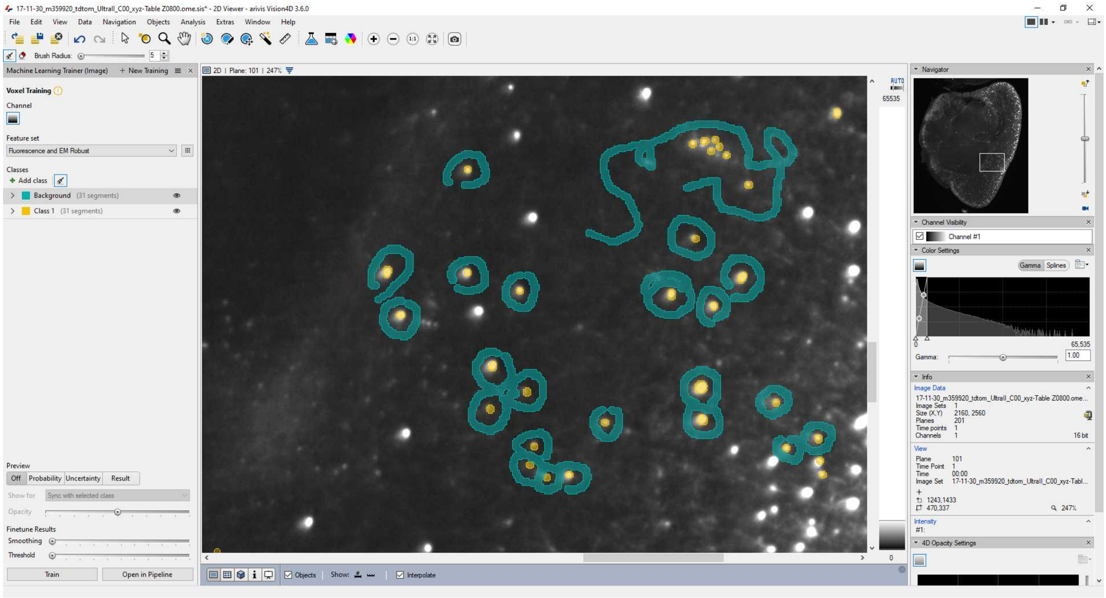

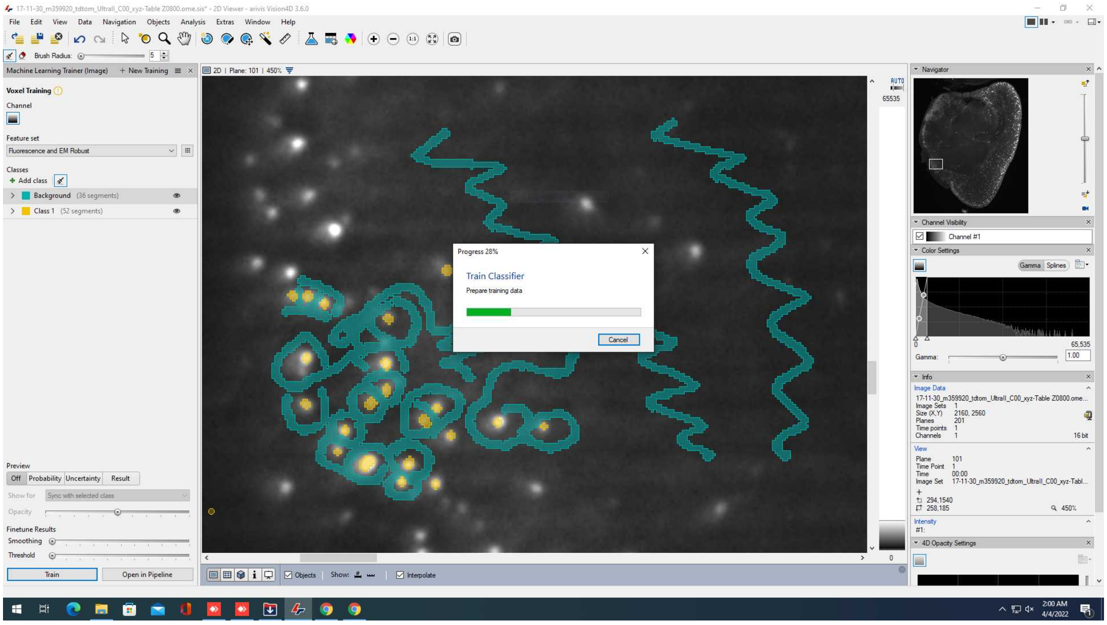


(If you already have a trained Ilastik file, you can also import it via the "Import from Ilastik" option).

Once trained, save the training so it can be reused.

**6.4** In the pipeline pane on the left, make sure to change the Input ROI to "current image set" (otherwise it will remain at the default "current view", i.e. the 2D cutout that you are viewing.

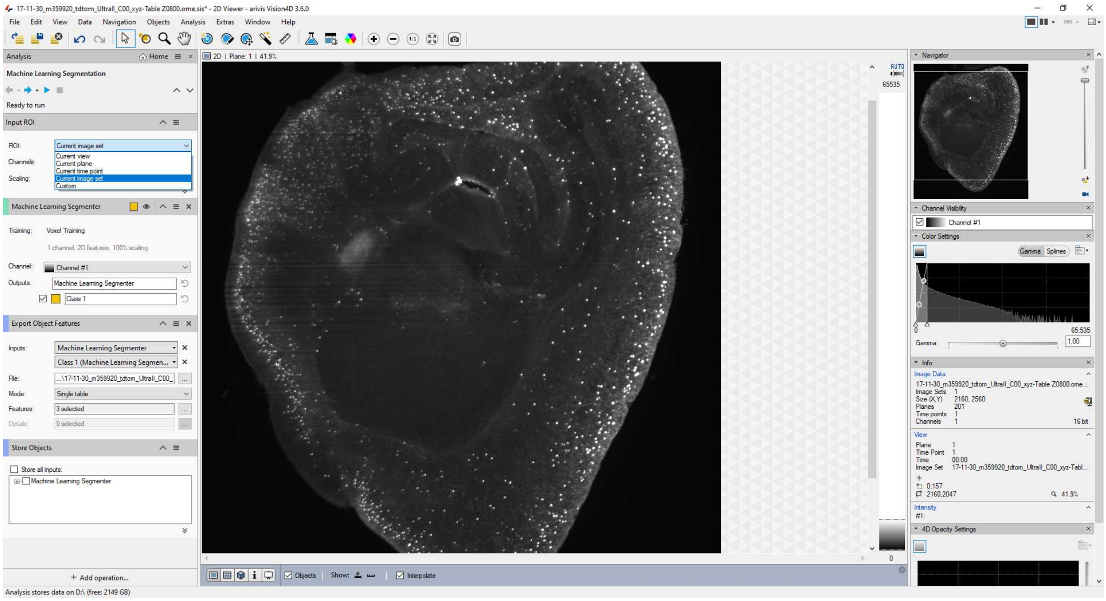

To export as a coordinate list, add a "Export Objects Features" step to the processing pipeline.

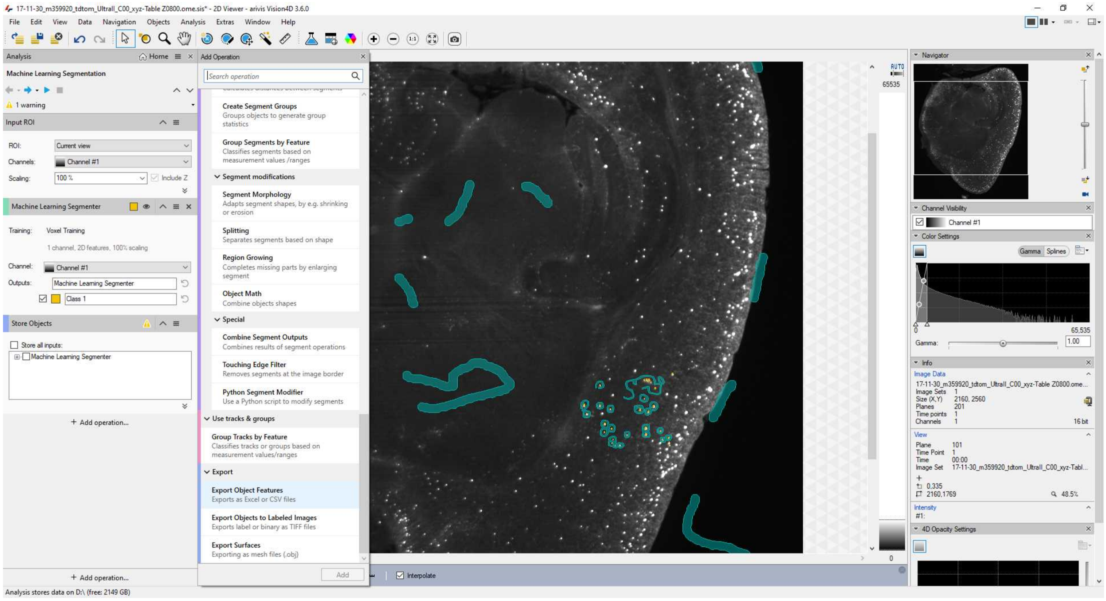

Specify the output as an Excel file. Select the following Object Properties:

- Deselect everything (defaults won't make much sense)
- Center of Gravity X / Y / Z (px)

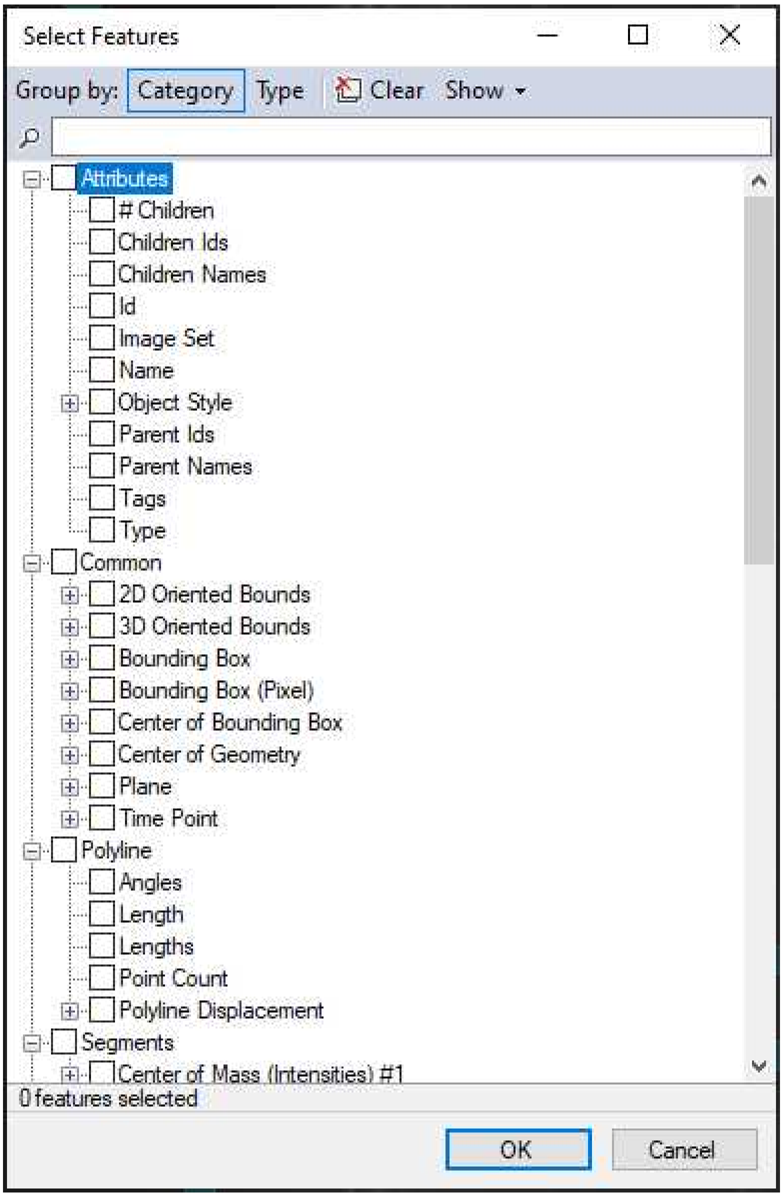
Select Features
Group by: Category Type Clear Show
Attributes
### Children
Children Ids
Children Names
Id
Image Set
Name
Object Style
Parent Ids
Parent Names
Tags
Type
Common
2D Oriented Bounds
3D Oriented Bounds
Bounding Box
Bounding Box (Pixel)
Center of Bounding Box
Center of Geometry
Plane
Time Point
Polyline
Angles
Length
Lengths
Point Count
Polyline Displacement
Segments
Center of Mass (Intensities) #1
0 features selected
OK
Cancel

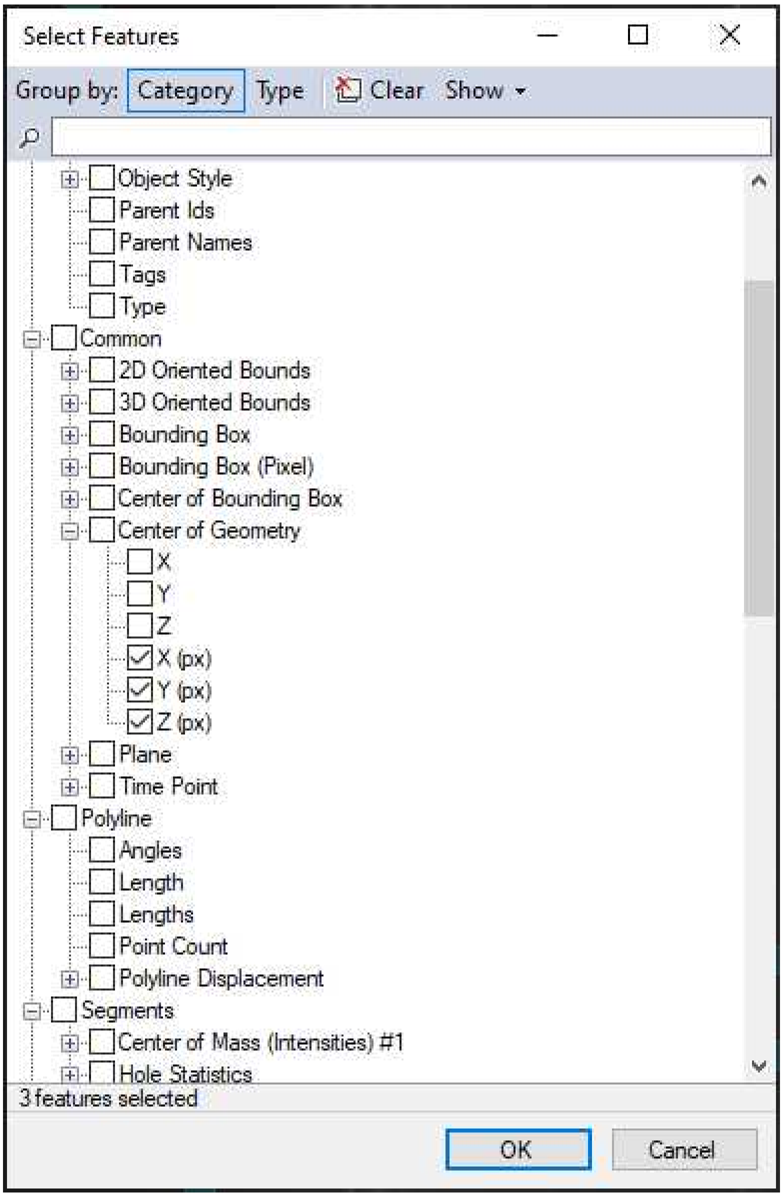


6.5 Save the pipeline.

Then, run it for the file at hand. The Machine Learning Segmenter plugin is CPU-heavy and will likely take approx. 24h for a 2160 x 2560 x 2000 pixel stack on a recent (14 core, 128GB RAM, fast NVMe SSD) workstation.

If you want to optimize Arivis' speed, make sure that the cache is written to a SSD.

Furthermore, the objects file is located in a SQL databse in the same folder as the SIS file. We found that with >100k cells, random read/write access to the database is slowing down processing, easily adding several days of processing time per file. We recommend to at least have the SIS file on a fast internal SSD.

If you have sufficient RAM to spare (total >64GB), you can create a Ramdisk (e.g. with ImDisk, [https://sourceforge.net/projects/imdisk-toolkit/](https://sourceforge.net/projects/imdisk-toolkit/)) and copy your SIS file there before starting

processing. This should considerably speed up processing time for samples where you expect a large number (more than 100k) of segmented cells.
As Arivis Vision4D itself is very conservative with RAM use (never exceeded 15 GB with our Machine Learning Segmenter-based pipeline), you shold be able to set aside everything above 32GB as a Ramdisk.
**Note:** Make sure to copy your files back onto a SSD or HDD before proceeding with the next stack. Ramdisks are volatile and their contents will disappear if the Ramdisk is unmounted or the workstation is shut off.

#### Atlas alignment with Dockerized Clearmap pipelines

7 First, set up your folder structure as follows:

1) One folder per sample
2) In this folder, have one subfolder per channel, containing only the image stack (one tiff image per z-plane)

(Note: Make sure that there are no spaces in the path to this folder names, as this will trip up Elastix when running the alignment. If you need to fix the filenames, try Bulk Rename Utility (https://www.bulkrenameutility.co.uk/Download.php) on Windows. On Linux use command line or the Thunar file manager (standard in XFCE-based distributions such as Xubuntu)).

7.1 Copy the parameter_file_Ilastik_template.py into the folder and adapt as follows:

```
cFosFile = os.path.join(BaseDirectory,
    'folder_with_signal_of_interest/
    name_of_the_first_tif_Z\d{4}.ome.tif');

#Example: os.path.join(BaseDirectory,
#             '200525_VIP_P26_m69366_TdTom_13-28-31/
#             13-28-31_VIP_P26_m69366_TdTom_UltraII_C00_xyz-Table
Z\d{4}.ome.tif');
```

In this case, the \d{4} signals that there are four numbers after the Z , starting from 0000.

```
Autofluofile = os.path.join(BaseDirectory,
    'folder_with_autofluo_stack/
    name_of_the_first_tif_Z\d{4}.ome.tif');

#If you only have one channel, uncomment the following:
#Autofluofile = cFosFile;
```

```
OriginalResolution = (2.95, 2.95, 3);
```

Adapt this to the dimensions of your scan.
In the case of LaVision / Miltenyi Ultramicroscopes, the tiff headers should contain a large amount of information. You can find this information by loading the first tiff file in FIJI and clicking Image -> Show Info.  In that case, save the content of the "File Info" window (File -> Save As, then choose a name and save as .txt).
Then re-open the resulting .txt file with a text editor such as Notepad++ (https://notepad-plus-plus.org/downloads/) on Windows or gedit on Linux. Search for "PhysicalDimension" and note the X, Y, and Z values.

```
#Resolution of the Atlas (in um/ pixel)
AtlasResolution = (25, 25, 25);
```

If you are using a custom reference atlas with a different resolution (e.g. 10 x 10 x 10 µm/voxel) , adapt the AtlasResolution values.

```
#Path to registration parameters and atlases
PathReg        = '/CloudMap/atlas_p21/';
AtlasFile      = os.path.join(PathReg, 
'Kim_ref_P21_v2_brain.tif');
LabeledImage   = os.path.join(PathReg, 
'Kim_ref_P21_v2.1_label.nrrd');
AnnotationFile = os.path.join(PathReg, 'regions_IDs.csv');
```

Adapt the AtlasFile, LabeledImage and AnnotationFile parameters to match the name of your atlas files.

**Note:** Specifically for the regions_IDs.csv, double-check the character encoding - a file prepared on Windows with Excel might get saved in a windows-specific character encoding, which means that the é or ö in names such as "dorsal Raphé" or "Pre-Bötzinger Complex" might cause issues later on. We recommend to double-check the file with Notepad++ or gedit and manually setting the character encoding to UTF-8 (Encoding -> UTF-8) to prevent errors.

```
ImageProcessingMethod = "Ilastik";

ilastikParameter = {
    "classifier" :
"/CloudMap/Classifiers/name_of_your_Ilastik_classifier.ilp",
    "classindex" : 0,
    "save"       : None, # (str or None)       file name to save
result of this operation if None dont save to file
    "verbose"    : True       # (bool or int)       print / plot
information about this step
}
```

First, adapt ImageProcessingMethod ("Ilastik" or "Builtin" or "Import")

If using Ilastik, also adapt the classifier name to the file name of your Ilastik .ilp project: Replace "name_of_your_Ilastik_classifier.ilp" with the proper filename.
(Note that the path does not need to be changed, as this will be changed when running the Docker container)

```
#Stack Processing Parameter for cell detection
StackProcessingParameter = {
    #max number of parallel processes. Be careful of the memory
footprint of each process!
    "processes" : os.cpu_count(),

    #chunk sizes: number of planes processed at once
    #optimized for 64GB RAM, can be higher if more available
    "chunkSizeMax" : 400,
    "chunkSizeMin" : 100,
    "chunkOverlap" : 20,

    #optimize chunk size and number to number of processes to
limit the number of cycles
    "chunkOptimization" : True,

    #increase chunk size for optimization (True, False or all =
automatic)
    "chunkOptimizationSize" : all,

    "processMethod" : "parallel"
   };
```

Those scripts are for a 12 core / 64 GB RAM system. If you find that the Docker container consumes too much RAM and does not finish the script, reduce the following values:

"processes" : int(os.cpu_count()/2), # this reduces the number of parallel threads to half the available number of processor cores

Then proceed to halve chunkSizeMax, chunkSizeMin and chunkOverlap until the Docker container runs.

8 Copy the 01_preprocess_process_Ilastik_run_this.py file into the folder and **rename into process_Ilastik_run_this.py**

Specifically, the file should look like this:

```
# -*- coding: utf-8 -*-
"""
Template to run the processing pipeline
"""

import os,numpy, sys

#pre-execute to make sure that Clearmap modules can be loaded
basedir = os.path.abspath('/CloudMap/FriendlyClearmap/clearmap/')

#insert into system folder
sys.path.insert(0, basedir)

import ClearMap

#load the parameters:
#with parameter file in the data folder
parameter_filename =
'/CloudMap/Data/parameter_file_Ilastik_template.py'
exec(open(parameter_filename).read())


#resampling operations:
#######################
#resampling for the correction of stage movements during the
acquisition between channels:
resampleData(**CorrectionResamplingParameterCfos);
resampleData(**CorrectionResamplingParameterAutoFluo);

#Downsampling for alignment to the Atlas:
resampleData(**RegistrationResamplingParameter);
```

This will generate (by default) three image stacks downsampled to 25µm/voxel, suitable for alignment to the Allen Brain Atlas.

9 Subsequently, run the Docker container for the first time. If everything goes according to plan, this will generate three additional tiffstacks downsampled to 25µm/voxel, suitable for alignment to

the Allen Brain Atlas. The Docker container should finish after running.

**On Windows**, open power shell, navigate to the folder with the process_Ilastik_run_this.py script and adapt the following code:

```
docker run -it --rm -v C:\path\to\brain:/CloudMap/Data
-v C:\path\to\custom\Ilastik\file:/CloudMap/classifiers
friendly_clearmap:3.0 ;
```

Note that file paths must not end on a trailing backslash.

**On Linux**, open a terminal, then navigate to the folder with the process_Ilastik_run_this.py script and run the following command:

```
docker run -it --rm -v /path/to/brain:/CloudMap/Data
-v /path/to/custom/Iastik/file:/CloudMap/classifiers
friendly_clearmap:3.0
```

**On MacOS**, open a terminal (Spotlight -> press Space -> type " Terminal"), then navigate to the folder with the process_Ilastik_run_this.py script and type the folllwing command:

```
docker run -it --rm -v /path/to/brain/:/CloudMap/Data
-v /path/to/custom/Iastik/file/:/CloudMap/classifiers
friendly_clearmap:3.0
```

By default, the container will execute the process_Ilastik_run_this.py script. If you want to run a different script, you can add the option

```
--entrypoint "python3 /CloudMap/Data/your_custom_script.py"
```

prior to the friendly_clearmap:3.0 argument.
Note that in this case the script needs to be in the folder that is mapped to /CloudMap/Data with the first -v command (i.e. the folder containing all image data).

#### 10 Atlas alignment using BigWarp

(In Windows/Linux, the Docker container should have finished)

1. In FIJI/Imagej. open both the atlas stack (e.g. template_25.tif) and the autofluo_resampled.tif files.

2. Open BigWarp (Plugins -> BigDataViewer -> BigWarp)

3. With BigWarp, open the atlas template as moving image, and autofluo_resampled as target image:

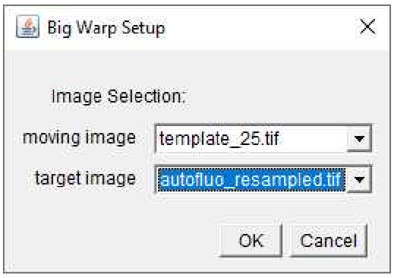


4. You should now see your autofluo_resampled on the left, and the atlas template on the right.

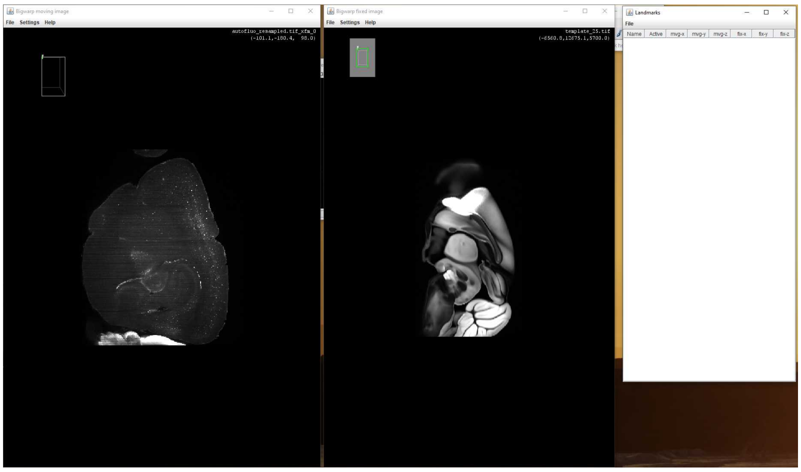

5. Go to the sagittal plane in both:
- Ctrl+A should change to a ZX projection
- Press Z to fix rotation around the Z (originally Y) axis
- Use the arrow keys to rotate around the axis until the orientations are roughly similar

6. Use the mosue wheel to scroll to a similar location in both images. Suggested:
    - Frontal Pole
    - Anterior Commissure crossing over
    - Maximum extent of the Somatosensory Cortex Barrel Field
    - Hippocampal landmarks, e.g. just before Fornix projects entirely ventrally
    - Posterior pole of V1
    - Cerbellar regions (if applicable)

7. Fine-tune the autofluo_resampled projection to match the coronal plane of the atlas template:
- Left-click on the point in the autofluo_resampled image that you would like to use as center of rotation
- Carefully move the mouse to rotate the cutting plane in 3D space. Repeat until it matches
- To reset, press Ctrl+A, then repeat step 5.

8. Generate landmarks:
- Press spacebar to start landmark mode.
- Identify corresponding points.
- Mark one point in autofluo_resampled, *then immediately* mark the corresponding point in the atlas template.

(Note that clicking several points in the same image will lead to non-matched point sets, which will not work for alignment later).
- Mark between 5 and 20 point pairs per location.
- Take care to mark a diversity of matched locations (e.g. pial surface, white matter, layer 4, hippcoampal landmarks)
- To change perspective (or undo landmarks), disengage landmark mode by pressing spacebar again. Re-engage once you are happy with the result.

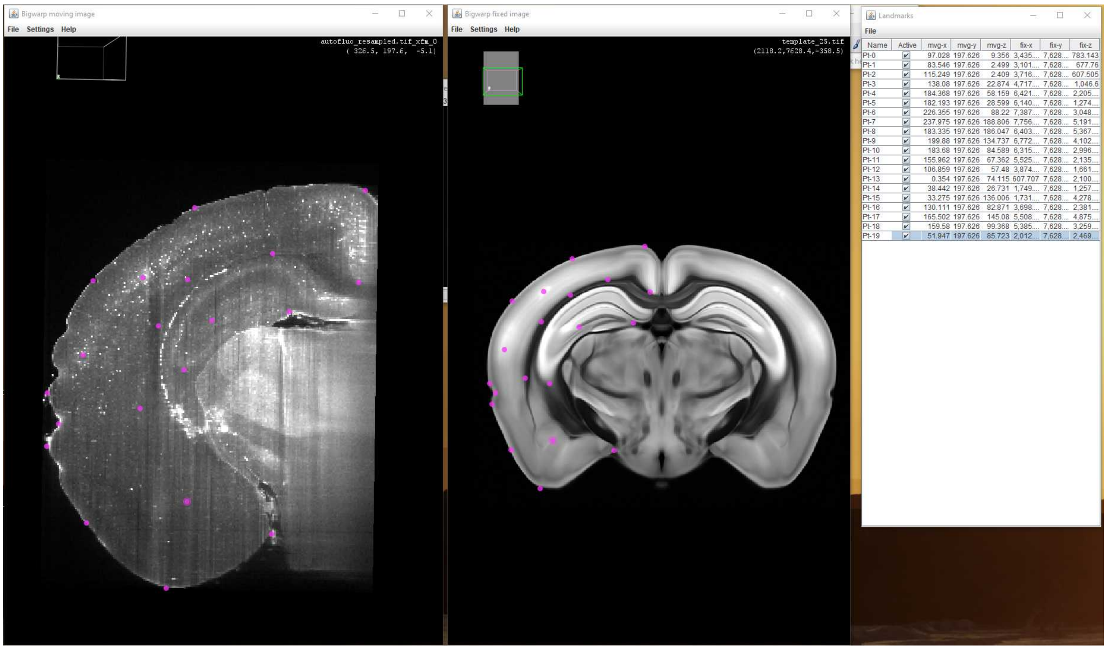

9. Repeat for at least all the locations mentioned under 6)

10. Save file and transform:
- Save the landmarks by clicking File -> Export Landmarks in the landmarks table on the right.
- Open the landmarks.csv file with an editor (e.g. LibreOffice Calc or use the provided python script.
- The first three columns are the point coordinates for the autofluo_resampled, those should be in pixel coordinates and generally not be larger than 400 (depending on stack dimensions). Open a new file, write as follows:

| | A | B | C |
|---|---|---|---|
| | point | | |
| | (number of points, e.g. 70) | | |
| | (x values) | (y values) | (z values) |

- Save as csv with the title "autofluo_landmarks.txt.csv". Importantly, save with spaces as separators instead of commas.
- Close the Calc window, then remove the .csv ending from the file.

- Repeat for the atlas template values, saving in a separate file as "atlas_landmarks.txt.csv" (Note: If the values are much higher (e.g. > 1000), then this is due to ImageJ representing them in micrometers and not in pixel values. To arrive at the correct pixel values, divide by 25 (in case of a 25 µm/voxel template).
- Again, close the Calc window, then remove the .csv ending from the file.

- If you haven't, move both atlas_landmarks.txt and autofluo_landmarks.txt into the brain folder.

11 If you have produced external cell annotations, e.g. via Arivis or as a list of manual coordinates, do the following:

- Make sure your coordinates are in an excel file with the only three columns corresponding to X, Y, and Z coordinates.
- Run a python script like the following (not in the docker container, e.g. use Spyder via Anacona):

Note that this assumes that the excel does not contain a header (x,y,z,... in the first line). If it does, then remove them from the file before proceeding.

```
# -*- coding: utf-8 -*-
import pandas as pd
import numpy as np

#load the file
coordinates = pd.read_excel("path_to_excel_file")

#save as numpy
coordinates_np = coordinates.to_numpy()
coordinates.astype(int)
np.save(coordinates_np,"brain_folder_path/cells.npy)
```

12 Copy the 02_process_Ilastik_run_this.py into the folder and **rename into process_Ilastik_run_this.py** (overwrite the previous file of that name)

The first block will align (register) the two downsampled stacks onto each other, which in our experience is not necessary. If you are using only a single channel as source, then it is definitely not necessary and shouid be commented out.
Subsequently, the downsampled stack is registered to the atlas using Elastix. Unless specified otherwise, Elastix will use the landmark files you provided as a result of step 10.

```
#Alignment operations:
######################
#correction between channels:
#resultDirectory  = alignData(**CorrectionAlignmentParameter);

#alignment to the Atlas:
resultDirectory  = alignData(**RegistrationAlignmentParameter);
```

12.1 Next, the cells are detected, either using Ilastik (default) or the built-in thresholding approach of Clearmap1. This should result in a cells.npy file being saved in the same directory.

```
#Cell detection:
################
detectCells(**ImageProcessingParameter);
```

**Alternatively, if you generated the cells externally, skip this step (comment out ).** Make sure you generated a "cells.npy" file from your external data and placed in the folder instead.

12.2 Subsequently, the detected cells willl be loaded.

```
#Filtering of the detected peaks:
#################################
#Loading the results:
points, intensities =
io.readPoints(ImageProcessingParameter["sink"]);

#Thresholding: the threshold parameter is either intensity or
size in voxel, depending on the chosen "row"
#row = (0,0) : peak intensity from the raw data
#row = (1,1) : peak intensity from the DoG filtered data
#row = (2,2) : peak intensity from the background subtracted
data
#row = (3,3) : voxel size from the watershed
points, intensities = thresholdPoints(points, intensities,
threshold = 20, row = 1);
io.writePoints(FilteredCellsFile, (points, intensities));
```

Then, Elastix will be used to transform the points, first for the inter-channel correction alignment, and then for the atlas registration. By default it is assumed that the inter-channel alignment is skipped, so only uncomment the three steps if you are actually using it.

```
# Transform point coordinates
#############################
points =
io.readPoints(CorrectionResamplingPointsParameter["pointSource"]
);
#points = resamplePoints(**CorrectionResamplingPointsParameter);
#points = transformPoints(points, transformDirectory =
CorrectionAlignmentParameter["resultDirectory"], indices =
False, resultDirectory = None);
#CorrectionResamplingPointsInverseParameter["pointSource"] =
points;
points =
resamplePointsInverse(**CorrectionResamplingPointsInverseParamet
er);
RegistrationResamplingPointParameter["pointSource"] = points;
points = resamplePoints(**RegistrationResamplingPointParameter);
points = transformPoints(points, transformDirectory =
RegistrationAlignmentParameter["resultDirectory"], indices =
False, resultDirectory = None);
io.writePoints(TransformedCellsFile, points);
```

Subsequently, generate heatmaps. The intensity heatmaps are mostly interesting if you are using the built-in Clearmap1 built-in processing. If using Ilastik (default) or external coordinates, then it is commented out.

```
# Heat map generation
#####################
points = io.readPoints(TransformedCellsFile)
#intensities = io.readPoints(FilteredCellsFile[1])

#Without weigths:
vox = voxelize(points, AtlasFile, **voxelizeParameter);
io.writeData(os.path.join(BaseDirectory, 'cells_heatmap.tif'),
vox.astype('int32'));

#With weigths from the intensity file (here raw intensity):
#voxelizeParameter["weights"] = intensities[:,0].astype(float);
#vox = voxelize(points, AtlasFile, **voxelizeParameter);
#io.writeData(os.path.join(BaseDirectory,
'cells_heatmap_weighted.tif'), #vox.astype('int32'));
```

Lastly, the overview tables are generated. Again, the intensities-adjusted table is mostly relevant for the CM1 built-in cell segmentation, and is deactivated by default. If you are using the CCF3 reference atlas, be aware that there are some regions in the annotation file that do not have a matching Region_IDs.csv entry. If you are using the CCF3, make sure to uncomment the line correcting the annotation values.

```
#Table generation:
##################
'''
#With integrated weigths from the intensity file (here raw
intensity):
ids, counts = countPointsInRegions(points, labeledImage =
LabeledImage, annotations = AnnotationFile, intensities =
intensities, intensityRow = 0);
table = numpy.zeros(ids.shape, dtype=[('id','int64'),
('counts','f8'),('name', 'a256')])

#Optional: if no collapse is used (collapse = None, or left out
entirely)
#ids = [0 if (x==1216) or (x==2656) or (x==12096) else x for x
in ids]

table["id"] = ids;
table["counts"] = counts;
table["name"] = labelToName(ids);
io.writeTable(os.path.join(BaseDirectory,
'Annotated_counts_intensities.csv'), table);
'''

#Without weigths (pure cell number):
ids, counts = countPointsInRegions(points, labeledImage =
LabeledImage, annotations = AnnotationFile, intensities = None);
table = numpy.zeros(ids.shape, dtype=[('id','int64'),
('counts','f8'),('name', 'a256')])

#Optional: if no collapse is used (collapse = None, or left out
entirely)
#ids = [0 if (x==1216) or (x==2656) or (x==12096) else x for x
in ids]

table["id"] = ids;
table["counts"] = counts;
table["name"] = labelToName(ids);
io.writeTable(os.path.join(BaseDirectory,
'Annotated_counts.csv'), table);
```

13 Subsequently, run the Docker container for the second time. If everything goes according to plan, this should generate a table for the cells per region. The Docker container should finish after running.

**On Windows**, open power shell, navigate to the folder with the process_Ilastik_run_this.py script and adapt the following code:

```
docker run -it --rm -v C:\path\to\brain:/CloudMap/Data
-v C:\path\to\custom\Ilastik\file:/CloudMap/classifiers
friendly_clearmap:3.0 ;
```

Note that file paths must not end on a trailing backslash.

**On Linux**, open a terminal, then navigate to the folder with the process_Ilastik_run_this.py script and run the following command:

```
docker run -it --rm -v /path/to/brain/:/CloudMap/Data
-v /path/to/custom/Iastik/file/:/CloudMap/classifiers
friendly_clearmap:3.0
```

**On MacOS**, open a terminal (Spotlight -> press Space -> type " Terminal"), then navigate to the folder with the process_Ilastik_run_this.py script and type the folllwing command:

```
docker run -it --rm -v /path/to/brain/:/CloudMap/Data
-v /path/to/custom/Iastik/file/:/CloudMap/classifiers
friendly_clearmap:3.0
```

By default, the container will execute the process_Ilastik_run_this.py script. If you want to run a different script, you can add the option

```
--entrypoint "python3 /CloudMap/Data/your_custom_script.py"
```

prior to the friendly_clearmap:3.0 argument.
Note that in this case the script needs to be in the folder that is mapped to /CloudMap/Data with the first -v command (i.e. the folder containing all image data).

#### 13.1 Hooray! If everything worked, you should have:

- an atlas-aligned set of points, both in .npy form (cells_aligned.npy)
- a region table (Annotated_counts.csv) with cell counts per region.
- A heatmap with cell densities in the same space as your atlas.
