## Supplementary Protocol 2: Docker for Clearmap 2 for "FriendlyClearMap: An optimized toolkit for mouse brain mapping and analysis"

CN54VG8W

**Protocol Info:** Moritz Negwer . Run Clearmap 2 docker. **protocols.io** https://protocols.io/view/run-clearmap-2-docker-cn54vg8wVersion created by Moritz Negwer

**Created:** Feb 09, 2023

**Last Modified:** Feb 15, 2023

**PROTOCOL integer ID:** 76700

### Run Clearmap 2 docker V.(cn54vg8w)

Moritz Negwer[1]

[1]Radboudumc Nijmegen

M Moritz Negwer

DISCLAIMER

Dropbox links are temporary and will be replaced eventually.

ABSTRACT

This protocol is a supplement to our upcoming publication "FriendlyClearMap: An optimized toolkit for mouse brain mapping and analysis".

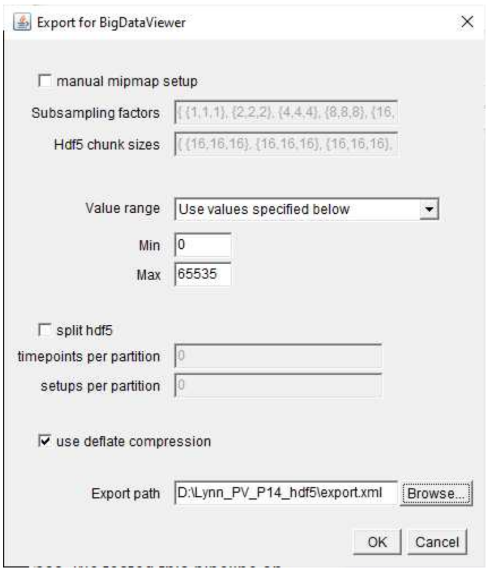


FIJI will then generate the image pyramid for the hdf5 stack and save it in the appointed location.

4 **Loading the data in Ilastik**

Generate a Pixel Classification workflow in Ilastik

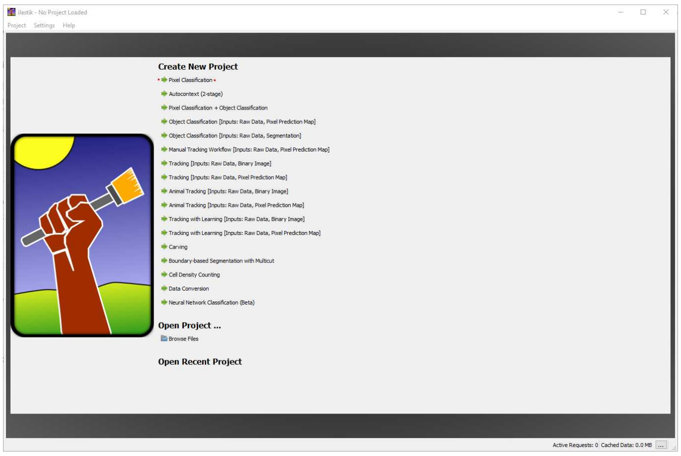


Load the hdf5 file via "Add Single Image", then select t0000/s00/1/cells to display the

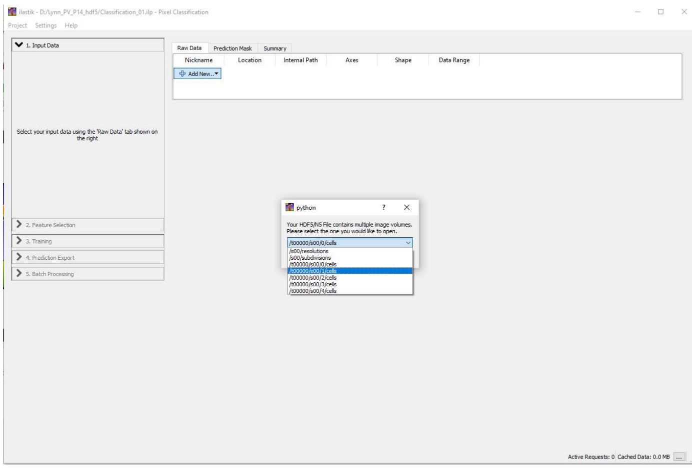

Copy the stack into the project file by right-clicking on the image -> "Edit", then selecting "Copy into project file". This results in larger Ilastik files, but is important for loading the same files in the docker environment (relative paths will not be preserved).

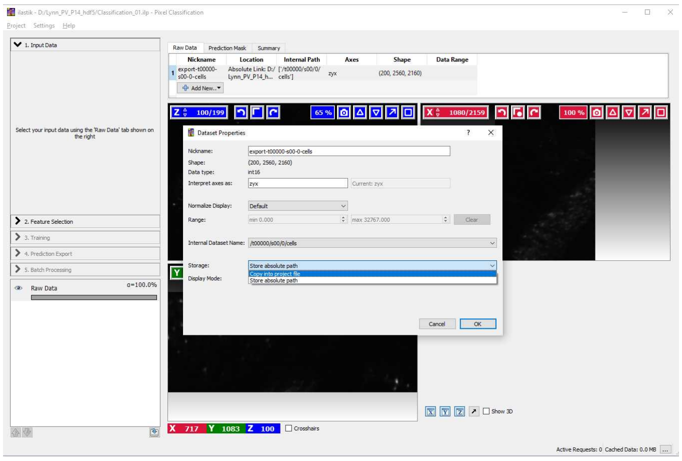

If you want to adjust the brightness of the displayed images, you can do so by right-clicking on "Raw Data" in the left column -> "Adjust Thresholds". Then uncheck "Automatic Range" and set the maximum value so that your cells of interest are visible.

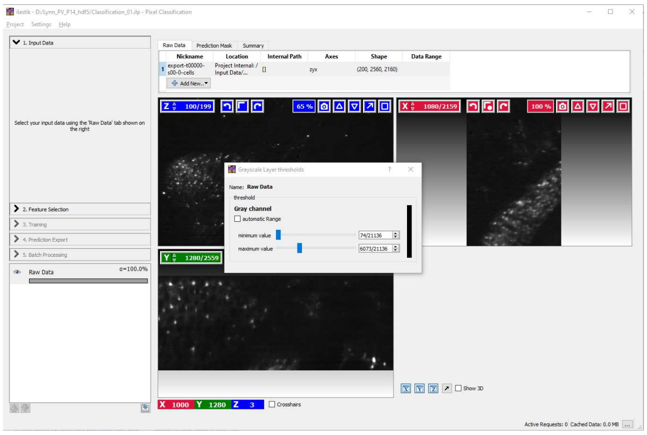

#### 5 Segmentation with Ilastik

In the "Select Features" pane, add a 30 (pixels wide) filter by clicking to the right of the 10 column and entering 30.
Then, select the "Edge" and "Texture" boxes for all σ (filters).

(Note that those are empirically determined settings. Adding a 30px filter approximately captures single neurons when imaged at 1.1x magnification on a 2160 x 2560 pixel camera. Different imaging conditions, notably with different magnifications, might require different-size filters. Also, skipping the intensity-based filters speeds up calculations without noticeable worse segmentation quality.)

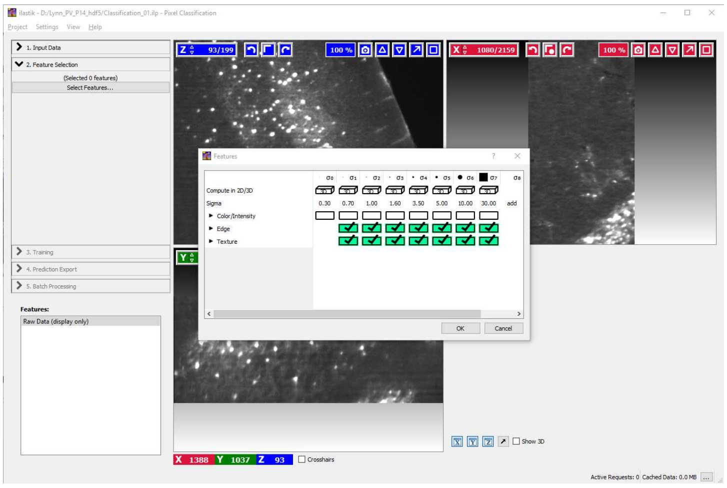

5.1 In the "Training" pane, define the background (Label 2)

- Zoom out in the XY projection (Ctrl + mousewheel) until the entire brain is visible
- Select Label 2, set to max size (61) and draw along the empty space besides the brain. Take

care not to draw over the edge of the image, as that can lead to problems down the road with some Ilastik versions.
- Subsequently, draw single dots along the edges of the brain and in the neuropil where there are no labelled cells.

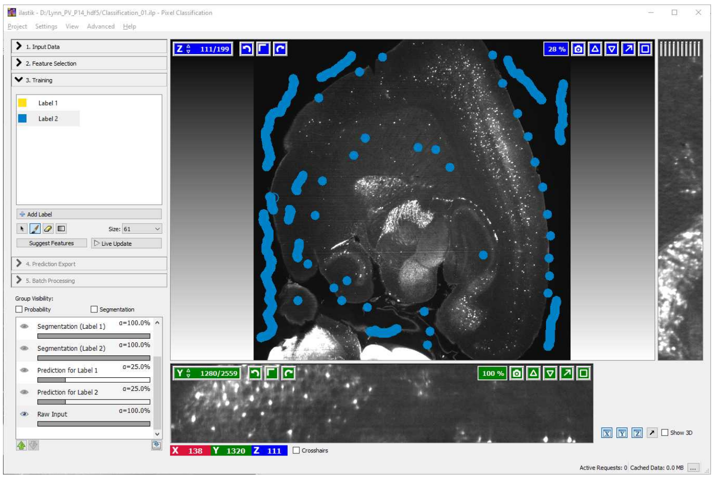

5.2 Subsequently, zoom in on your signal (Ctrl + Scroll wheel). To syncronze the view between the XY projection and the other two, double-leftclick on an object of interest, e.g. a cell. The voxel you clicked will then be centered in the other projections as well.

1) Select a few cells as signal (Label 1, yellow). For our images, a size 5 brush covered most of the labelled cell bodies at their maximum extent.

2) Surround the selected cell with a thin circle of background (1-3 px wide). We found this to be especially helpful when marking cells close to each other, as otherwise Ilastik may wrongly mark the region in between also as signal.

3) Repeat for the same cells for the XZ and YZ projections.

4) We recommend marking ca. 100 cells in total, spread out over the stack. Take care to select regions that are representative of the entire brain, i.e. with different densities and background intensities.

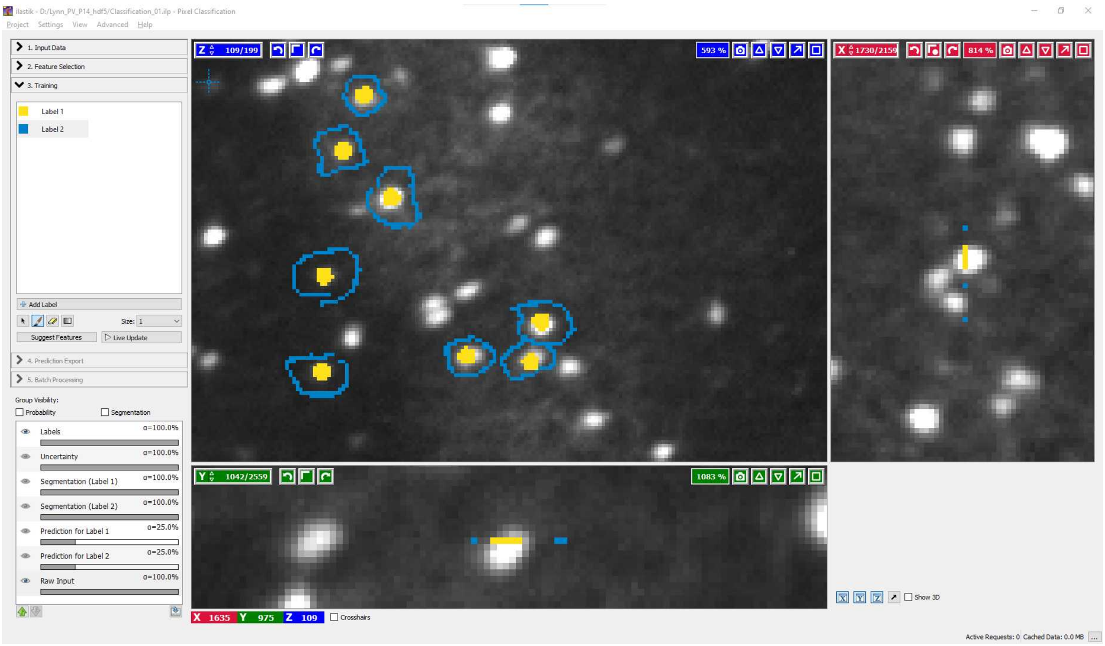


5.3 Once you have marked ca. 100 cells, zoom in on one region in all three projections and click "Live Preview" to trigger the training of the random forest algorithm with your input data.

**Note:** Do *not* add new input data during training, as this will trigger a re-training from the start.

Depending on the number and speed of your CPU cores, this may take a long time (10-45 minutes) to complete. Once it is completed, the prediction will be overlaid over your field of view.

Once the training has finished, verify that the segmentation is correct. If not and more training data needs to be added, **turn off "Live Preview" before adding new data** - otherwise, every new addition will trigger a fresh round of training, leading to long delays.

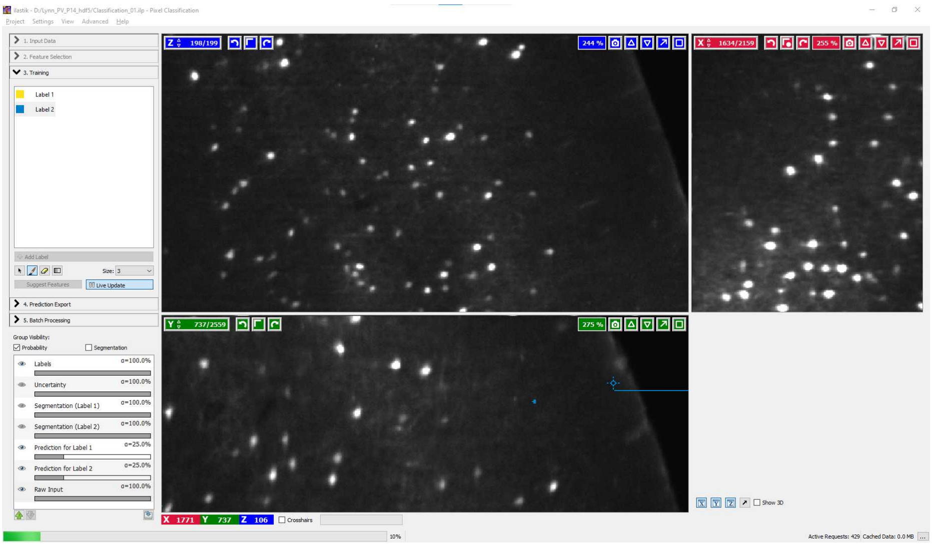

Training in progress. Note the progress bar in the bottom left corner.

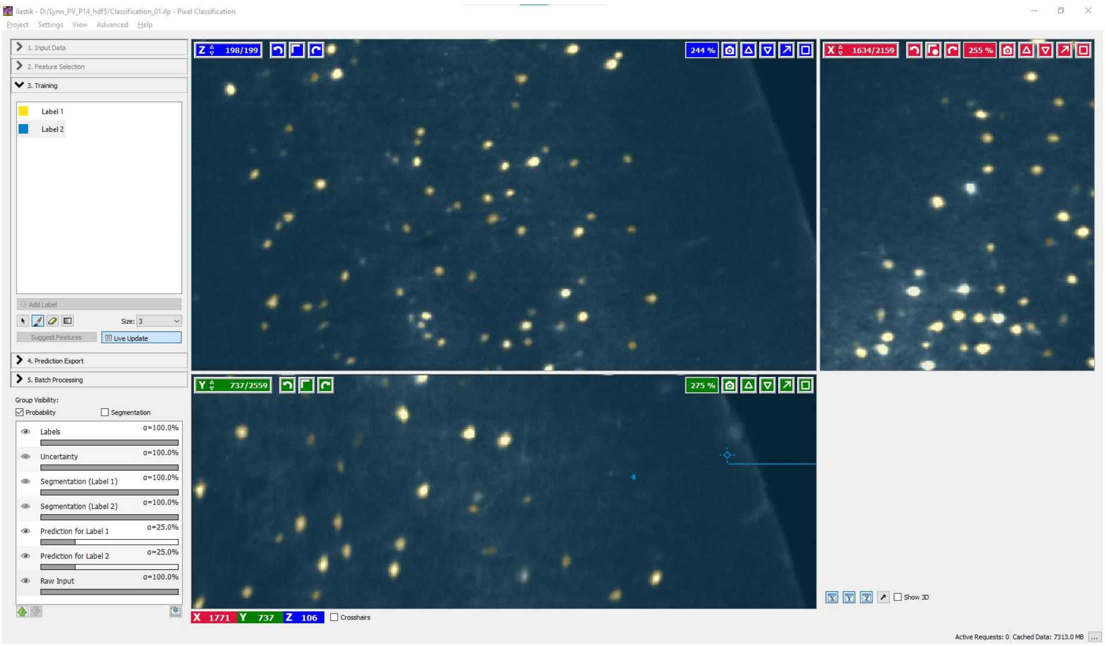

Training finished. Note the cell in the center of the XZ projection is predicted to be background. However, scrolling through the XZ projection will reveal that the following plane is segmented as a cell. Therefore, no action is needed.

#### 5.4 Auto-selecting the best features for training (optional)

Ilastik offers a "Suggest Features" option. After successful training, this option compares subsets of features to find which reduced subset still represents the training data.

Doing so can take a long time (several hours), but once done, it reduces the set of required features from the 14 selected in Step 5 of this example to a smaller amount (default 7).

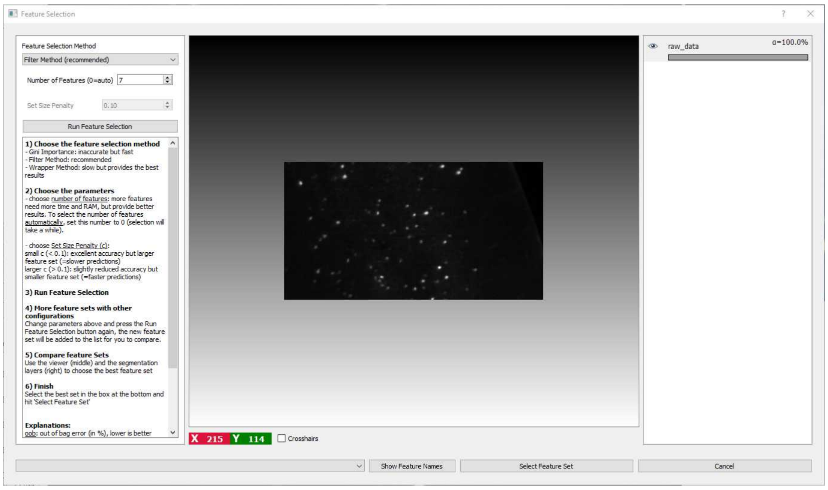

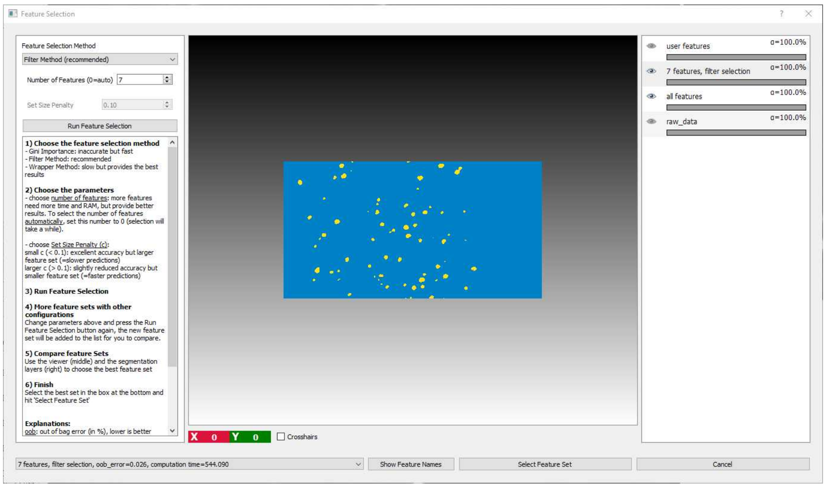

Compare the feature by clicking on the eye symbols in the left column. If you agree with the selected subset ("7 features"), change the features in the bottom right drop-down menu. There, you can also see the computation time as a comparison. In our example case, the auto-selected subset is approx. 40% faster than the full set of features.

Finally, click "Select Feature Set" to confirm. Then, retrain the random forest algorithm with the new subset of parameters by clicking "Live Preview" once more.

#### 5.5 Setting the Ilastik Export settings correctly

Once the training is complete, proceed to the "Prediction Export" pane. There, click "Choose Export Image Settings", and set the following parameters:

- Change "Convert to data type" to "unsigned 8-bit"
- Check "Renormalize [min,max] from 0.0 / 1.0 to 0 / 255

The rest can be left at default. For the final settings, see below:

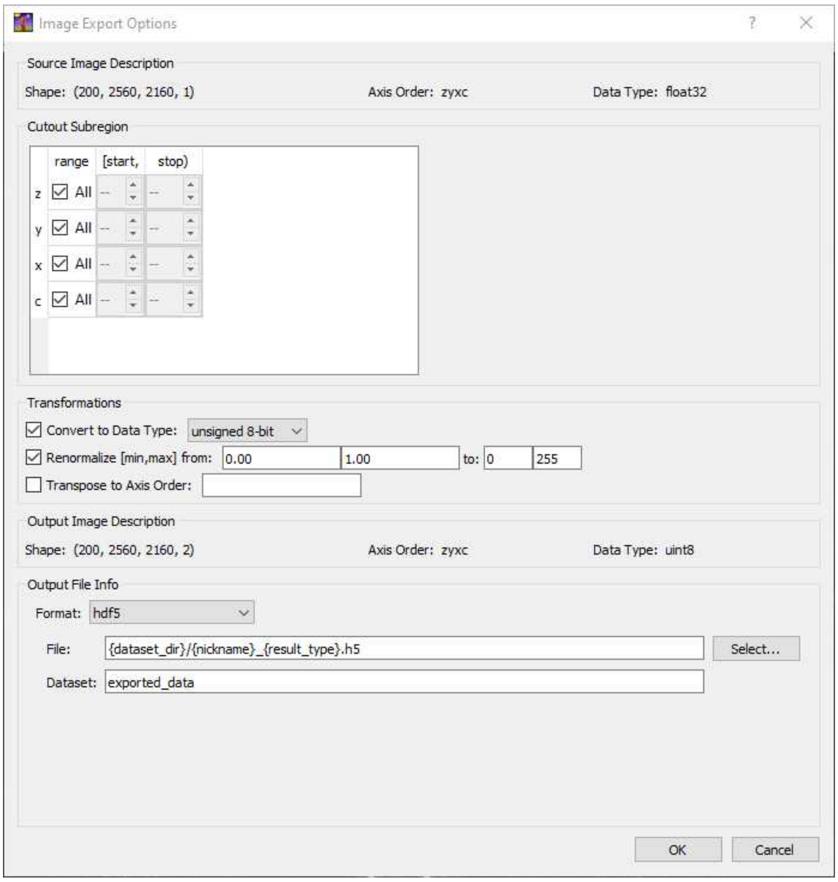


Confirm with OK. Then save (File -> Save), and close the project. It should now be ready for use in both Clearmap1 and Clearmap 2.

#### Docker Setup

6 If you have an install of Arivis Vision4D with the Machine Learning Segmenter package, you can proceed as follows to generate a finished cell coordinate file.

6.1 Convert your dataset with the Arivis SIS converter

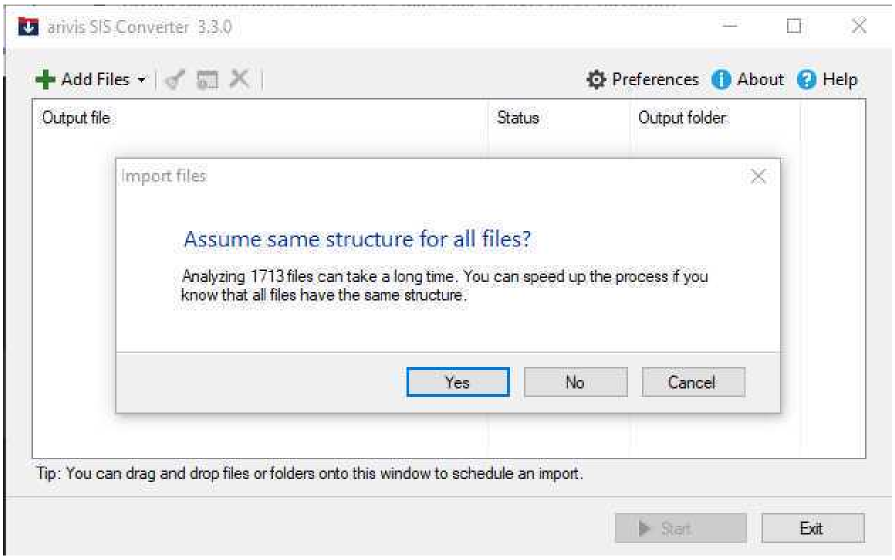


Load all files from the image stack as z-planes into SIS converter

6.2 Then, load them into Arivis, and in the left Processing Pane, add a "Machine Learning Segmenter" processing step. No pre-processing should be necessary.

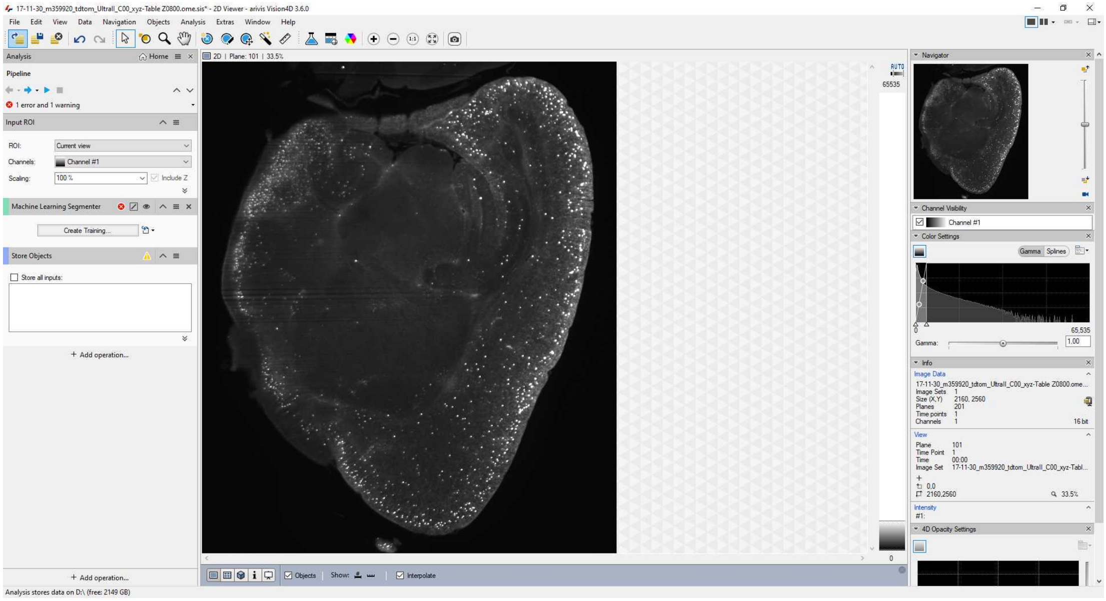

6.3

Subsequently, open the "Train Machine Learning Segmenter" pane and start adding foreground and background labels. We recommend the following:
- Make sure to assign some background label to the blank space around the brain and the edges of the sample.
- Mark a representative subset of cells (we recommend 100 throughout the brain).

- Mark a small circle of background around the marked cells. In our experience, this helps considerably with separation of dense cells.
- Make sure to mark dense projections, fiber tracts, membranes, blood vessels, etc. as background.

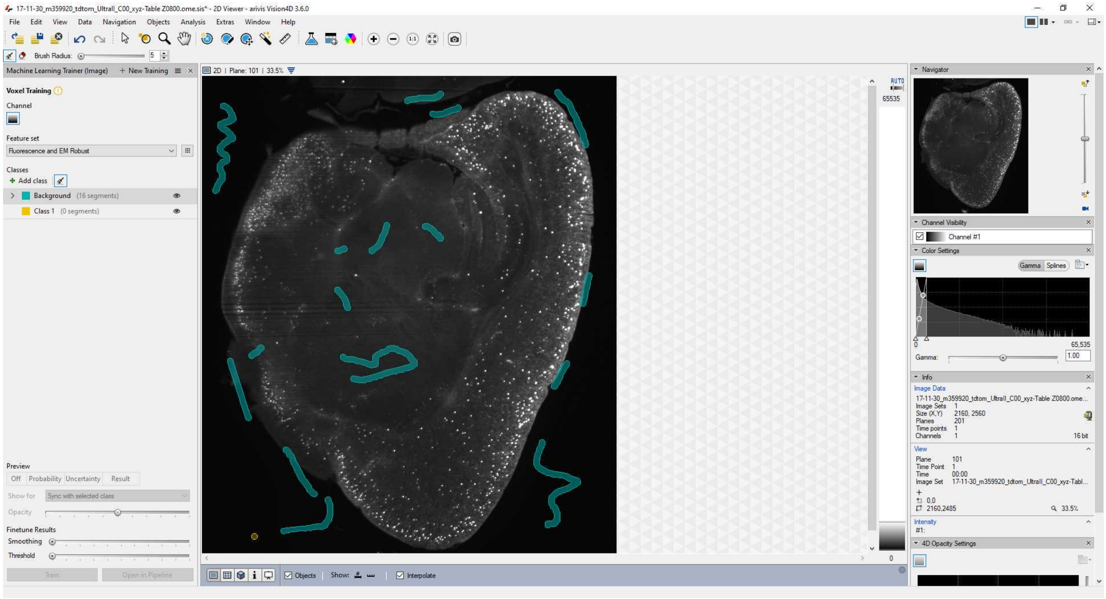


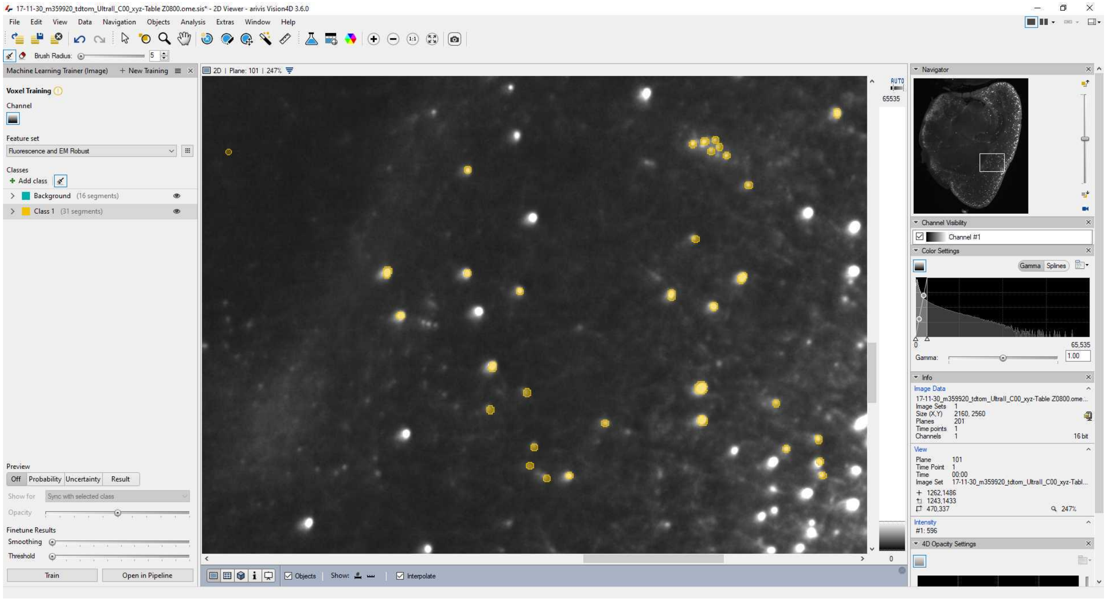

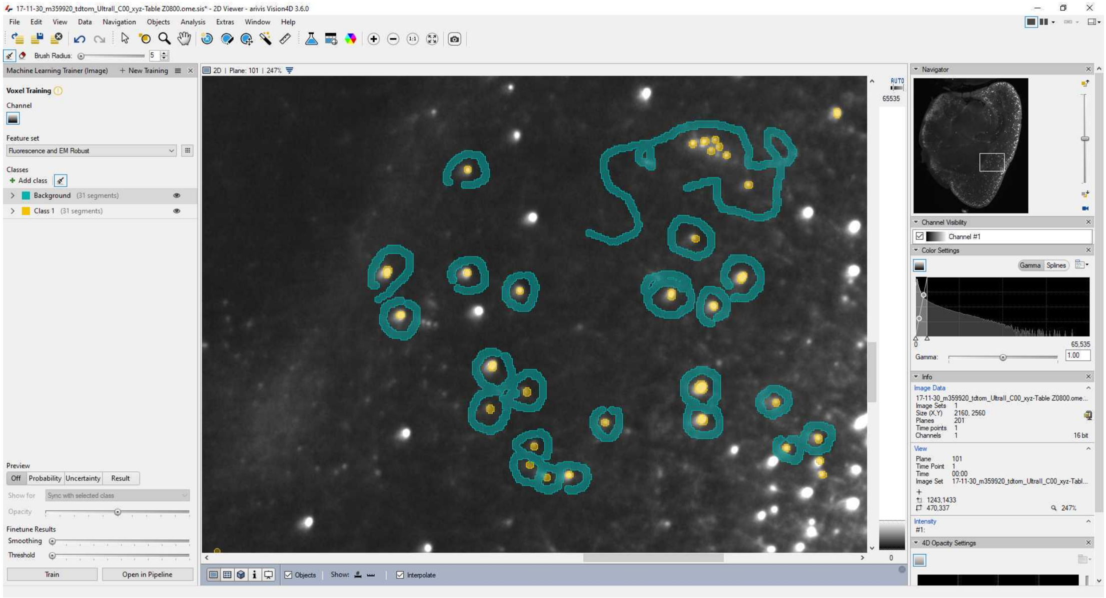

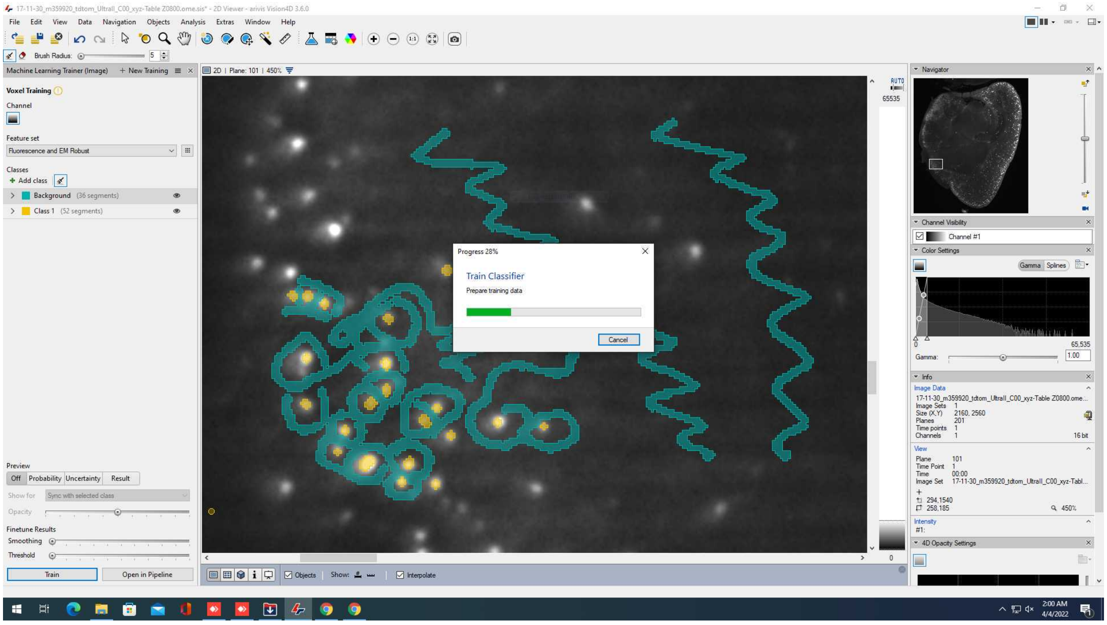


(If you already have a trained Ilastik file, you can also import it via the "Import from Ilastik" option).

Once trained, save the training so it can be reused.

6.4 In the pipeline pane on the left, make sure to change the Input ROI to "current image set" (otherwise it will remain at the default "current view", i.e. the 2D cutout that you are viewing.

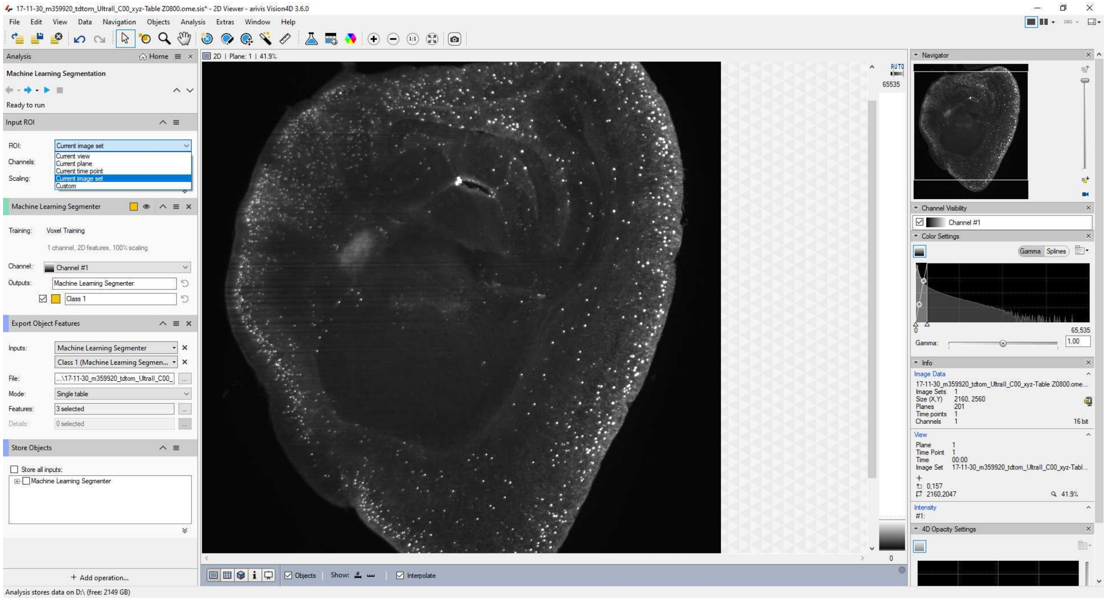

To export as a coordinate list, add a "Export Objects Features" step to the processing pipeline.

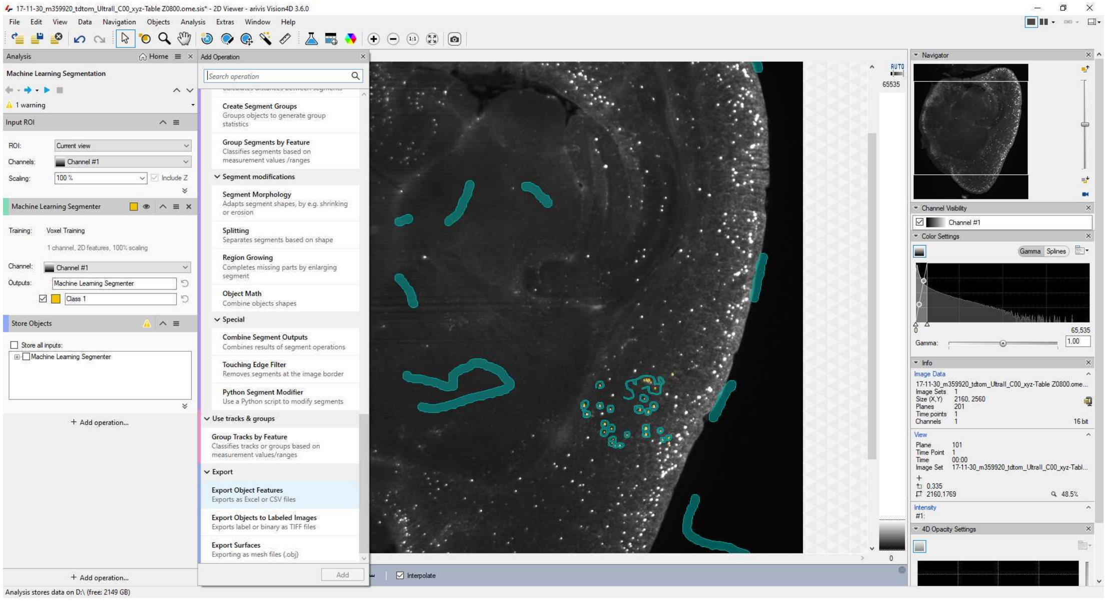

Specify the output as an Excel file. Select the following Object Properties:
- Deselect everything (defaults won't make much sense)
- Center of Gravity X / Y / Z (px)

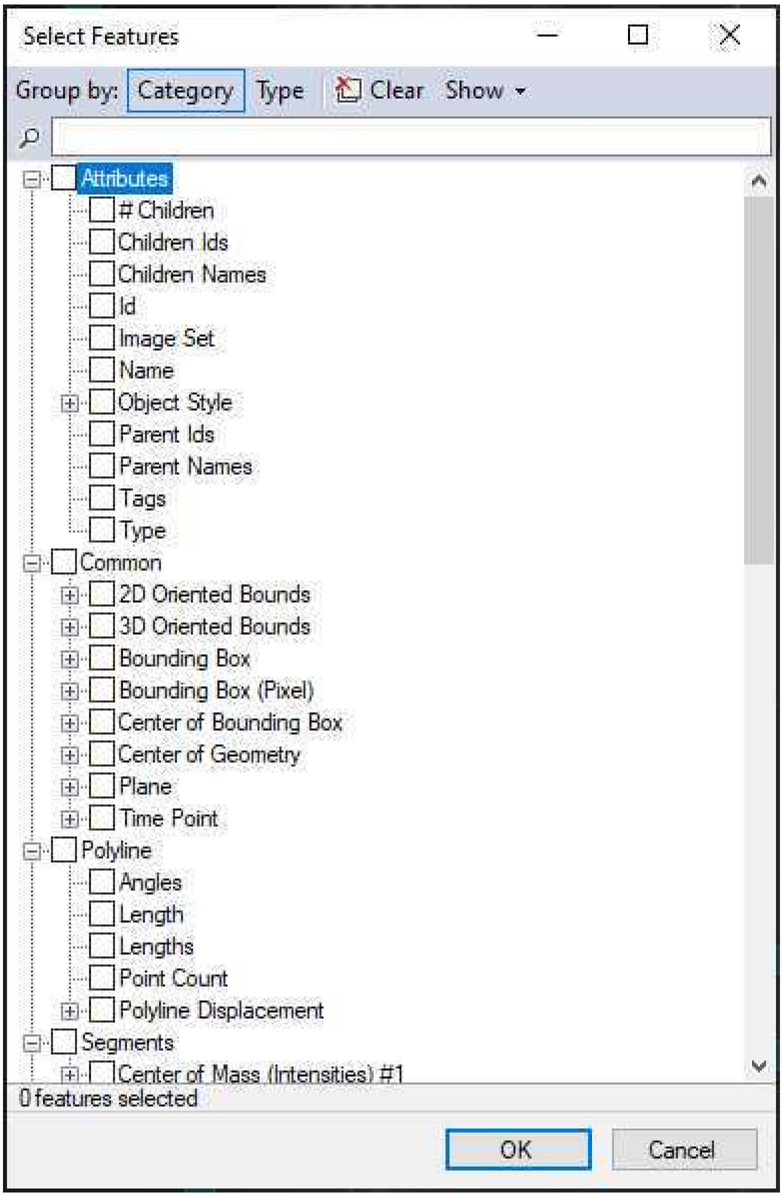
Select Features
Group by: Category Type Clear Show
Attributes
### Children
Children Ids
Children Names
Id
Image Set
Name
Object Style
Parent Ids
Parent Names
Tags
Type
Common
2D Oriented Bounds
3D Oriented Bounds
Bounding Box
Bounding Box (Pixel)
Center of Bounding Box
Center of Geometry
Plane
Time Point
Polyline
Angles
Length
Lengths
Point Count
Polyline Displacement
Segments
Center of Mass (Intensities) #1
0 features selected
OK
Cancel

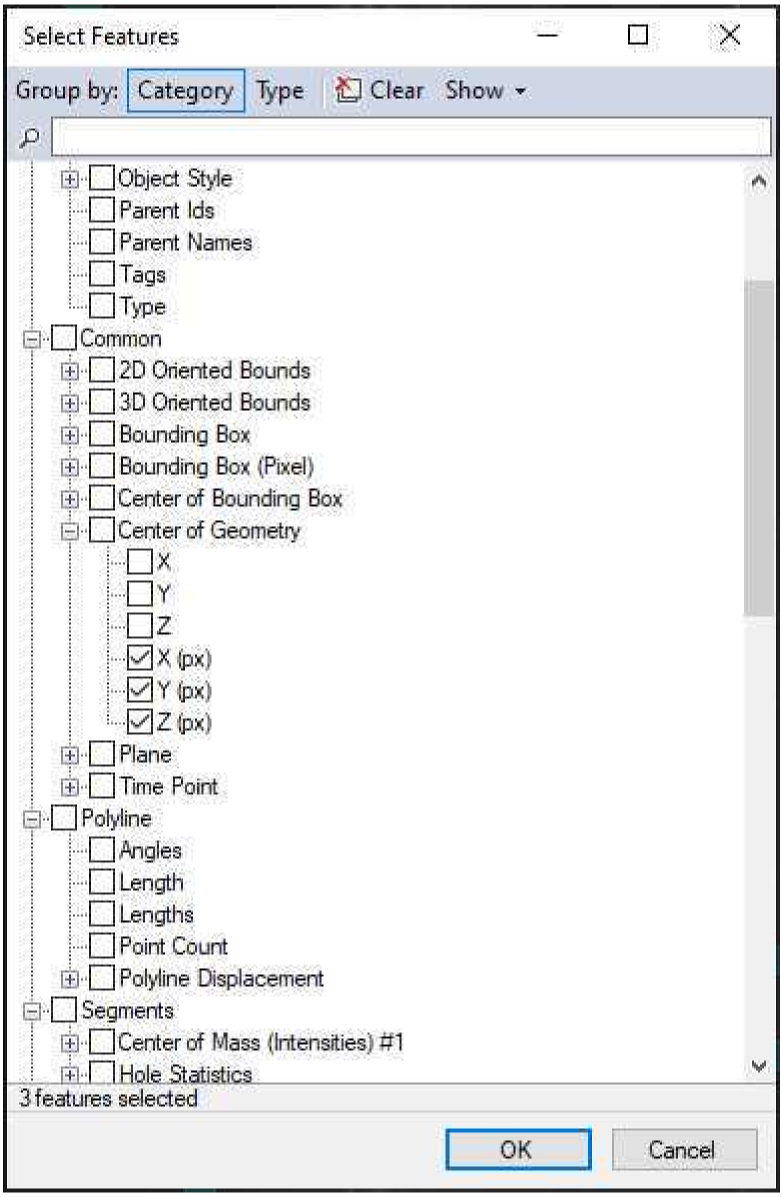


**6.5** Save the pipeline.

Then, run it for the file at hand. The Machine Learning Segmenter plugin is CPU-heavy and will likely take approx. 24h for a 2160 x 2560 x 2000 pixel stack on a recent (14 core, 128GB RAM, fast NVMe SSD) workstation.

8.1 Copy the process_Ilastik_run_this.py into the folder and adapt as follows:

```
expression_raw = 'name_of_your_autofluo_tif_file Z<Z,4>.ome.tif'
```

In this case, the Z<Z,4> signals that there are four numbers after the Z , starting from 0000.

```
expression_auto = expression_raw #if only one channel is present

#expression_auto = 'name_of_your_signal_tif_file Z<Z,4>.ome.tif'
```

If only one channel is used (e.g. if you used a 94 fluorophore and have plenty of autofluorescent background), then set expression_auto = expression raw. Otherwise uncomment the next line with the file name of your signal (e.g. 647 channel) image tiffs.

Subsequently, adapt the atlas properties (line 78+)

```
  #init atlas and reference files

  annotation_file, reference_file,
distance_file=ano.prepare_annotation_files(
      slicing=(slice(None),slice(None),slice(0,256)),
      orientation=(1,-2,3),
      overwrite=False, verbose=True);

  #alignment parameter files
  align_channels_affine_file   = io.join(resources_directory,
'Alignment/align_affine.txt')
  align_reference_affine_file  = io.join(resources_directory,
'Alignment/align_affine.txt')
  align_reference_bspline_file = io.join(resources_directory,
'Alignment/align_bspline.txt')
```

Leave this block uncommented if you want to use the built-in CCF3 atlas of clearmap2. Otherwise block-comment it ("" at beginning and end).

In case you want to use your own atlas, uncomment the block and adapt the following:

```
  #in case you want to use your own atlas:
  atlas_dir = '/CloudMap/Data/atlas/P56_CCF2/'
  atlas_file = io.join(atlas_dir, 'template_25.tif')
  atlas_annotation_file =
io.join(atlas_dir,'annotation_25_full.nrrd')

  #custom atlas
  annotation_file, reference_file, distance_file =
ano.prepare_annotation_files(
      slicing = None,
      directory = atlas_dir,
      annotation_file = atlas_annotation_file,
      reference_file = atlas_file,
      distance_to_surface_file = None,
      orientation = (1,2,3),
      overwrite=False,
      verbose=True)

  align_channels_affine_file   =
io.join(atlas_dir,'Par0000affine_acquisition.txt')
  align_reference_affine_file  =
io.join(atlas_dir,'Par0000affine_acquisition.txt')
  align_reference_bspline_file =
io.join(atlas_dir,'Par0000bspline.txt')
```

Also, uncomment the and adapt custom annotations in line 414:

```
# only for atlases other than standard atlas
  custom_label_csv =
pandas.read_csv('/CloudMap/Data/atlas/P56_CCF2/regions_IDs.csv')
```

**Note** this means the atlas folder needs to be placed in the atlas folder (which in the Docker container will be represented as /CloudMap/Data/)

Then adapt the follwing parameters (line 139+)

```
#%% Resample

  resample_parameter = {
      "source_resolution" : (3.25, 3.25, 3),
      "sink_resolution"   : (25,25,25),
      "processes" : None,
      "verbose" : False,
      "orientation" : (1,2,3)
      };
```

Adapt this to the dimensions of your scan.
In the case of LaVision / Miltenyi Ultramicroscopes, the tiff headers should contain a large amount of information. You can find this information by loading the first tiff file in FIJI and clicking Image -> Show Info. In that case, save the content of the "File Info" window (File -> Save As, then choose a name and save as .txt).
Then re-open the resulting .txt file with a text editor such as Notepad++ (https://notepad-plus-plus.org/downloads/) on Windows or gedit on Linux. Search for "PhysicalDimension" and note the X, Y, and Z values.

If you are using a custom reference atlas with a different resolution (e.g. 10 x 10 x 10 µm/voxel) , adapt the sink_resolution values.

Finally, set the action you want to take (line 58):

```
#TODO: Specifiy which action you want to take. Uncomment what
you want to do
  Action = 'Preprocess' #only generate scaled-down resamples of
expression + autoflu, if you want to manually align to the atlas
```

#### Atlas alignment with Dockerized Clearmap pipelines

9 Subsequently, run the Docker container for the first time. If everything goes according to plan, this will generate three additional tiffstacks downsampled to 25µm/voxel, suitable for alignment to the Allen Brain Atlas. The Docker container should finish after running.

**On Windows**, open power shell, navigate to the folder with the process_Ilastik_run_this.py script and adapt the following code:

```
docker run -it --rm -v C:\path\to\brain:/CloudMap/Data
-v C:\path\to\custom\Ilastik\file:/CloudMap/classifiers
friendlyclearmap:4.2 ;
```

Note that file paths must not end on a trailing backslash.

**On Linux**, open a terminal, then navigate to the folder with the process_Ilastik_run_this.py script and run the following command:

```
docker run -it --rm -v /path/to/brain/:/CloudMap/Data
-v /path/to/custom/Iastik/file/:/CloudMap/classifiers
friendlyclearmap:4.2
```

**On MacOS**, open a terminal (Spotlight -> press Space -> type " Terminal"), then navigate to the folder with the process_Ilastik_run_this.py script and type the folllwing command:

```
docker run -it --rm -v /path/to/brain/:/CloudMap/Data
-v /path/to/custom/Iastik/file/:/CloudMap/classifiers
friendlyclearmap:4.2
```

By default, the container will execute the process_Ilastik_run_this.py script. If you want to run a different script, you can add the option

```
--entrypoint "conda -n cm2 /bin/bash python3
/CloudMap/Data/your_custom_script.py"
```

prior to the friendlyclearmap:4.2 argument.
Note that in this case the script needs to be in the folder that is mapped to /CloudMap/Data with the first -v command (i.e. the folder containing all image data).

#### 10 Atlas alignment using BigWarp

(In Windows/Linux, the Docker container should have finished)

1. In FIJI/Imagej. open both the atlas stack (e.g. template_25.tif) and the autofluo_resampled.tif files.

2. Open BigWarp (Plugins -> BigDataViewer -> BigWarp)

3. With BigWarp, open the atlas template as moving image, and autofluo_resampled as target image:

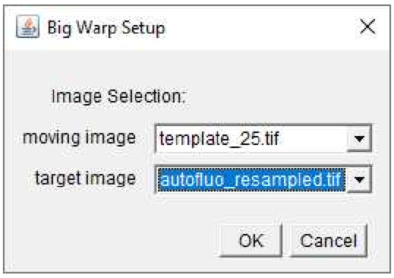


4. You should now see your autofluo_resampled on the left, and the atlas template on the right.

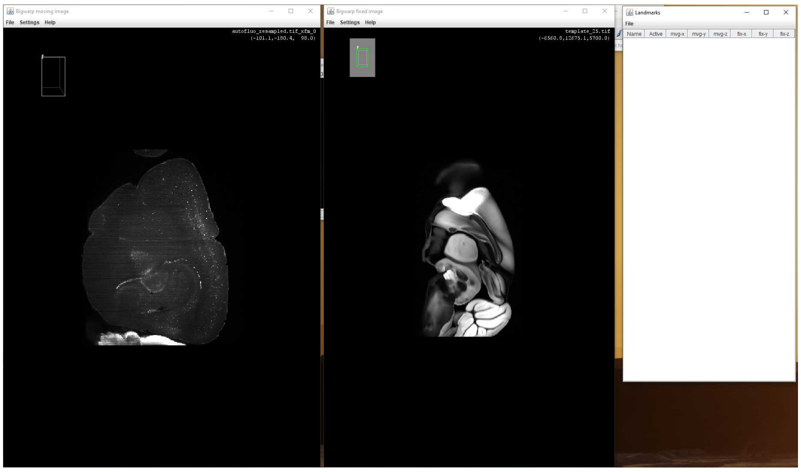

5. Go to the sagittal plane in both:
- Ctrl+A should change to a ZX projection
- Press Z to fix rotation around the Z (originally Y) axis
- Use the arrow keys to rotate around the axis until the orientations are roughly similar

6. Use the mosue wheel to scroll to a similar location in both images. Suggested:
  - Frontal Pole
  - Anterior Commissure crossing over
  - Maximum extent of the Somatosensory Cortex Barrel Field
  - Hippocampal landmarks, e.g. just before Fornix projects entirely ventrally
  - Posterior pole of V1
  - Cerbellar regions (if applicable)

7. Fine-tune the autofluo_resampled projection to match the coronal plane of the atlas template:
- Left-click on the point in the autofluo_resampled image that you would like to use as center of rotation
- Carefully move the mouse to rotate the cutting plane in 3D space. Repeat until it matches
- To reset, press Ctrl+A, then repeat step 5.

8. Generate landmarks:
- Press spacebar to start landmark mode.
- Identify corresponding points.
- Mark one point in autofluo_resampled, *then immediately* mark the corresponding point in the atlas template.
(Note that clicking several points in the same image will lead to non-matched point sets, which will not work for alignment later).
- Mark between 5 and 20 point pairs per location.
- Take care to mark a diversity of matched locations (e.g. pial surface, white matter, layer 4, hippcoampal landmarks)
- To change perspective (or undo landmarks), disengage landmark mode by pressing spacebar again. Re-engage once you are happy with the result.

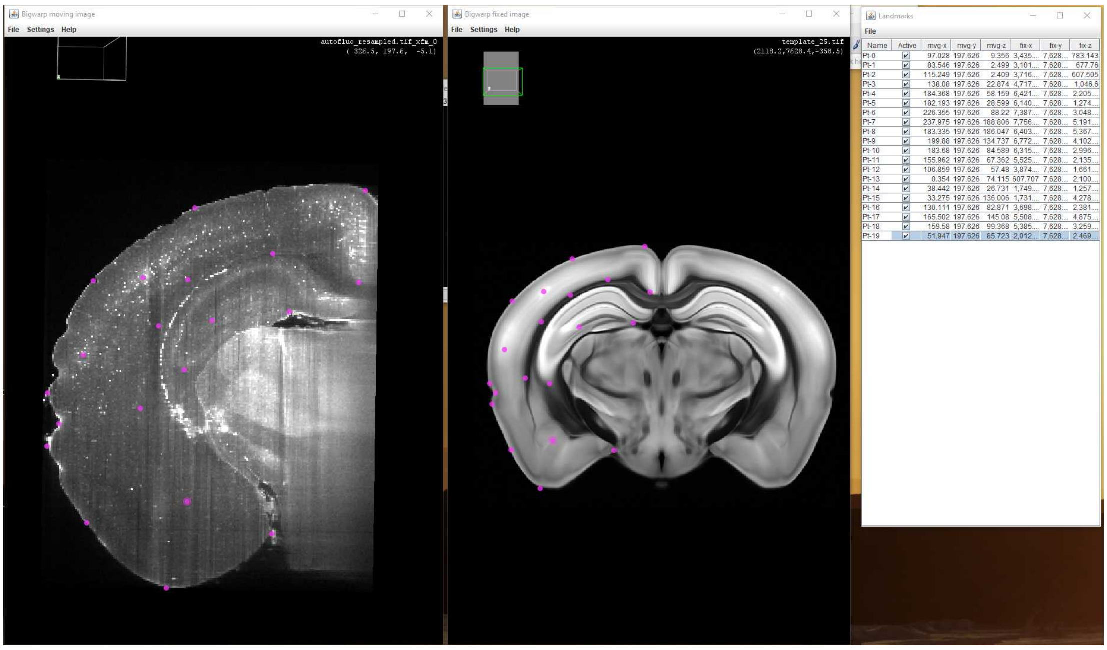


9. Repeat for at least all the locations mentioned under 6)

10. Save file and transform:
- Save the landmarks by clicking File -> Export Landmarks in the landmarks table on the right.
- Open the landmarks.csv file with an editor (e.g. LibreOffice Calc or use the provided python script.
- The first three columns are the point coordinates for the autofluo_resampled, those should be in pixel coordinates and generally not be larger than 400 (depending on stack dimensions). Open a new file, write as follows:

| | A | B | C |
|---|---|---|---|
| | point | | |
| | (number of points, e.g. 70) | | |
| | (x values) | (y values) | (z values) |

- Save as csv with the title "autofluo_landmarks.txt.csv". Importantly, save with spaces as separators instead of commas.
- Close the Calc window, then remove the .csv ending from the file.

- Repeat for the atlas template values, saving in a separate file as "atlas_landmarks.txt.csv" (Note: If the values are much higher (e.g. > 1000), then this is due to ImageJ representing them in micrometers and not in pixel values. To arrive at the correct pixel values, divide by 25 (in case of a 25 µm/voxel template).
- Again, close the Calc window, then remove the .csv ending from the file.

- If you haven't, move both atlas_landmarks.txt and autofluo_landmarks.txt into the brain folder.

### Cell detection

11 Adapt the process_Ilastik_run_this.py file as follows:

Firstly, comment out the previous action (line 58):

```
#TODO: Specifiy which action you want to take. Uncomment what you want to do
  #Action = 'Preprocess' #only generate scaled-down resamples of expression + autoflu, if you want to manually align to the atlas
```

Then, select (uncomment) one of the three following options:

```
  #Action = 'Find_Cells_With_Ilastik' #find cells with Ilastik. If so, you will need to specify the Ilastik process file below
  #Ilastik_Classifier = "/CloudMap/Data/Classifiers/name_to_your_Ilastik_project.ilp"

  #Action = 'Import_cells_csv' #Import external cell counts from e.g. Arivis (needs to be csv with only x/y/z coordinates, no header. You'll need to specify the file below.
  #Cells_csv = "/CloudMap/Data/cells.csv"

  #Action = 'ClearMap_2_detection' #Choose this if you want to run the standard ClearMap2 cell detection.
```

Default is the first Action = 'Find_Cells_With_Ilastik'. If you choose this, make sure to also adapt the name of your Ilastik classifier to match.

**Note**: make sure you have sufficient amounts of RAM (or Swap under Linux) present. By default the Ilastik segmentation will run on as many cores as possible and consume >200 GB of memory.

11.1

Then, Elastix will be used to transform the points, first for the inter-channel correction alignment, and then for the atlas registration.

```
#%% Cell alignment

  #source = ws.source('cells', postfix='filtered')
  source = ws.source('cells', postfix='filtered')

  def transformation(coordinates):
    #debug
    print ("starting resampling points, coords:
",coordinates[0])

    coordinates = res.resample_points(
                    coordinates, sink=None, orientation=None,

source_shape=io.shape(ws.filename('stitched')),

sink_shape=io.shape(ws.filename('resampled')),
                    );

    #debug
    print ("starting transforming points resamnpled -> auto,
coords: ",coordinates[0])

    coordinates = elx.transform_points(
                    coordinates, sink=None,

transform_directory=ws.filename('resampled_to_auto'),
                    binary=False, indices=False);

    #debug
    print ("starting transforming points auto -> reference,
coords: ",coordinates[0])

    coordinates = elx.transform_points(
                    coordinates, sink=None,

transform_directory=ws.filename('auto_to_reference'),
                    binary=False, indices=False);

    return coordinates;


  coordinates = np.array([source[c] for c in 'xyz']).T;

  coordinates_transformed = transformation(coordinates);
```

Next, the overview tables are generated. This function will generate both a CM2-style table as well as a CM1-style table. If you only need one or the other, block-quote the relevan block.

Note that the intensities-adjusted table is mostly relevant for the CM2 built-in cell segmentation, and is deactivated by default.

```
#%%################################################################
###############
  ### Cell csv generation for external analysis

################################################################
###############

  #%% CSV export

  source = ws.source('cells');
  header = ', '.join([h[0] for h in source.dtype.names]);
  np.savetxt(ws.filename('cells', extension='csv'), source[:],
header=header, delimiter=',', fmt='%s')

  #%% ClearMap 1.0 export

  source = ws.source('cells');

  clearmap1_format = {'points' : ['x', 'y', 'z'],
                      'points_transformed' : ['xt', 'yt', 'zt'],
                      'intensities' : ['source', 'dog',
'background', 'size']}

  for filename, names in clearmap1_format.items():
    sink = ws.filename('cells', postfix=['ClearMap1',
filename]);
    data = np.array([source[name] if name in source.dtype.names
else np.full(source.shape[0], np.nan) for name in names]);
    io.write(sink, data);


  #CM1-style table export
  #TODO: add extra label
  counts =
ano.count_label(label['order'],weights=None,key='order',
hierarchical=True)
  ids_list = ano.get_list('id')
  name_list = ano.get_list('name')

  #additional info
  acronym_list = ano.get_list('acronym')
  order_list = ano.get_list('graph_order')
  parent_list = ano.get_list('parent_structure_id')
  level_list = ano.get_list('level')
  color_list = ano.get_list('color_hex_triplet')
```

```
  table = np.zeros(counts.shape, dtype=[('id','int64'),
('counts','f8'),('name', 'U256'),('acronym', 'U256'),
('order','U256'),('parent_structure_id','U256'),
('level','U256'),('color_hex_triplet','U256')])
  table['id'] = ids_list
  table['counts'] = counts
  table['name'] = name_list

  #additional info
  table['acronym'] = acronym_list
  table['order'] = order_list
  table['parent_structure_id'] = parent_list
  table['level'] = level_list
  table['color_hex_triplet'] = color_list

  #export to csv file
  np.savetxt(io.join(directory,
'Annotated_counts_ClearMap_1.csv'), table, delimiter=';',
fmt='%s',
             header="id;
counts;name;acronym;order;parent_structure_id;level;color_hex_tr
iplet")
```

Subsequently, generate heatmaps. The intensity heatmaps are mostly interesting if you are using the built-in Clearmap2 built-in processing. If using Ilastik (default) or external coordinates, then it is commented out.

```
#%%################################################################
###############
  ### Voxelization - cell density

################################################################
###############

  source = ws.source('cells')

  coordinates = np.array([source[n] for n in
['xt','yt','zt']]).T;
  #intensities = source['source'];

  #%% Unweighted

  voxelization_parameter = dict(
        shape = io.shape(annotation_file),
        dtype = None,
        weights = None,
        method = 'sphere',
        radius = (7,7,7),
        kernel = None,
        processes = None,
        verbose = True
      )

  vox.voxelize(coordinates, sink=ws.filename('density',
postfix='counts'), **voxelization_parameter);
```

#### Atlas alignment with Dockerized Clearmap pipelines

12 Subsequently, run the Docker container for the second time. If everything goes according to plan, this should generate a table for the cells per region. The Docker container should finish after running.

**On Windows**, open power shell, navigate to the folder with the process_Ilastik_run_this.py script and adapt the following code:

```
docker run -it --rm -v C:\path\to\brain:/CloudMap/Data
-v C:\path\to\custom\Ilastik\file:/CloudMap/classifiers
friendlyclearmap:4.2 ;
```

Note that file paths must not end on a trailing backslash.

**On Linux**, open a terminal, then navigate to the folder with the process_Ilastik_run_this.py script and run the following command:

```
docker run -it --rm -v /path/to/brain/:/CloudMap/Data
-v /path/to/custom/Iastik/file/:/CloudMap/classifiers
friendlyclearmap:4.2
```

**On MacOS**, open a terminal (Spotlight -> press Space -> type " Terminal"), then navigate to the folder with the process_Ilastik_run_this.py script and type the folllwing command:

```
docker run -it --rm -v /path/to/brain/:/CloudMap/Data
-v /path/to/custom/Iastik/file/:/CloudMap/classifiers
friendlyclearmap:4.2
```

By default, the container will execute the process_Ilastik_run_this.py script. If you want to run a different script, you can add the option

```
--entrypoint "conda -n cm2 /bin/bash python3
/CloudMap/Data/your_custom_script.py"
```

prior to the friendlyclearmap:4.2 argument.
Note that in this case the script needs to be in the folder that is mapped to /CloudMap/Data with the first -v command (i.e. the folder containing all image data).

**12.1** Hooray! If everything worked, you should have:

- an atlas-aligned set of points, both in .npy form (cells_aligned.npy)
- a region table (Annotated_counts.csv) with cell counts per region.
- A heatmap with cell densities in the same space as your atlas.
