## Supplementary Protocol 3: iDISCO+ protocol for tdTomato for "FriendlyClearMap: An optimized toolkit for mouse brain mapping and analysis"

### tdTomato iDISCO+ Protocol

Moritz Negwer[1]

[1]Radboudumc Nijmegen

M Moritz Negwer

ABSTRACT

This protocol is a supplement to our upcoming publication "FriendlyClearMap: An optimized toolkit for mouse brain mapping and analysis".

In this protocol, we describe how to prepare the samples for light-sheet imaging. Specifically, how to extract the mouse brains and permeabilize, immunostain, and clear them using the iDISCO+ protocol.

**DOI:**
dx.doi.org/10.17504/protocols.io.36wgq77m5vk5/v1

**Protocol Citation:** Moritz Negwer . tdTomato iDISCO+ Protocol. **protocols.io** https://dx.doi.org/10.17504/protocols.io.36wgq77m5vk5/v1

**License:** This is an open access protocol distributed under the terms of the Creative Commons Attribution License, which permits unrestricted use, distribution, and reproduction in any medium, provided the original author and source are credited

**Created:** Feb 06, 2022

**Last Modified:** Feb 13, 2022

**PROTOCOL integer ID:** 57866

#### Sample Isolation

30m

1 **Prepare:**
- 1 ice box
- Isoflurane container
- Sharp scissors or rodent guilloutine for decapitation
- For brain isolation: Sharp small scissors, forceps, Freer elevator (or other flattened, slightly sharpened spatula)
- Scalpel

**Prepare per sample:**
- 50 ml tube with PBS to rinse the sample prior to fixation (cooled on ice)
- 15 ml tube with 4% PFA + 4% Sucrose in PBS, properly labelled (cooled on ice)
- 1.5ml Eppendorf tube for genetic material sampling for re-genotyping

2 Deeply anesthetize the mouse with Isoflurane in a closed container. Use a fume hood to prevent exposure to Isoflurane vapors. Ascertain deep anesthesia by absence of the paw pinch reflex, then quickly decapitate the animal.

3 Collect the head on an inverted petri dish on a box full of ice. Using the small scissors, cut the scalp along the midline from posterior -> anterior.
Subsequently, carefully cut the base of the skull on both sides, from the entry of the spinal column to the ear canals. Carefully cut with the small scissors along the skull's midline posterior -> anterior, taking care not to cut into the brain tissue. Cut until reaching the anterior portion of the olfactory bulbs, then cut laterally in a 45° angle to the outer reaches of the OBs.

4 Using the forceps, grab one of the cranial bones along the midline at the height of the cerebellum and pull laterally. It should break in one clean break, exposing the brain. Repeat for the other side.

Using the scalpel, cut sagitally along the midline to get two brain hemispheres.

Holding the skull over the open PBS washing tube, carefully slide the spatula (or Freer elevator) under the brain, sever the cranial nerves at the bottom and lift the brain hemisphere out of the skull. Repeat for the other hemisphere.

Quickly rinse the brain in PBS to wash off the blood, then transfer to the 15 ml tube with ice-cold 4% PFA in PBS. Fix at 4°C overnight.

Sample a small bit of unfixed tissue (e.g. from the ear) and keep in the 1.5 ml Eppendorf tube for confirmatory genotyping.

The next day, remove the fixative and wash the brains 3x 5 min in PBS. Use immediately or store at 4°C in PBS + 0.005% Sodium Azide until use.

#### iDISCO+ clearing

6d

5 Modified from the original iDISCO+ protocol:

CITATION

Renier N, Adams EL, Kirst C, Wu Z, Azevedo R, Kohl J, Autry AE, Kadiri L, Umadevi Venkataraju K, Zhou Y, Wang VX, Tang CY, Olsen O, Dulac C, Osten P, Tessier-Lavigne M (2016). Mapping of Brain Activity by Automated Volume Analysis of Immediate Early Genes.. Cell.
LINK
https://doi.org/10.1016/j.cell.2016.05.007

Transfer the hemispheres into 5 ml screw-lid or snap-lid Eppendorf tubes.

**Note:** As the solvents used in this protocol dissolve most labelling pens, we recommend to scratch the essential labelling information into the tube (body, not lid) with a sharp implement.

6 Wash 1x 5 min in PBS at RT on a shaker.

Dehydrate the samples by a rising gradient of Methanol / $H_2O$ , 1h each at RT on a shaker. Work in a fume hood when handling Methanol.

20 / 40 / 60 / 80 / 100 % MeOH in water, 1h each at RT on a shaker

6.1 Then repeat 100% MeOH, but start chilling the sample at 4°C (preferrably on a shaker).

Pre-chill the Methanol for the bleaching step as well.

6.2 Overnight, bleach the sample with 5% $H_2O_2$ in MeOH at 4°C (preferably on a shaker).

Freshly prepare this solution using 30% $H_2O_2$. Dilute 1:5 in cold Methanol for a final concentration of 5% $H_2O_2$ .

7 Rehydrate the samples at RT on a shaker with a falling Methanol gradient.

In the meantime, start preparing the other solutions:

**PTx.2** (for 1L)
1L PBS
20 g Triton-X 100 for a final concentration of 0.2% (W/V)
(drip from a cut-off bulb pipette into the flask while weighing it on a scale, as Triton is very viscous an therefore difficult to pipet reliably).

**Permeabilization solution** (for 100 ml)
50 ml PTx.2
20 ml DMSO
2.3g Glycine
Sodium Azide (from stock solution) for a final concentration of 0.005% (w/v)
Stir, then fill to 100 ml with PTx.2

**Blocking Buffer** (for 100 ml)
50 ml PTx.2
10 ml DMSO
6 ml Neutral Donkey Serum
Sodium Azide (from stock solution) for a final concentration of 0.005% (w/v)
Stir, then fill to 100 ml with PTx.2

**PTwH** (for 100 ml)
50 ml PTx.2
Heparin from stock for a final concentration of 0.01 g/L
Sodium Azide (from stock solution) for a final concentration of 0.005% (w/v)
Stir, then fill to 100 ml with PTx.2

7.1 Rehydrate the sample by a decreasing Methanol / $H_2O$ gradient

(Note to start with 80% as the $H_2O_2$/MeOH solution is essentially already a ~83% MeOH solution, with the rest being 5% $H_2O_2$ and ~12% water that came with the $H_2O_2$ ).

80 / 60 / 40 / 20 % Methanol in H2O, 1h each at RT on a shaker

7.2 Wash in PTx.2, 30 min @ RT on a shaker

7.3 Repeat: Wash in PTx.2, 30 min @ RT on a shaker

7.4 Permeabilize in permeabilization solution (see above) for 2 days @ 37°C in an incubator with rotor.

7.5 Block the samples in blocking solution (see above) for 2 days @ 37°C in an incubator with rotor.

#### iDISCO+ antibody incubation

2w 1d

8 Prepare **primary antibody solution**, 2 ml per sample:

PTwH (see above)
5% DMSO
3% Neutral Donkey Serum
Primary antibody: Rabbit anti-RFP (600-401-379), diluted 1:2000

Perform this step in 2 ml Eppendorf screw-cap or tight-seal flip-cap tubes. Label the tubes by scratching with a sharp implement to prevent the label being washed off. Keep the 5ml tubes for the later dehydration step.

Transfer the samples to the tubes, either by carefully pouring them over, or by careful use of a spatula.

Fill with primary antibody incubation solution (to the top to avoid air bubbles as much as possible). Seal the outsides with a thin strip of Parafilm.

Incubate for 6-7 days @ 37°C in an incubator with rotor.

9 Wash 4-5x with PTwH, each for 1h @ RT on a rotor.

Wash overnight with PTwH @ RT on a rotor.

10 Prepare **secondary antibody solution**, 2 ml per sample:
PTwH (see above)
5% DMSO
3% Neutral Donkey Serum
Secondary antibody: Goat anti-Rabbit Alexa 568 or 647 (Invitrogen A-11036 / A-21245), dilution 1:500

Incubate for 5-7 days @ 37°C in an incubator with rotor.

#### iDISCO+ clearing

3d

11 Transfer back to the 5ml Eppendorf tubes. Keep the tubes wrapped in aluminium foil or place under a box while incubating to prevent light exposure. Work in a fume hood when handling Methanol.

Wash 4-5x with PTwH, each for 1h @ RT on a rotor.

12 Dehydrate the samples by a rising gradient of $H_2O$ / Methanol:

20 / 40 / 60 / 80 / 100 / 100 % Methanol in H2O, 1h each at RT on a shaker.
Leave last 100% Methanol overnight

13 Delipidate the sample in a freshly prepared 66% DCM / 33% Methanol solution, 3h @ RT on a shaker, in a fume hood.

**Note** that DCM is a strong solvent that will dissolve many plastics (most notably Polystyrene). The Eppendorf tubes should be made from Polypropylene to withstand DCM. **Avoid pipetting DCM**, and instead only pour it from and into glass containers. Pre-mix the DCM/MeOH mixture in a measurement cylinder and pour into the 5ml Eppendorf tubes. To empty those, carefully pour out the fluid while keeping the dehydrated sample in place with forceps.

14 Incubate in 100% DCM on a shaker in a fume hood for 15 minutes to wash out the Methanol.

15 To clear, incubate the samples in 100% Dibenzyl Ether (DBE) at RT in the dark, no shaking required. Make sure to fill the tubes as much as possible to prevent oxidation of the DBE. The sample should become completely transparent within 24 hours. Store at RT in the dark, with tightly locked lids, until imaging with a suitable light-sheet microscope.
