## Supplementary material for "FriendlyClearMap: An optimized toolkit for mouse brain mapping and analysis": Supllementary Protocol 4: Visualize Data with BrainRender

Moritz Negwer[1]

[1]Radboudumc Nijmegen

M Moritz Negwer

ABSTRACT

Supplemementary protocol for our submitted paper "FriendlyClearMap: An optimized toolkit for mouse brain mapping and analysis".
This protocol describes how to pre-process cell locations in mouse brains (in different reference atlases) and visualize them using BrainRender (Claudi et al,. eLife 2021).

**DOI:**
dx.doi.org/10.17504/protocols.io.dm6gpbdwdlzp/v1

**Protocol Citation:** Moritz Negwer . Visualize Data with Brainrender. **protocols.io** https://dx.doi.org/10.17504/protocols.io.dm6gpbdwdlzp/v1

**License:** This is an open access protocol distributed under the terms of the Creative Commons Attribution License, which permits unrestricted use, distribution, and reproduction in any medium, provided the original author and source are credited

**Created:** Apr 11, 2022

**Last Modified:** May 22, 2022

**PROTOCOL integer ID:** 60561

### Setup

1 After you have generated your cell table with Clearmap 1 or 2, do the following:
(**Note** this has only been tested on Ubuntu Linux 20.04 and 21.10)

1. If you haven't, install Conda and Spyder

https://docs.anaconda.com/anaconda/install/linux/

```
sudo apt-get install libgl1-mesa-glx libegl1-mesa libxrandr2
libxrandr2 libxss1 libxcursor1 libxcomposite1 libasound2 libxi6
libxtst6
```

Create a new conda environment:

```
conda create --name brainrender
conda activate brainrender
conda update --all
```

(this might take several minutes)

Install Spyder with all dependendcies via conda:

```
conda install spyder
```

2. Install Brainrender from here: https://github.com/brainglobe/brainrender

CITATION

Claudi F, Tyson AL, Petrucco L, Margrie TW, Portugues R, Branco T (2021). Visualizing anatomically registered data with brainrender.. eLife.
LINK
https://doi.org/pii:e65751.10.7554/eLife.65751

3. Install Elastix (e.g. on Ubuntu / Debian Linux):

```
sudo apt-get update && sudo apt-get install elastix
```

4. Download the brainrender script from github:
https://github.com/MoritzNegwer/FriendlyClearMap-scripts/tree/main/Young_Mouse_Atlas_Alignments

### Transform your data to CCF3

2 If you used a different atlas from CCF3 (e.g. when using Clearmap 1 or one of the young mouse atlases), you need to transform it to CCF3 space.

We have pre-computed a number of transforms for CCF2 (=CM1) and the young mouse atlases from the Kim lab, using Elastix. Download them from github here: XXX_needs_to_be_added

### Visualize neurons in the brain with Brainrender

3 Adapt the visualize_brainrender script as follows:

1. Adapt the transformix_path variable to your own. Check in the command line:

```
whereis transformix
```

and take the first entry, e.g. "/usr/bin/transformix"

2. Make a list of your mice in the target_mice list, pointing to the cellls_transformed_to_Atlas_aligned.npy in each folder:

```
target_mice =
[["path_to_my_mouse_1/cells_transformed_to_Atlas_aligned.npy",
myterial.black],
["path_to_my_mouse_2/cells_transformed_to_Atlas_aligned.npy",
myterial.black]
]
```

The colors are defined via the myterial package. Refer here for the color palette: [https://github.com/FedeClaudi/myterial](https://github.com/FedeClaudi/myterial)

3. Define the target_folder. If it does not exist yet, it will be created when running the script. **Note**: If it does contain images already, those will be overwritten during the process!

```
target_folders = ["this_is_where_I_want_my_images_to_end_up"]
```

4. Define the transformation you want to use to get your cells to CCF3 coordinate space (refer to step 2 above).

for Clearmap 1:

```
target_transformation =
["downloaded_transformation_location/elastix_files/Par_rotate90dega
roundX_Clearmap1.txt"]
```

for young mouse atlases, a series is needed. In this case adapt the location and the atlas in question (here: P21):

```
target_transformation =
["downloaded_transformation_location/elastix_files/Par_rotate90dega
roundX_CCF3_nrrd_directions.txt",

"downloaded_transformation_location/Kim_ref_P21_v2_brain_CCF3/Trans
formParameters.0.txt",

"downloaded_transformation_location/Kim_ref_P21_v2_brain_CCF3/Trans
formParameters.1.txt"]
```

5. Adapt the regions you want to make specific visualizations for (optional). If you do, the script will create one image for each region. An easy way to check for the regions and the acronyms is via the 3D Atlas Viewer of the Allen Brain Institute.
We used the following list:

```
target_regions = ["grey", #All brain
                  "HIP", #Hippocampal region
                  "Isocortex", #Cortex proper
                  "CNU", #Striatum + GP
                  "TH", #Thalamus
                  "CTXsp", #cortical subplate
                  "BLA", #Basolateral Amygdala
                  "MB", #Superior / inferior colliculus
                  "SSp-bfd", #S1 Barrel Field
                  "SS", #S1
                  "AUD", #A1
                  "VIS" #V1
                  ]
```

6. Adapt the camera position. The default camera can be found:
brainrender_folder/brainrender/cameras.py
If you want to define your own, run the following code in spyder (after adapting the brainrender_folder):

```
import sys
sys.path.append('brainrender_folder/')

# Import variables
from brainrender import * # <- these can be changed to personalize
the look of your renders

# Import brainrender classes and useful functions
from brainrender import Scene
scene = Scene (title=None,screenshots_folder=None,inset=None)
scene.render()
```

Subsequently, a window with an empty mouse brain silhouette should open.

**Note**: If it does not appear, brainrender´s default setting could be set to offscreen rendering. Change the last line of brainrender_folder/brainrender/settings.py from OFFSCREEN = False to OFFSCREEN = True

Move the brain around until you like the angle, then type "C" to export the current camera parameters to your spyder console. It should looke like this:

```
Camera parameters:
    {
     'pos': (7572, -1308, -49730),
     'viewup': (0, -1, 0),
     'clippingRange': (32602, 59249),
   }
Additional, (optional) parameters:
     'focalPoint': (6888, 3571, -5717),
     'distance': 44288,
```

Reformat them to this format to use in the script:

```
techpaper_cam_01 = {
    "pos": (2093, 2345, -49727),
    "viewup": (0, -1, 0),
    "clippingRange": (33881, 52334),
    "focalPoint": (6888, 3571, -5717),
    "distance": 44288,
    }
```

Define the camera to be used in the script as follows (boldened part):

```
for mouse in target_mice:
    #transform to CCF3
    cells = transform_points(mouse[0],target_transformation)
    #multiply to get to µm value(assuming atlas has 25µm voxels)
    cells = cells*25
    #flip axes because of CCF2. Comment out if using any other
atlas
    cells = flip_CCF2_axes(cells)
    #add to the cells_all array (uncomment if you want to create a
group average)
    #cells = np.concatenate((cells,cells),axis=0)

    for region in target_regions:
        render_screenshot (target_folder, cells, mouse[0],
mouse[1], region,techpaper_cam_01,render_hemisphere=True)
```

7. If you want to render only one hemisphere, set render_hemisphere to True (boldened part):

```
render_screenshot (target_folder, cells, mouse[0], mouse[1],
region,techpaper_cam_01,render_hemisphere=True)
```

4 Run the script. If all goes according to plan, you should get a series of images from each mouse

brain, one per brain region defined under substep 5.

If you suspect that the cell coordinates are wrong, run the following code in the spyder command line:

```
cells = transform_points(target_mice[0][0],target_transformation)
render_screenshot (target_folder, cells, mouse[0], mouse[1],
"",techpaper_cam_01)
```

This should render the brain without any regions cropping and reveal the full extent of your cells. If for example all cells are rendered outside of the CCF3 mouse brain, then you will need to adapt the transformation or the cells, e.g. refer to the flip_CCF2_axes function.

### Render cell densities

5 If you want to render the points densities (such as in Fig 2 c / f), do the following:

For densities for individual brains, simply adapt the render_screenshot call with render_densities:

```
render_screenshot (target_folder, cells, mouse[0], mouse[1],
region,techpaper_cam_01,render_hemisphere=True,render_densities=Tru
e)
```

6 If you want to sum all densities across a group, uncomment the following line and move the next for loop out of the for mouse [...] loop:

```
for mouse in target_mice:
    #transform to CCF3
    cells = transform_points(mouse[0],target_transformation)
    #multiply to get to µm value(assuming atlas has 25µm voxels)
    cells = cells*25
    #flip axes because of CCF2. Comment out if using any other
atlas
    cells = flip_CCF2_axes(cells)
    #add to the cells_all array (uncomment if you want to create a
group average)
    cells = np.concatenate((cells,cells),axis=0)

for region in target_regions:
    render_screenshot (target_folder, cells, name_path, mouse[1],
region,techpaper_cam_01,render_hemisphere=True,render_densities=Tru
e)
```

Then, adapt the name_path. The end_folder is the name the resulting average density will have. (Note that this folder does not need to exist, this is just a workaround to create a name).

```
name_path = "/some_random_path/names/PV_P56_WT_avg/"
```

7 By default this script renders rather coarse maps, suitable to assess changes across e.g. large cortical regions. If you want more fine-grained heatmaps, you can set the "radius" parameter in the render_screenshot function to a lower value (i.e. 250).

```
scene.add(PointsDensity(cells,colormap="twilight",radius=250))
```

If you want the heatmap object itself to have a higher resolution (and have lots of RAM and time), you can additionally define the "dims" parameter for the PointsDensity function:

```
scene.add(PointsDensity(cells,dims=
(200,200,200),colormap="twilight",radius=750))
```

The default is (40,40,40), which works as a reasonable compromise between computing speed and display accuracy.

8 Run the script. Depending on the number of cells, computing the density might take some time (>30 min on a Ryzen 4600H with 64GB RAM).

**Note:** The color scales for the heatmaps are set to the min/max of the dataset and are not comparable between different datasets. The underlying data can be retrieved by digging deep into the vtk code underlying brainrender, but is not easily accessible (and then needs to be re-rendered).
